# Refining RDoC Using Individual-Level Task fMRI Factor Models Reveals Reproducible and Clinically Relevant Brain-Wide Motifs

**DOI:** 10.1101/2025.10.13.682124

**Authors:** S.K.L. Quah, S. Pirzada, S. Madsen, B. Jo, L.Q. Uddin, J.A. Mumford, D.M. Barch, I.H. Gotlib, D.A. Fair, R.A. Poldrack, M. Saggar

**Affiliations:** Department of Psychiatry & Behavioral Sciences, Stanford University, Stanford, CA, USA; Department of Psychiatry and Biobehavioral Sciences, University of California, Los Angeles, CA, USA; Department of Psychology, Stanford University, Stanford, CA, USA; Departments of Psychological & Brain Sciences, Psychiatry, and Radiology, Washington University in St. Louis, St Louis, MO, USA; Department of Pediatrics, University of Minnesota Medical School, Minneapolis, MN, USA; Wu Tsai Neurosciences Institute, Stanford University, Stanford, CA, USA

**Author notes:** Corresponding author: Manish Saggar. These authors contributed equally (S.K.L. Quah, S. Pirzada).

## Abstract

The Research Domain Criteria (RDoC) framework was introduced to guide psychiatric research using biologically grounded, dimensional constructs of mental function. However, its hierarchical domain structure remains largely unvalidated against individual-level brain and behavioral data. Building on prior group-level work, we applied a multi-stage validation framework to Human Connectome Project (HCP) task-fMRI data to test whether individual-level, data-driven models more accurately capture the organization of brain activity and behavior than RDoC-based models. Using confirmatory factor analysis in two independent cohorts, we found that data-driven bifactor models consistently outperformed RDoC-based models across multiple fit indices. The general factor derived from these models revealed a reproducible, low-dimensional axis spanning visual–attentional to default mode–auditory systems, aligning with canonical macroscale cortical gradients. Community detection further identified reproducible spatial motifs whose centroids corresponded to interpretable functional systems and whose alignment predicted individual performance on working memory and relational reasoning tasks. To assess whether these findings extended beyond neural data, we analyzed behavioral measures in HCP and in an independent transdiagnostic dataset (LA5c). In both datasets, data-driven behavioral models outperformed RDoC-based models, although the relative support for bifactor versus specific factor structure differed by dataset. Extending the neural analyses to LA5c, which included healthy controls and individuals with ADHD, bipolar disorder, and schizophrenia, showed that data-driven bifactor models generalized across diagnostic groups and that alignment with data-driven community centroids related to symptom severity, whereas RDoC-based representations showed weaker or no associations. Finally, topological analysis of task-evoked brain activity revealed that data-driven representations better captured the global organization of brain states than RDoC domains. Together, these findings demonstrate that individual-level, empirically derived models provide a more accurate, generalizable, and behaviorally relevant account of brain organization than the current RDoC framework. By integrating neural, behavioral, and clinical validation, this work advances precision neuroscience and supports the empirical refinement of dimensional psychiatric frameworks.

## 1. Introduction

The Research Domain Criteria (RDoC) framework was developed to advance precision psychiatry by grounding mental health constructs in neurobiological systems^1,2^. Rather than relying on categorical diagnoses, RDoC proposes that mental disorders arise from variation in core psychological functions and their underlying neural circuitry^3,4^. Over the past decade, this framework has provided a widely used ontology for organizing psychiatric research across multiple levels of analysis, from genes to behavior.

Despite its influence, a central assumption of RDoC that its predefined domains map onto distinct neural systems remains only partially validated. Emerging evidence suggests that brain activation patterns often cut across RDoC domains^5^. For example, meta-analytic work has shown substantial overlap in neural circuitry across tasks nominally assigned to different domains^6^, while domains such as cognitive systems may encompass heterogeneous processes engaging diverse brain networks^5,7^. More broadly, the RDoC framework was intended as a heuristic starting point rather than a fixed taxonomy^8^, highlighting the need for empirical evaluation of its structure.

To address these limitations, recent work has turned to data-driven approaches that derive latent dimensions directly from neuroimaging and behavioral data^5,7,9^. These methods offer a flexible alternative to predefined domain boundaries and have shown that empirically derived factors can better capture the structure of brain activity than RDoC-based models^7^. However, most such approaches operate at the group level, leaving open whether these latent dimensions reflect subject-specific organization or generalize across individuals, datasets, and levels of analysis.

In parallel, several alternative frameworks have sought to organize psychopathology using empirically derived dimensions. For example, the Hierarchical Taxonomy of Psychopathology (HiTOP)^10^ proposes a data-driven, transdiagnostic structure based on patterns of symptom covariance, emphasizing continuous dimensions over categorical diagnoses. Related efforts in computational psychiatry and network-based approaches have similarly aimed to derive latent structure directly from behavioral and neural data. While these frameworks differ in scope and methodology, they share a common goal of grounding psychiatric constructs in empirical structure rather than predefined categories. However, it remains unclear how these alternative dimensional frameworks relate to the organization of brain activity, particularly at the level of individual neural patterns.

A key unresolved question is therefore whether the organization of brain activity at the individual level aligns with existing dimensional frameworks, such as RDoC or other empirically derived models, or instead reflects alternative latent structure not captured by current ontologies. Addressing this question is complicated by the nature of task-fMRI data, which primarily captures state-dependent neural responses, whereas psychiatric constructs are typically conceptualized as trait-like dimensions. As a result, the correspondence between task-evoked brain activity and dimensional models of psychopathology cannot be assumed and must be evaluated empirically.

Here, we test whether data-driven factor models derived from individual-level task-fMRI data provide a better account of functional brain organization than RDoC-based models (Fig. 1). Using task-fMRI data from the Human Connectome Project (HCP), we compare the fit and generalizability of data-driven and RDoC-based models across two independent cohorts, estimating models separately within each individual to capture subject-specific covariance structure. To evaluate whether findings extend beyond neural data, we additionally assess the structure of behavioral measures, enabling comparison of data-driven and RDoC-based models across both brain and behavior.

**Figure 1.**
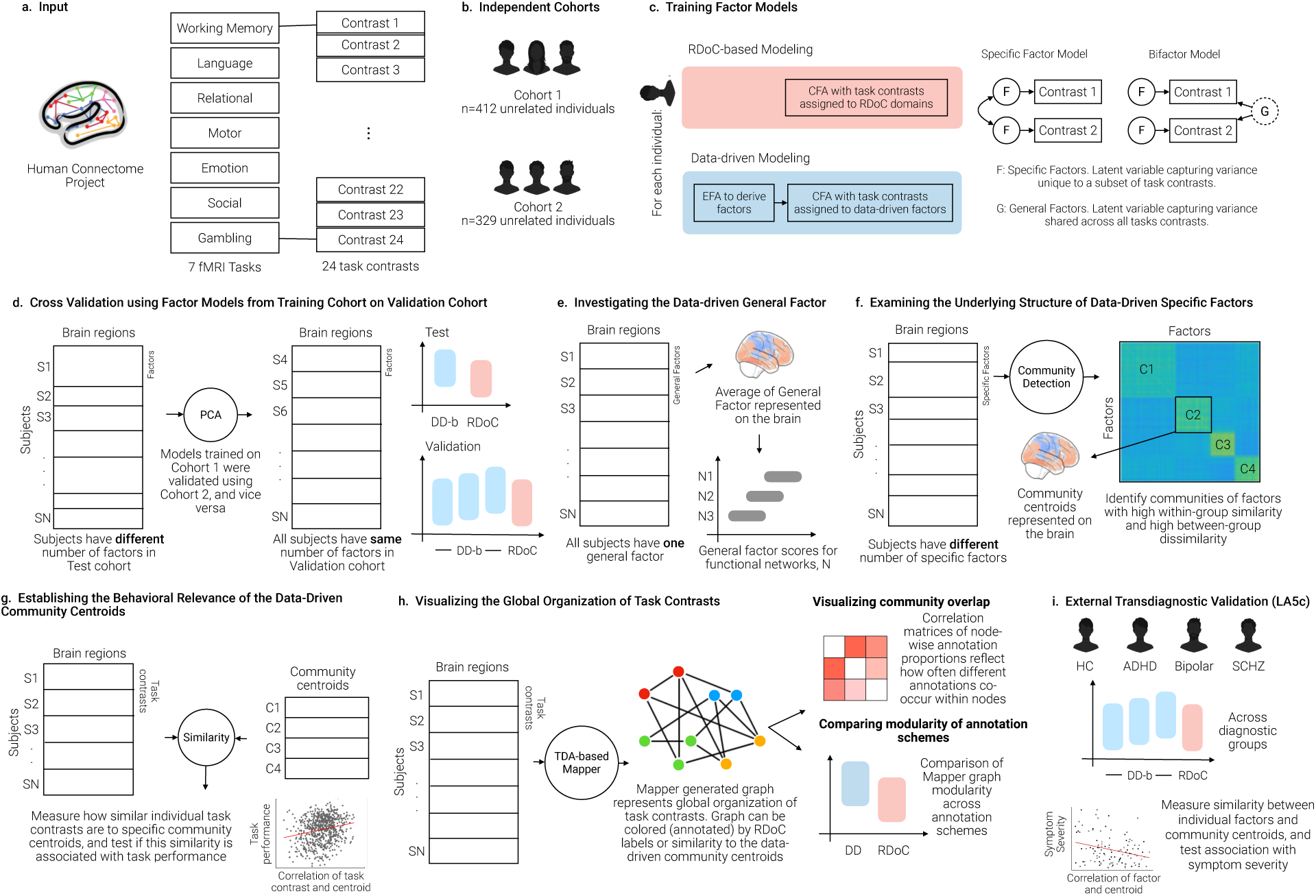
Overview of approach. Using the Human Connectome Project (HCP) task-based fMRI data, we examined whether data-driven factor models derived from individual-level task-fMRI data outperform RDoC-based models in explaining functional brain organization. (a) Preprocessed and parcellated contrast maps from seven HCP tasks were used as input. (b) Data from 741 unrelated subjects were split into two cohorts (Cohort 1: N = 412, and Cohort 2: N = 329). (c) Four CFA model types were trained: RDoC-specific, RDoC-bifactor, data-driven specific, and data-driven bifactor. (d) Cross-validation across the two cohorts was conducted such that the models trained on one cohort were validated on held-out data from another cohort. Principal components analysis (PCA) was applied to data-driven specific factors from the data-driven bifactor model to derive the same number of factors across all individuals (detailed in Methods). (e) Mean general factor scores from the data-driven bifactor models were mapped onto the brain. Reproducibility across cohorts, correspondence with resting-state gradients^12^, and network-wise distributions were evaluated. (f) To examine the underlying structure of the data-driven specific factors, an unsupervised community detection algorithm was used to derive communities. (g) Associations between similarity of individual task activation maps to specific community centroids and behavioral performance were estimated with linear mixed-effects models. Similarity was entered as a fixed effect, with Parental ID included as a random intercept to control for genetic relatedness. (h) We used the TDA-based Mapper approach to visualize the global organization of task contrasts and examined whether RDoC or data-driven groupings (annotation) better capture this structure. (i) Finally, to assess external generalizability and clinical relevance, the framework was extended to an independent transdiagnostic dataset (LA5c), including healthy controls and individuals with ADHD, bipolar disorder, and schizophrenia. Model comparisons were performed across diagnostic groups, and associations between similarity to data-driven community centroids and symptom severity were tested.

To further examine generalizability and clinical relevance, we extend our analyses to an independent transdiagnostic dataset (LA5c^11^), including healthy controls as well as individuals with ADHD, bipolar disorder, and schizophrenia. This allows us to test whether the structure identified in HCP replicates across datasets, populations, and diagnostic groups, and whether data-driven representations capture clinically meaningful variation.

To interpret the resulting latent dimensions, we examine their spatial organization and relationship to established macroscale features of brain organization. Specifically, we assess whether a general factor captures a low-dimensional axis aligned with intrinsic cortical gradients^12^, whether data-driven representations reveal reproducible functional motifs associated with behavior, and whether they better capture the global topological structure of task-evoked brain activity^13,14^.

Together, this multi-level validation framework enables us to test whether the organization of brain activity and behavior aligns with predefined cognitive ontologies or is better described by empirically-derived dimensions^15,16^. By integrating individual-level modeling, complementary group-level behavioral validation, and cross-dataset replication, this work provides a foundation for refining dimensional frameworks of mental function, including RDoC.

## 2. Results

We analyzed task-based fMRI data from the Human Connectome Project (HCP) in 741 unrelated healthy adults, divided into two independent cohorts. Each participant completed seven tasks spanning five core RDoC domains, and whole-brain activation maps were summarized using a standardized brain parcellation.

Sections 2.1–2.6 focus on HCP analyses. We first compared data-driven and RDoC-based models in their ability to explain individual-level patterns of task-evoked brain activity. We then examined the structure of the data-driven models, focusing on the spatial organization of the general factor and its relationship to established cortical gradients. Next, we identified reproducible functional motifs across individuals using community detection and tested their behavioral relevance.

To assess whether these findings extend beyond neural data within HCP, we evaluated HCP behavioral measures at the group level. We then used topological data analysis to characterize the global organization of HCP brain activity and to compare how well data-driven versus RDoC-based representations capture this structure. Finally, in Section 2.7, we tested generalizability and clinical relevance in an independent transdiagnostic dataset, LA5c, including individuals with ADHD, bipolar disorder, and schizophrenia.

### 2.1 Data-driven bifactor models outperform RDoC models in both training and validation cohorts

To evaluate whether individual-level data-driven models better capture the structure of task-evoked brain activity than RDoC-based models, we applied CFA to each subject’s activation maps, treating contrast maps as variables and brain parcels as observations. Data-driven models were constructed using EFA followed by bifactor CFA, while RDoC models used fixed domain assignments. Model fit was assessed using four standard indices: AIC, RMSEA, CFI, and TLI.

In Cohort 1, data-driven bifactor models demonstrated significantly superior model fit compared with RDoC-based models across all evaluated indices. For RMSEA and AIC, data-driven models showed markedly better fit (RMSEA: *M* = .159, *SD* = .014 vs. .207, *SD* = .012; *t*(411) = –75.10, *p* < .001, *d* = –3.70; AIC: *M* = 16,275.1, *SD* = 1040.3 vs. 17,898.8, *SD* = 1141.6; *t*(411) = – 79.05, *p* < .001, d = –3.89) (Figure 2). Similar advantages were observed for the CFI (Data-driven: *M* = .789, *SD* = .032; RDoC: *M* = .613, *SD* = .060; *t*(411) = 66.66, *p* < .001; *d* = 3.28) and TLI (Data-driven: *M* = .743, *SD* = .039; RDoC: *M* = .562, *SD* = .068; *t*(411) = 60.54, *p* < .001; *d* = 2.98) (Supplementary Figure 1).

**Figure 2.**
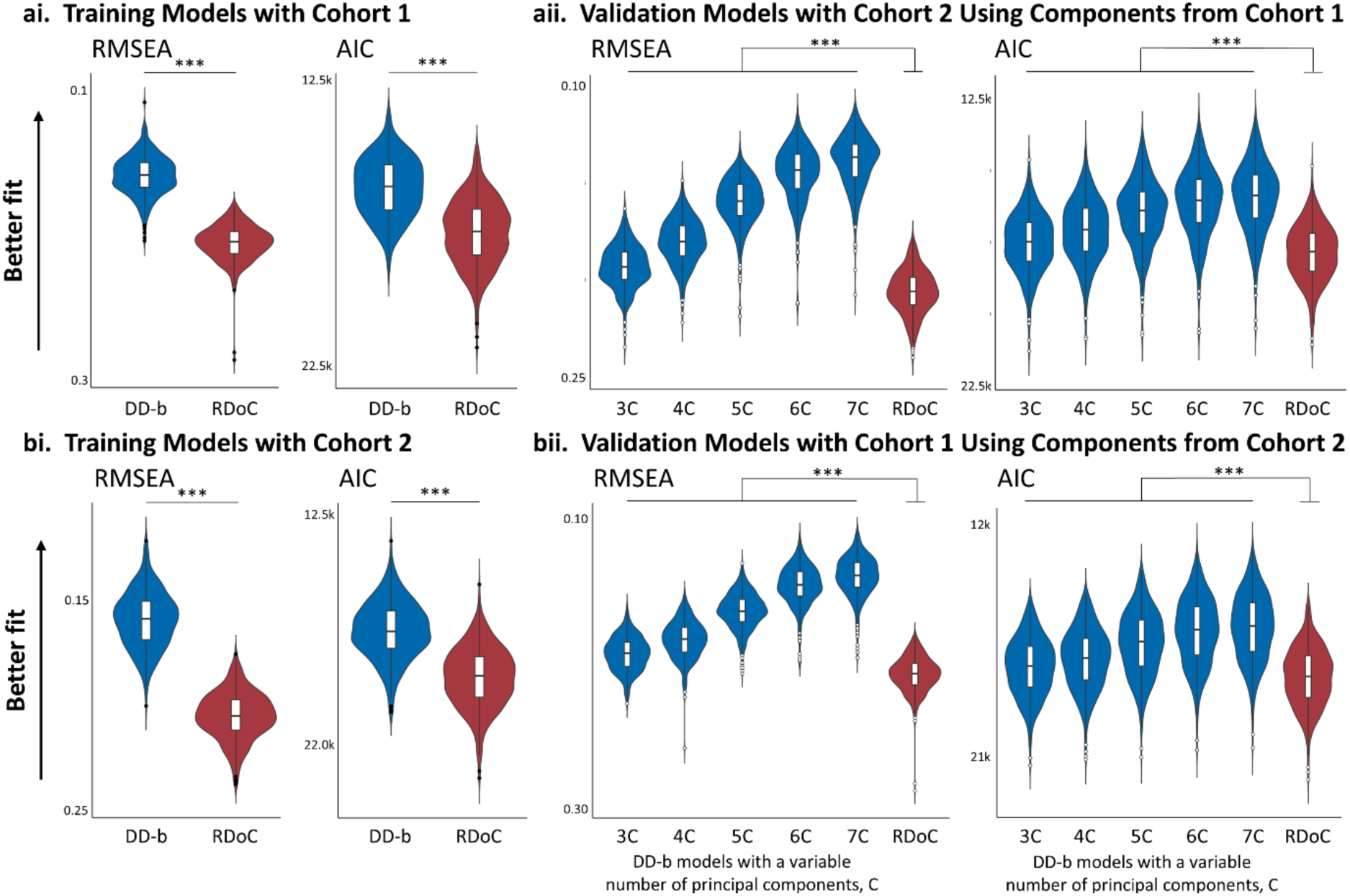
Data-driven bifactor models consistently showed superior model fit compared to RDoC-based models across training and validation comparisons. Root Mean Square Error of Approximation (RMSEA) and Akaike Information Criterion (AIC) were used to assess model performance. For both fit indices, lower values indicate better model fit; y-axes are inverted to display better-fitting models higher on the plot. (ai, bi) Within-cohort model fit comparison for data-driven bifactor (DD-b) and RDoC models trained using Cohort 1 and Cohort 2, respectively. (aii) Models derived from Cohort 1 and validated in Cohort 2 using principal component-based projection. (bii) Models derived from Cohort 2 and validated in Cohort 1. For validation, data-driven bifactor models were tested with 3 to 7 principal components (3C–7C). Violin plots show distributions of fit indices across individuals. Data-driven bifactor models consistently outperformed RDoC-based models across all comparisons. \*\*\**p* < .001.

Swapping the training and validation cohorts yielded the same pattern: data-driven bifactor models consistently outperformed RDoC-based models across all indices. This replication supports the robustness of the data-driven bifactor solution across independent datasets.

To test generalizability more directly, we projected principal components from the data-driven bifactor model in Cohort 1 onto Cohort 2 and repeated the procedure in reverse. These cross-cohort validations focused on comparing the data-driven bifactor model with the canonical RDoC-specific model, allowing us to isolate the contribution of the general factor. In both directions and across all tested component numbers, the data-driven bifactor model outperformed the RDoC model on all fit indices (*p* < .001; Supplementary Tables 1 & 2).

To formally compare all four model types—(i) RDoC-specific, (ii) RDoC bifactor, (iii) data-driven specific, and (iv) data-driven bifactor—we conducted repeated-measures ANOVAs. Mauchly’s test revealed violations of sphericity for all fit indices (*p* < .001), so Greenhouse–Geisser corrections were applied. Post hoc Tukey-adjusted comparisons confirmed that the data-driven bifactor model consistently yielded the best fit (all \*\**p* < .001). Full results, including detailed statistics and pairwise comparisons, are reported in Supplementary Figure 2 and Supplementary Table 3.

Because bifactor models can sometimes overfit correlated-factor alternatives, we next quantified bifactor quality indices for the HCP data-driven bifactor solution (Supplementary Table 4), following recommendations for adjudicating among alternative structural models and for evaluating bifactor solutions^17,18^ Across both cohorts, the general factor was strong and highly replicable, with high omega total, omega hierarchical, factor determinacy, and construct replicability. Percentage of uncontaminated correlations and explained common variance indicated a substantial, but not strictly unidimensional, structure. Specific factors were recoverable, although they accounted for comparatively modest unique variance. Together, these indices indicate that the data-driven bifactor solution is psychometrically well-posed, with a strong general factor and interpretable, albeit weaker, specific factors.

To assess the sensitivity of model comparisons to RDoC domain assignments, we performed a permutation analysis in which contrast-to-domain labels were randomly reassigned to generate a null distribution of model fit (Supplementary Figure 3).

Across both cohorts and all fit indices (AIC, RMSEA, CFI, TLI), the prespecified contrast-to-domain assignment consistently fell within the top 0.3–0.4% of the permutation distribution (Supplementary Table 5), corresponding to approximately the 99.6th–99.7th percentile. These results indicate that model fit is strongly dependent on assignment structure and that the observed mapping performs substantially better than arbitrary alternatives.

Together, these findings demonstrate that data-driven bifactor models provide a significantly better and more generalizable account of task-evoked brain activity than expert-defined RDoC models.

### 2.2 Data-driven General Factor Reflects a Gradient from Visual–Attentional to Auditory–Default Mode Networks

We next examined the spatial topography of the general factor recovered from the data-driven bifactor models in both cohorts. Whole-brain general factor maps showed highly reproducible spatial patterns, with prominent positive loadings in visual and parietal cortices and lower or negative loadings in auditory, somatomotor, and default mode regions (Figure 3a).

**Figure 3.**
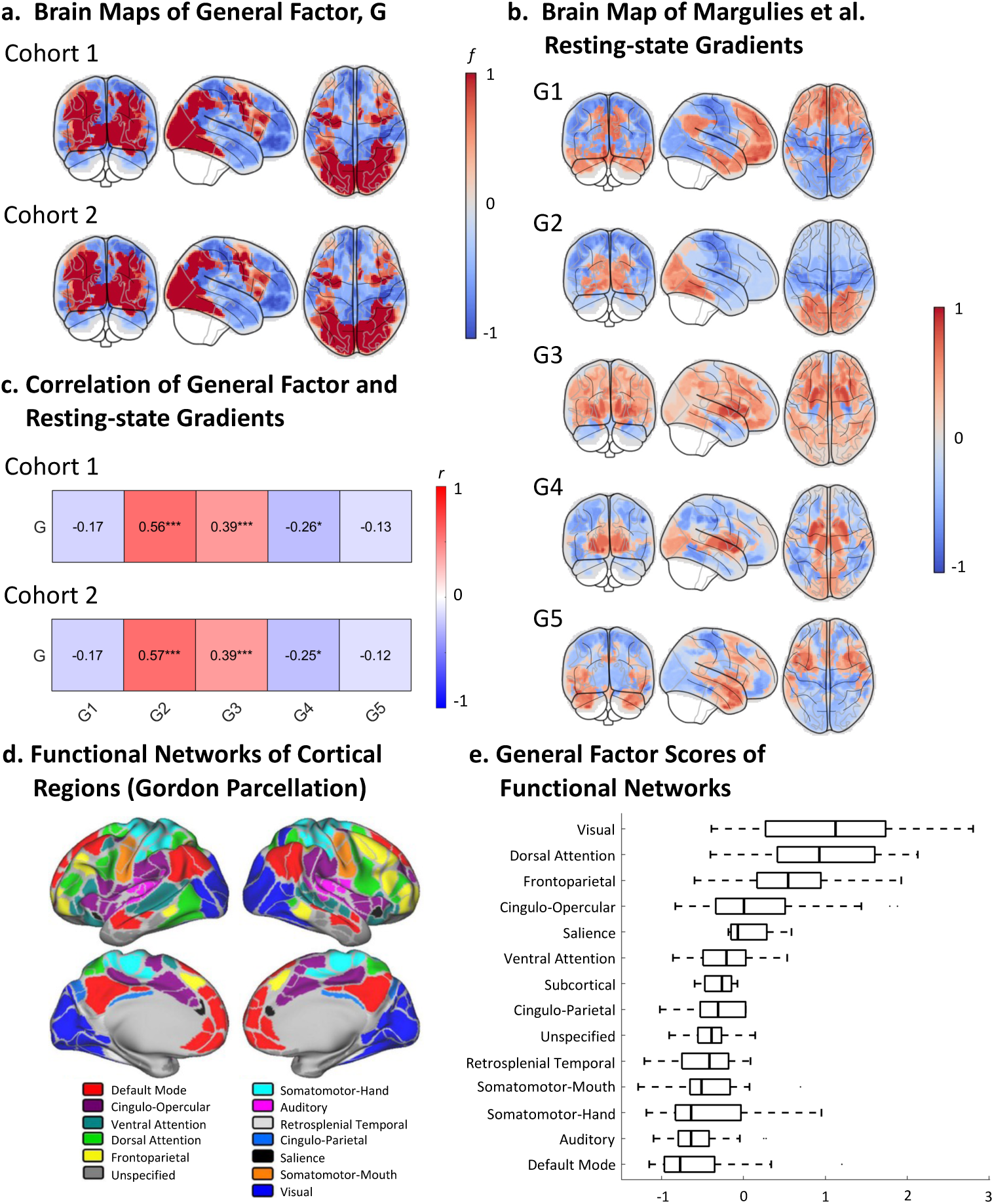
The general factor shows reproducible topography, aligns with resting-state gradients, and maps onto a visual–dorsal attention to auditory–default mode axis. (a) Whole-brain spatial maps of the general factor from the data-driven bifactor models in both cohorts, with reproducible organization across cohorts. (b) Resting-state gradients 1–5 from Margulies et al.^12^ are shown for comparison. (c) Correlations between general factor maps and the resting-state gradients. The general factor was most strongly associated with gradient 2 in both cohorts, followed by a moderate association with gradient 3. Weaker negative correlations were observed with gradient 4. P-values were corrected for spatial autocorrelation. (d) Functional networks of cortical regions based on the Gordon parcellation. Figure adapted from Vazquez-Trejo et al.^21^. (e) Distribution of general factor scores across regions of different functional networks. Networks have been ordered by their median factor scores. Activation follows a gradient from the highest positive values in visual and dorsal attention networks to the lowest/most negative values in somatomotor, auditory, and default mode networks. \**p* < .05, \*\*\**p* < .001.

This pattern captures a latent dimension of shared variance across task contrasts. To assess its relationship to established cortical organization, we compared the general factor maps with the first five resting-state gradients reported by Margulies et al.^12^ (Figure 3b), which describe dominant modes of functional connectivity variation.

In both cohorts, the general factor showed its strongest positive association with Gradient 2 and a moderate positive association with Gradient 3, with weaker negative associations with Gradient 4 and no significant association with Gradients 1 or 5 (Figure 3c; adjusted p-values in Supplementary Table 6). This pattern indicates that the general factor aligns with a visual–attentional axis spanning sensory processing and task-positive control systems, rather than with the principal unimodal–transmodal gradient.

To further contextualize this dimension, we examined its distribution across the Gordon network parcellation^19^. The general factor exhibited a graded organization, with highest scores in visual and dorsal attention networks and lower or negative values in default mode, auditory, and somatomotor networks (Figure 3d–e).

Together, these results indicate that the general factor reflects a reproducible, low-dimensional axis of brain organization that aligns with established macroscale connectivity gradients. This suggests that task-evoked brain activity is structured along intrinsic cortical hierarchies, providing a link between task-driven responses and large-scale intrinsic functional organization.

To further test whether the HCP general factor could be explained primarily by canonical task-positive organization, we compared it with independent Neurosynth^20^ meta-analytic maps of task-related and default mode organization (Supplementary Figure 4; Supplementary Table 7).

Direct correspondence among general factor maps is summarized in Supplementary Table 8. Across both HCP cohorts, the general factor showed only partial correspondence with these external maps. Using the most direct comparison, namely the task-positive association and default mode association maps, the full model explained 6.9–7.2% of the variance in the HCP general factor, leaving the majority of variance unexplained. Broader task-related maps explained more variance, with the strongest models explaining up to 32.3% of the HCP general factor but still did not account for most of the observed spatial structure. Together, these findings indicate that the HCP general factor overlaps with canonical task-related organization but is not reducible to it.

### 2.3 Reproducible Brain-Wide Communities in Individual Factor Representations

Community detection applied to individual-specific factor scores from both cohorts replicated topographically distinct communities (Fig. 4a, C1–C4). The modal partition across 1,000 Louvain iterations revealed four communities in Cohort 1 and five in Cohort 2. However, the final consensus solution contained five communities in both cohorts, indicating that a frequently occurring community (>50% of runs) in Cohort 1 was further subdivided in the consensus.

**Figure 4.**
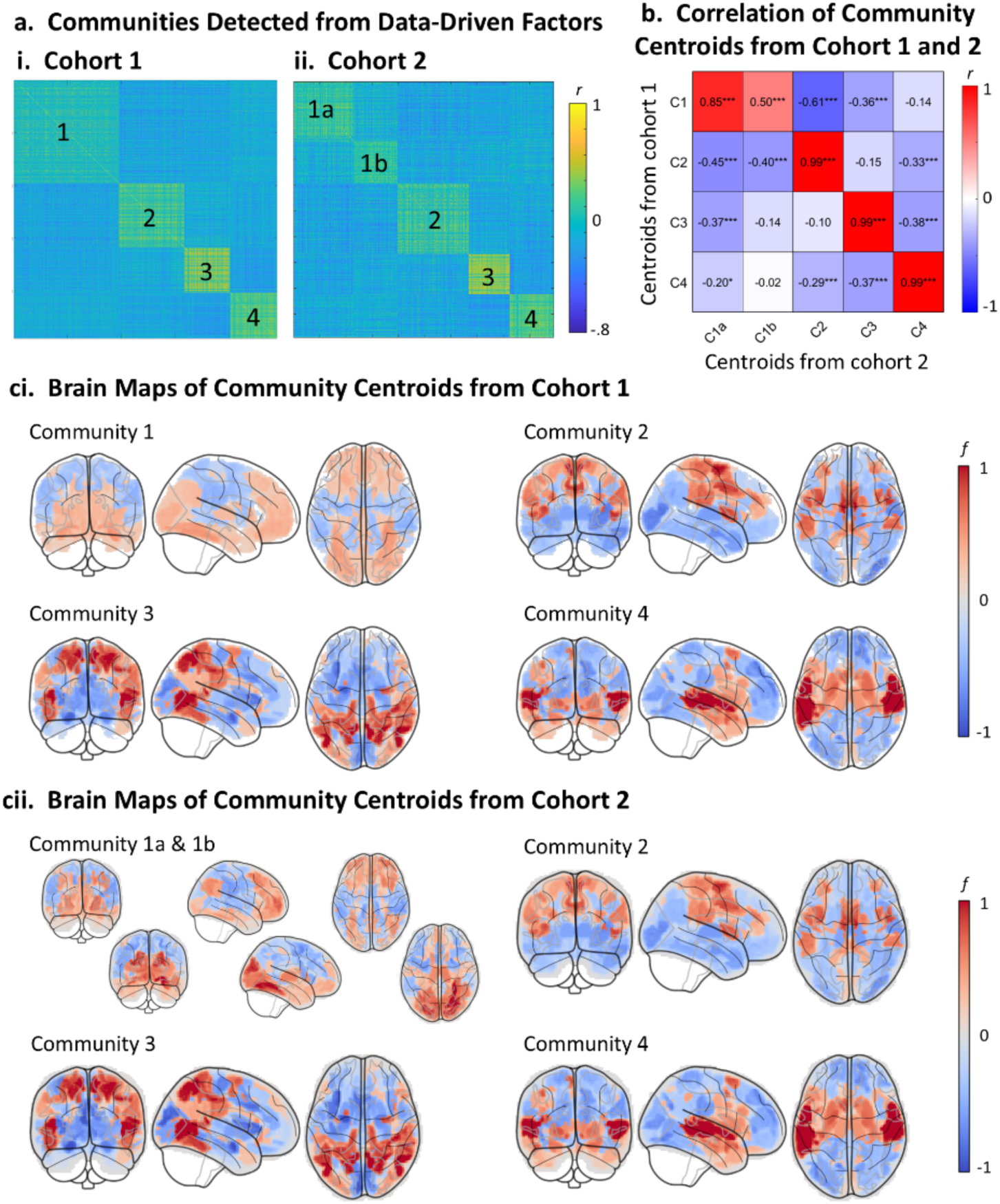
Community detection using data-driven factors reveals reproducible and functionally distinct factor communities across cohorts. (ai) Consensus clustering of correlation matrices from data-driven factor scores in Cohort 1 identified four stable communities. (aii) Community detection in Cohort 2 yielded a similar five-community structure, with Community 1 from Cohort 1 corresponding to two communities (1a, 1b). (b) Correlations between community centroid maps derived from Cohort 1 and Cohort 2. Centroids from both cohorts exhibit strong cross-cohort correspondence, particularly for C2, C3 and C4 (*r* > 0.99). C1-4: Community 1-4. (ci, cii) Brain maps of the resulting community centroids from Cohort 1 and Cohort 2, respectively. Each centroid reflects the mean spatial topography of factor maps assigned to that community. Consistent patterns of activation were observed across cohorts, indicating stable and reproducible community structure. Warm colors denote positive factor scores; cool colors denote negative factor scores.

Stability of the recovered community structure was high in both cohorts, with adjusted Rand index (ARI) values indicating strong agreement between individual runs and the consensus (Cohort 1: ARI = 0.893, SD = 0.127; Cohort 2: ARI = 0.956, SD = 0.095). These spatial patterns were consistent across both training and validation cohorts, with high reproducibility (Pearson correlation coefficient, *r* = .5-.99) between matched centroids of Cohort 1 and Cohort 2 (Fig. 4b; Supplementary Table 9). Each community centroid exhibited a characteristic spatial topography (Fig. 4c): C1 was predominantly prefrontal and occipital activation, C2 engaged somatomotor and subcortical regions, C3 included the parietal cortex and fusiform gyri, and C4 was centered in temporal and subcortical structures.

To evaluate external validity, we compared these data-driven centroids with group-level factors from an independent dataset based on group-averaged task-fMRI maps from Neurovault^7^. The centroids showed significant and differential alignment with both data-driven and RDoC-specific factor maps from that study (Fig. 5; Supplementary Table 10), supporting the interpretability and robustness of the identified community structures. Notably, centroids in both cohorts aligned more strongly (both positively and negatively) with RDoC-defined factors related to the sensorimotor systems and positive valence systems, and less so with those for social processes, cognitive systems, or negative valence systems (Figure 5aii, bii). This suggests that the former domains are associated with more stereotyped and robust functional architectures in the HCP tasks, making them more detectable in both individual- and group-level analyses.

**Figure 5.**
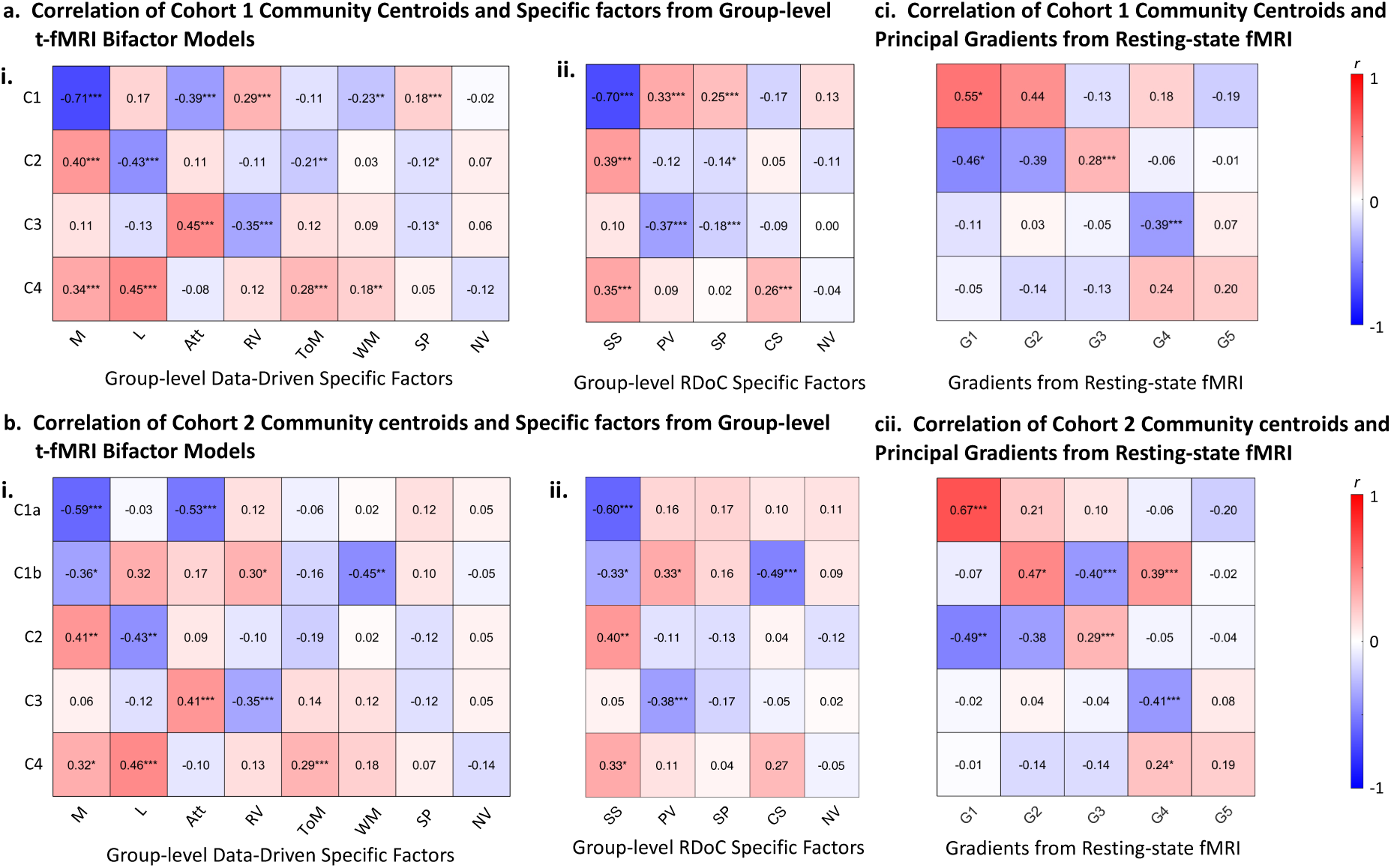
Community centroids align with specific factors from group-level bifactor models and resting-state gradients. (a) Correlation of community centroids from Cohort 1 with group-level bifactor model factors from Quah et al.^7^. (ai) Correlations with data-driven specific factors. (aii) Correlations with RDoC-based specific factors. (b) Correlation of community centroids from Cohort 2 with group-level bifactor model factors from Quah et al.^7^. (bi) Correlations with data-driven specific factors. (bii) Correlations with RDoC-based specific factors. Columns for (a) and (b) have been ordered from highest total correlation coefficient to lowest. (ci–cii) Correlations between task-derived community centroids and the resting-state connectivity gradients from Margulies et al.^12^. P-values have been adjusted for spatial autocorrelation^22^. Additionally, p-values here were corrected for multiple comparisons using the Benjamini–Hochberg false discovery rate (FDR) procedure. C1-4: Community 1-4; M: Motor; L: Language; Att: Attention; ToM: Theory-of-Mind; RV: Reward Valuation; WM: Working Memory; SP: Social Processing; NV: Negative Valence. G1-5: Gradient 1-5. \**p_adj_* < .05, \*\**p_adj_* < .01; \*\*\**p_adj_* < .001.

To further evaluate convergence with other established functional brain topographies, we correlated task-derived community centroids with the gradients of resting-state fMRI reported by Margulies et al.^12^. In both cohorts, the centroids showed significant associations with the first four gradients. Specifically, centroid 1 in Cohort 1 and 1a in Cohort 2 are positively associated with the principal gradient, gradient 1. Additionally, centroid 1 in Cohort 1 is positively associated with gradient 2. Centroid 1b in Cohort 2 is positively associated with gradients 2 and 4 and negatively associated with gradient 3. Centroid 2 in both cohorts is positively associated with gradient 3 and negatively associated with gradients 1 and 2. Finally, centroids 3 and 4 in both cohorts are negatively and positively associated with gradient 4, respectively. All p-values were corrected for multiple comparisons using the Benjamini–Hochberg false discovery rate (FDR) procedure (adjusted values in Supplementary Table 10). These results support the interpretation of task-derived communities as canonical axes of functional brain organization, grounded in both task and resting-state systems.

Finally, we assessed within-subject consistency of community assignments by computing Shannon entropy of factor map distributions across communities. In both the four-community solution from Cohort 1 (mean = 1.83, median = 1.92; maximum possible = 2.00) and the five-community solution from Cohort 2 (mean = 2.13, median = 2.24; maximum possible = 2.32), entropy values were close to the theoretical maximum. This indicates that most participants exhibited diverse factor–community mappings, with maps distributed across multiple modules rather than concentrated in a single one.

### 2.4 Community Alignment Predicts Individual Cognitive Performance

We next tested whether individual alignment with community-level factor representations was associated with behavioral performance on working memory and relational reasoning tasks. These tasks were selected because they were the only ones in the HCP dataset with well-quantified and sufficiently variable performance measures (see Supplementary Methods 1). Other tasks lacked usable metrics, showed ceiling effects, or exhibited low variability, limiting their suitability for individual-level correlation analyses.

To maximize statistical power, this analysis included all participants from the HCP dataset (*N* = 962). For each subject, we computed two similarity indices between their average task activation map and a community centroid: (i) Pearson correlation and (ii) continuous Dice similarity. We selected centroids from Community 4 (C4) and Community 3 (C3) in Cohort 1, as these showed the strongest correlations with group-level factors related to working memory and attention, respectively (Figure 5ai). Associations between similarity and behavior were estimated using linear mixed-effects models, with Parental ID included as a random intercept to control for genetic relatedness.

Higher alignment to community centroids was significantly associated with better behavioral performance. Specifically, similarity to the C4 centroid was positively associated with accuracy on the working memory (*β* = 0.115, *SE* = 0.030, *t*(941) = 3.81, *p\*\*\** <.001; Dice: *β* = 0.174, *SE* = 0.032, *t*(957) = 5.68, *p*\*\*\*< .001). Similarly, similarity to the C3 centroid was positively associated with relational reasoning accuracy (*β* = 0.088, *SE* = 0.032, *t*(957) = 2.72, *p*\*\* = .007). The corresponding Dice-based association was positive but marginal (β = 0.065, SE = 0.033, t(953) = 1.95, p = .051) (Figure 6ai, bi; full Dice results in Supplementary Figure 5).

**Figure 6.**
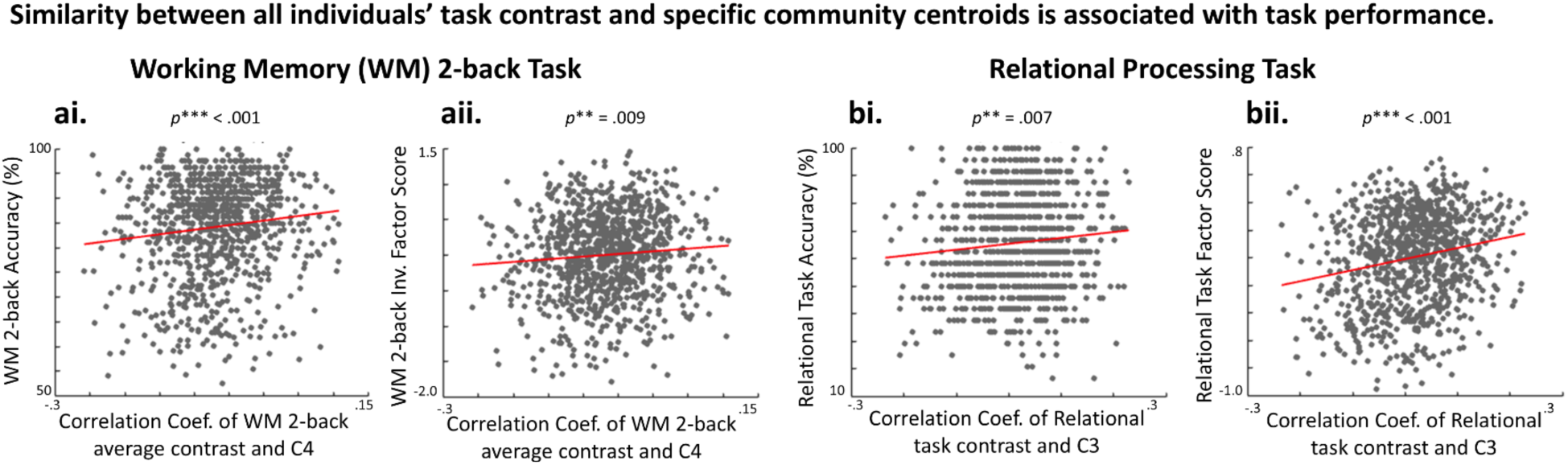
Similarity between individuals’ task contrast and specific community centroids predicts behavioral performance in mixed-effects models. Similarity (quantified by Pearson correlation) was entered as a fixed effect, with Parental ID included as a random intercept to control for genetic relatedness. Higher similarity between an individual’s average task map and community centroid was associated with better performance on the working memory and relational processing tasks. This association was observed both in raw task accuracy (panels ai and bi) and in task performance factor scores derived using EFA (panels aii and bii). Subpanels (i) show predictions for raw accuracy; subpanels (ii) show predictions for latent performance factor scores derived via EFA of accuracy and reaction time (working-memory factor inverted so higher = faster and more accurate responses).

We next examined whether these similarity measures also predicted latent cognitive ability, derived using EFA from both accuracy and reaction time metrics. For working memory, the extracted factor loaded negatively on accuracy and positively on reaction time, indicating that higher scores reflected slower and less accurate performance. To aid interpretation, we inverted these scores so that higher values corresponded to better performance (i.e., faster and more accurate responses). For relational processing, both accuracy and reaction time loaded positively, meaning higher scores reflected greater accuracy with longer response times—a speed–accuracy tradeoff factor.

Similarity to the C4 and C3 centroids significantly predicted these latent performance factor scores (Figure 6aii, bii). Specifically, similarity to the C4 centroid was positively associated with working memory factor scores (Pearson’s: *β* = 0.083, *SE* = 0.032, *t*(946) = 2.63, *p\*\** = .009; Dice: *β* = 0.131, *SE* = 0.032, *t*(942) = 4.06, *p*\*\*\* < .001), and similarity to the C3 centroid was positively associated with relational processing factor scores (Pearson’s: *β* = 0.198, *SE* = 0.032, *t*(956) = 6.14, *p*\*\*\* < .001; Dice: *β* = 0.190, *SE* = 0.033, *t*(952) = 5.73, *p*\*\*\* < .001).

These findings confirm that individual similarity to shared spatial motifs derived from data-driven communities is not only associated with raw task accuracy but also reflects generalizable latent cognitive abilities across tasks. Dice-based results are shown in Supplementary Figure 5.

### 2.5 Behavioral factor structure reveals complementary organization across levels

To assess whether the observed model structure extends beyond neural data within HCP, we examined the latent organization of HCP behavioral measures at the group level (Supplementary Table 11). Unlike the neural factor models, which were estimated separately within each participant because brain parcels provide repeated observations across task contrasts, behavioral factor models were necessarily fit at the group level because the repeated observations in the behavioral data are participants rather than measures. Data-driven measure-to-factor mappings were derived by EFA in one cohort and applied unchanged to the other cohort, and vice versa. Because these behavioral CFAs are defined on cohort-level covariance matrices, model fit was quantified using leave-one-out refitting: on each iteration, one participant was omitted, the covariance matrix was recomputed from the remaining participants, and the model was refit. This procedure tests whether the relative ordering of models is stable to perturbations of the cohort-level covariance structure, although it is not a participant-level held-out test fully analogous to the neural analyses.

Across both HCP cohorts, cross-cohort transported data-driven behavioral models showed better fit than RDoC-based models across all indices (AIC, CFI, RMSEA, TLI; all p < .0001). Adding a general factor to the RDoC model also improved fit relative to the RDoC specific factor model alone. In contrast to the neural results, however, the best-fitting HCP behavioral model was the data-driven specific factor model, which outperformed the data-driven bifactor model in both cohorts (Cohort 1: ΔAIC = −45.97; Cohort 2: ΔAIC = −157.76; all indices p < .0001; Supplementary Figure 6; Supplementary Table 12). This suggests that behavioral variance in the HCP battery is better captured by multiple correlated latent factors than by a single dominant orthogonal general factor.

Together, the HCP behavioral analyses indicate that data-driven models outperform RDoC-based models, while the balance between general and specific variance differs from the neural analyses. Bifactor quality indices for the HCP data-driven behavioral bifactor models are reported in Supplementary Table 13.

### 2.6 Data-driven Community Centroids Capture Topological Structure Better than RDoC Domains

To evaluate how well different annotation schemes capture the global organization of individual brain activity, we applied the Mapper algorithm from topological data analysis to the contrast maps in Cohort 1 and Cohort 2, separately. Mapper constructs a simplified graph representation of high-dimensional data by applying binning and local clustering to a low-dimensional embedding. In the resulting graphs, nodes represent clusters of similar contrast maps, and edges connect nodes with overlapping contrasts. We annotated each node using two alternative schemes: (i) RDoC domain labels, based on the task-condition mappings; and (ii) data-driven community centroids, based on the highest spatial correlation between contrast maps and centroid patterns (Figure 7a).

**Figure 7.**
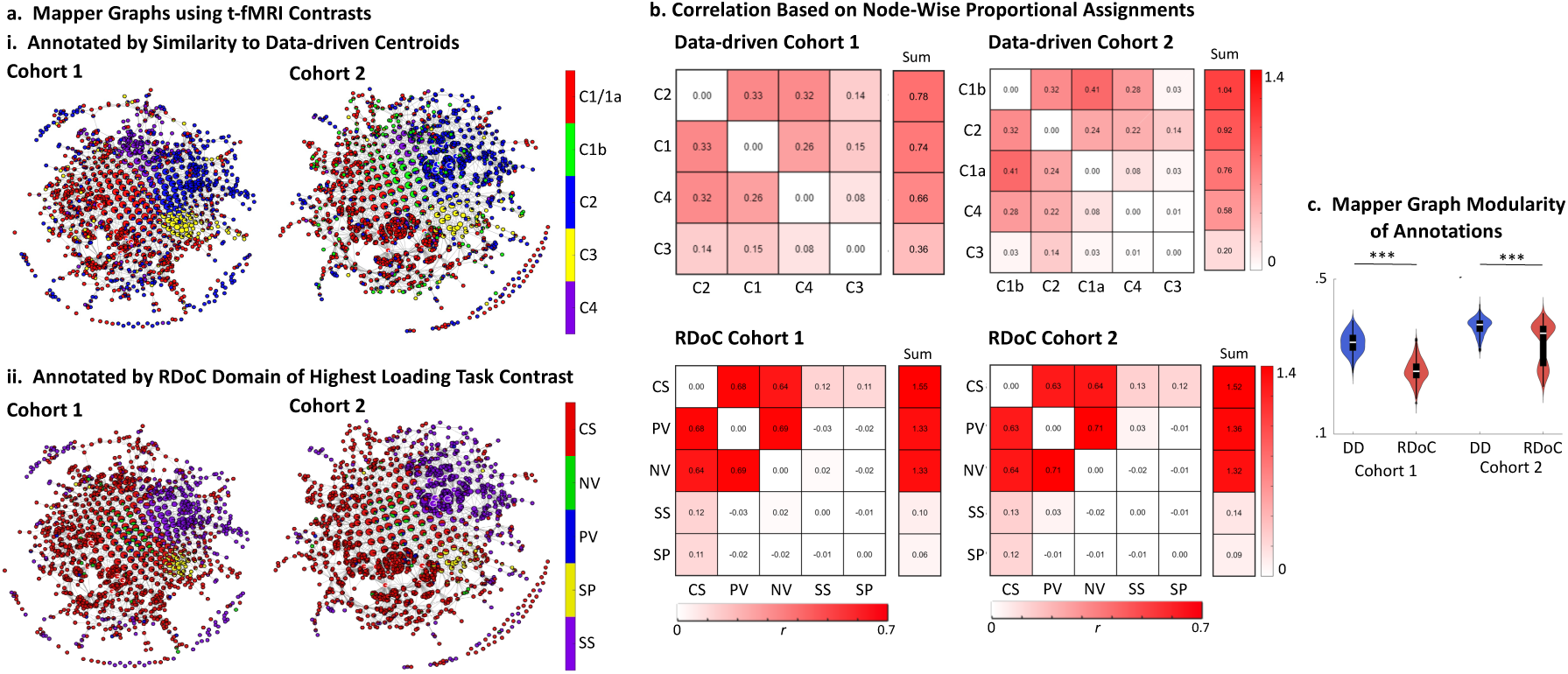
Topological analysis of individual contrast maps using Mapper. (a) Mapper graphs constructed from individual contrast maps from Cohort 1 and Cohort 2, annotated using either data-driven community centroids (top) or RDoC domain assignments (bottom). Each node represents a cluster of similar activation patterns, with edges connecting nodes sharing overlapping contrast maps. (b) Correlation matrices showing the similarity between annotation groups, based on node-wise proportions. For each node in the Mapper graph, we calculated the proportion of contrasts belonging to each community or RDoC domain. The resulting matrices reflect how often different annotations co-occur within nodes. Lower off-diagonal correlations in the data-driven models indicate that the communities are more distinct and less overlapping. The final column (“Sum”) shows how much each group overlaps with all others, with lower values indicating greater separation. (c) Data-driven community annotations exhibit higher Mapper graph modularity than RDoC domain labels. Violin plots show the distribution of modularity values for data-driven (DD, blue) and RDoC-based (red) annotations applied to Mapper graphs in Cohort 1 and Cohort 2. Modularity quantifies the degree to which annotations align with topological community structure. In both cohorts, data-driven annotations yielded significantly higher modularity, indicating stronger alignment with intrinsic graph structure. \*\*\**p* < .001.

To assess the distinctiveness of each annotation scheme, we computed node-wise assignment proportions and their correlation matrices across nodes (Fig. 7b). For each Mapper node, we calculated the proportion of contrasts assigned to each RDoC domain or data-driven community. The resulting correlation matrices reflect how often different categories co-occur within the same topological neighborhood. Compared to RDoC annotations, data-driven communities showed more moderate off-diagonal correlations, indicating more balanced and less redundant structure. In contrast, RDoC domains showed extreme values (high highs and low lows), reflecting greater inconsistency—some domains were overrepresented across nodes, while others were nearly absent. This pattern suggests that RDoC labels are more unevenly distributed in the intrinsic topological space, whereas data-driven annotations yield more consistent and balanced partitions of brain states.

Crucially, across both cohorts, Mapper graphs annotated with community centroids exhibited higher modularity than those annotated with RDoC domains (Fig. 7c). This indicates that the community-based annotation more closely aligns with the underlying topological organization of individual contrast maps. Together, these findings support the conclusion that data-driven community centroids offer a more accurate and functionally coherent framework for summarizing task-evoked brain activity than expert-defined RDoC categories.

### 2.7 Data-driven structure generalizes to an independent transdiagnostic clinical dataset

To assess generalizability and clinical relevance, we applied the model-comparison framework to an independent transdiagnostic dataset, LA5c^11^, including healthy controls, ADHD, bipolar disorder, and schizophrenia. Across all diagnostic groups and component solutions in the projection analyses, healthy-control-derived data-driven bifactor models outperformed RDoC-based models across all fit indices (RMSEA, AIC, CFI, TLI; all p < .001; Figure 8a; Supplementary Table 14). In the pooled direct-estimation analysis, the data-driven bifactor model showed the best fit, followed by the data-driven specific model, the RDoC bifactor model, and the RDoC specific model (Supplementary Figure 7). The data-driven bifactor model significantly outperformed all other models across fit indices (all p < .001), and both data-driven models outperformed both RDoC-based models, although the data-driven specific versus RDoC bifactor comparison for RMSEA was comparatively modest (p = .037). Bifactor quality indices for the LA5c data-driven task-fMRI bifactor models are reported in Supplementary Table 15.

**Figure 8.**
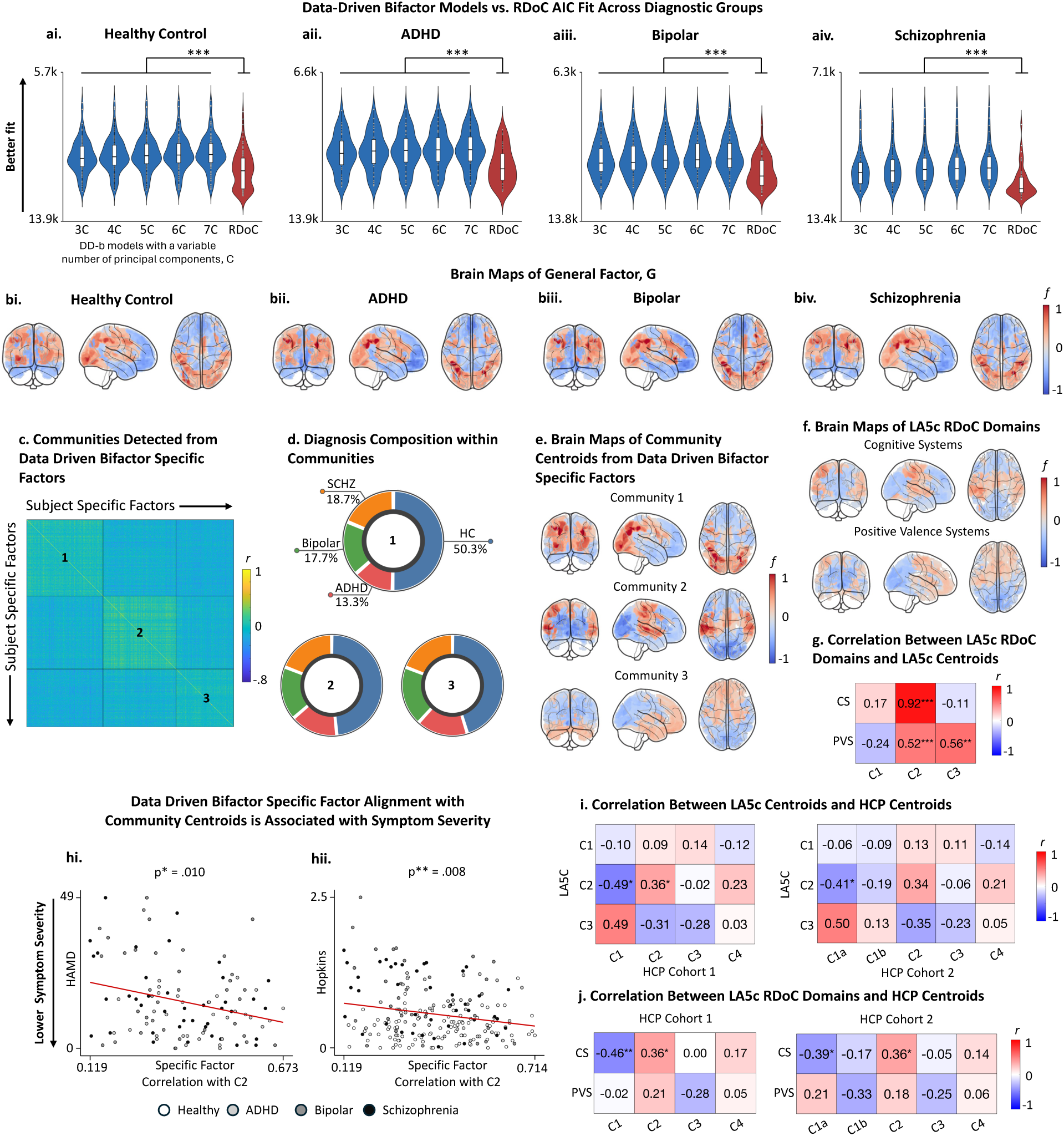
Data-driven bifactor representations generalize to the independent LA5c sample, reveal transdiagnostic organization, and relate to clinical measures. (a) Model fit comparisons between healthy control derived data-driven bifactor (DD-b) component solutions and RDoC models in healthy controls and in participants with ADHD, bipolar disorder, and schizophrenia. For the LA5c healthy control group, DD-b fit was evaluated using repeated 80:20 splits within the healthy control subgroup. For ADHD, bipolar disorder, and schizophrenia, DD-b component solutions derived from LA5c healthy controls were projected onto each clinical group, whereas RDoC models were evaluated within the corresponding group. Comparisons were performed across 3- to 7-component solutions. Across all groups and all tested component numbers, DD-b models outperformed RDoC models on RMSEA, AIC, CFI, and TLI. AIC is shown here; all fit indices are summarized in Supplementary Table 14. (b) Brain maps of the diagnosis-specific general factor, G, estimated separately in LA5c healthy controls and participants with ADHD, bipolar disorder, and schizophrenia. Pairwise parcel-wise correlations among group-average general factor maps are summarized in Supplementary Table 19. (c,d) Community detection of specific factors and diagnostic composition of participants within communities in the LA5c sample. Pie charts show the diagnostic composition of each community. The proportions of ADHD, bipolar disorder, healthy controls, and schizophrenia were similar across the three communities, indicating that community structure was not driven by diagnosis alone. (e,f) Brain maps of LA5c data-driven bifactor centroids and LA5c RDoC classical brain maps. Shown are whole-brain centroid maps derived from the data-driven bifactor specific factors in the LA5c sample and the corresponding LA5c RDoC factor maps. (g) Correlations between LA5c RDoC domains and LA5c centroids. (h) Associations between alignment to data-driven community centroids and clinical measures in the LA5c sample. Community-based alignment was significantly associated with symptom severity. (i) Correlations between LA5c centroids and HCP centroids. (j) Correlations between LA5c RDoC domains and HCP centroids. Corresponding Pearson correlation coefficients are shown in g, i, and j. P values were corrected for spatial autocorrelation using BrainSMASH and then adjusted for multiple comparisons using the Benjamini-Hochberg false discovery rate procedure. *padj < .05; **padj < .01.

We also repeated the *behavioral* model comparison in LA5c. Data-driven measure-to-factor mappings were derived in the LA5c healthy control subgroup and then held fixed when fitting models in the pooled clinical sample (Supplementary Table 16). In LA5c, the best-fitting behavioral model was the data-driven bifactor model, followed by the RDoC bifactor model, the data-driven specific factor model, and the RDoC specific factor model (DD-b > RDoC-b > DD > RDoC; all pairwise p < .0001; Supplementary Figure 8; Supplementary Table 17). Bifactor quality indices for the LA5c data-driven behavioral bifactor model are reported in Supplementary Table 18. Relative to HCP, this stronger support for a general factor may reflect the narrower and more cognitively concentrated LA5c battery.

General factor maps were well correlated across diagnostic groups (Figure 8b), with corresponding pairwise parcel-wise correlations among diagnostic group general factor maps summarized in Supplementary Table 19. To further test whether the LA5c general factor could be explained primarily by canonical task-positive organization, we compared the pooled LA5c general factor map with independent Neurosynth^20^ meta-analytic maps. Using the task-positive association and default mode association maps, the full model explained 9.9% of the variance in the pooled LA5c general factor, leaving the majority of variance unexplained. Broader task-related maps explained more variance, with the strongest model explaining 30.1% of the pooled LA5c general factor, but still did not account for most of the observed spatial structure.

Direct correspondence between the pooled LA5c and HCP general factor maps was stronger than correspondence with any single Neurosynth-derived template: the pooled LA5c general factor correlated with the HCP general factor maps at r = .583, whereas its strongest single-template Neurosynth correspondence was r = .493 (Supplementary Tables 7 and 8). Together, these findings support the reproducibility of the recovered pattern across datasets and indicate that the general factor overlaps with canonical task-related organization but is not reducible to it.

Communities detected from data-driven bifactor specific factors are shown in Figure 8c. The diagnostic composition of participants within communities was similar across groups (Figure 8d; Supplementary Table 20), indicating that the recovered motifs were not simple relabeling of diagnostic categories but instead reflected transdiagnostic dimensions of brain organization. Brain maps of community centroids from data-driven bifactor specific factors are shown in Figure 8e, and RDoC classical factor maps are shown in Figure 8f. Community centroids identified in LA5c showed significant alignment with LA5c RDoC classical factor maps (Figure 8g) after correction for spatial autocorrelation and multiple comparisons (Supplementary Table 21).

To assess clinical relevance, we examined associations between community-based representations and symptom severity. Alignment of individual data-driven specific factors with data-driven community centroids was significantly associated with symptom measures (Figure 8h; Supplementary Table 22), such that greater alignment corresponded to lower symptom severity, whereas RDoC-based template alignments did not show significant associations (Supplementary Table 22). We next assessed cross-dataset correspondence of the recovered spatial motifs. Community centroids identified in LA5c also showed significant alignment with HCP-derived centroids (Figure 8i), and LA5c RDoC classical factor maps showed significant alignment with HCP-derived centroids (Figure 8j), after correction for spatial autocorrelation and multiple comparisons (Supplementary Table 21). Together, these findings indicate that the data-driven structure generalizes across diagnostic groups and captures behaviorally relevant variation in clinical populations.

## 3. Discussion

### 3.1 Validating the RDoC at the single-participant level

The RDoC framework has been widely applied in psychiatric neuroimaging research, yet its empirical grounding—particularly the mapping between functional domains and neural circuits—remains underdeveloped. Here, we present a latent factor analysis of task-based fMRI data aimed at refining RDoC to more closely align with the functional organization of brain activity and to identify the large-scale organization of task-evoked brain circuits. While prior efforts have derived latent dimensions from neuroimaging and behavioral data at the group level, it has remained unclear how well such models capture subject-specific activation patterns or generalize across datasets, clinical populations, and measurement domains.

Our study systematically evaluated the structure, generalizability, and behavioral relevance of data-driven factor models estimated at the individual level and then tested whether those advantages extended across measurement domains and datasets. In neural data, data-driven models consistently outperformed RDoC-based alternatives in statistical fit, replicated across HCP cohorts, generalized to an independent transdiagnostic LA5c sample, and captured clinically relevant variation in symptom severity. Complementary behavioral analyses in HCP and LA5c further showed that data-driven models also outperformed RDoC-based behavioral models, although the relative support for bifactor versus specific factor structure varied across datasets. The general factor derived from the neural bifactor models revealed a reproducible macroscale axis from visual–attentional to default mode systems. Beyond the general factor, data-driven models recovered stable, topographically distinct functional motifs associated with cognitive performance and better captured the topological structure of task-evoked brain states. Together, these findings provide a scalable framework for advancing precision neuroscience and refining psychiatric ontologies such as RDoC.

We first showed that bifactor models derived from EFA of individual-level task-fMRI data offered significantly better model fit than RDoC-based models across multiple indices (AIC, RMSEA, CFI, TLI). This advantage was consistent across two independent HCP cohorts and held when data-driven solutions from one cohort were projected onto the other. Because bifactor models can sometimes overfit correlated-factor alternatives, we also quantified recommended bifactor quality indices and found a strong, highly replicable general factor alongside recoverable, albeit weaker, specific factors. Because the mapping between task contrasts and RDoC domains is not uniquely defined, we also tested whether model comparisons depended on the prespecified contrast-to-domain assignment. The observed RDoC mapping substantially outperformed randomly reassigned alternatives, indicating that model-comparison results were not invariant to assignment structure and that the prespecified mapping captured meaningful alignment between task design and domain definitions. At the same time, this mapping should be interpreted as one plausible operationalization rather than a definitive classification. Future work should evaluate alternative mappings, including probabilistic or multi-domain assignments, to further refine the relationship between task paradigms and dimensional constructs.

### 3.2 Evidence for the general factor

The general factor reflects a latent, low-dimensional axis of task-evoked brain organization, characterized by positive loadings in visual, dorsal attention, and frontoparietal networks, and negative loadings in default mode, auditory, and somatomotor networks. Importantly, this factor is derived from the shared covariance across task contrasts and therefore captures a domain-general structure of brain activity rather than simple activation differences.

Quantitatively, this axis shows its strongest correspondence with Gradient 2 and a moderate correspondence with Gradient 3 from Margulies et al.^12^, indicating that it is anchored in a visual–attentional system that spans sensory processing and task-positive control networks, while opposing default mode and non-task-engaged systems. Notably, the general factor does not align with the principal unimodal–transmodal gradient (Gradient 1), suggesting that task-evoked activity is organized along a distinct but related axis of cortical hierarchy.

This finding provides a link between extrinsic (task-driven) and intrinsic (resting-state) brain organization, demonstrating that a similar macroscale structure emerges from task-based covariance without relying on resting-state connectivity. In this sense, the general factor reflects a task-derived analogue of intrinsic cortical gradients, supporting a unifying framework in which both spontaneous and evoked activity are constrained by shared large-scale organizational principles.

Functionally, the emergence of a single brain-wide factor spanning multiple task domains supports the presence of a domain-general dimension of brain activity^7^. This interpretation is consistent with prior work showing that variance across diverse tasks is dominated by shared engagement of visual and attention systems alongside suppression of default mode activity^23^. More broadly, these findings reinforce the view that the brain’s functional architecture is best described along continuous gradients and hierarchies, rather than discrete domain partitions^24^.

Comparing the general factor to independent meta-analytic maps from Neurosynth^20^ further clarifies its interpretation. Although it shares some variance with canonical task-related and default mode patterns, the amount of variance explained by Neurosynth templates remained incomplete, indicating that the general factor is not simply a relabeling of known task-positive networks. Rather, it reflects a broader dimension of shared task-evoked activity that partially overlaps with canonical task-related organization while retaining additional spatial structure.

### 3.3 Evidence for specific motifs and communities

We next applied community detection to uncover shared structure among specific factors. This analysis identified four stable community motifs in Cohort 1, each with distinct spatial signatures. In Cohort 2, these patterns were replicated, with one community (C1) subdivided into two closely related subcommunities (C1a and C1b). The most prominent community (C1) involved co-activation of primary visual cortex and DMN regions—notably posterior cingulate and medial prefrontal cortex. This topography aligns with the principal gradient of cortical organization^12^ and with the first complex principal component in resting-state data^25^, supporting its interpretation as a core organizational axis of human brain function. The presence of visual–DMN co-activation likely reflects the strong visual demands embedded in many HCP tasks.

Importantly, these spatial motifs (or communities) were reproducible across cohorts and aligned with both group-level data-driven factors and canonical task contrasts. This suggests that they reflect recurrent, interpretable modes of co-activation, transcending individual tasks or datasets. The presence of these discrete motifs supports the formulation that large-scale brain activity can be decomposed into a relatively small number of interpretable functional modes. Notably, these communities aligned most strongly with RDoC domains associated with sensorimotor and positive valence systems, and less so with cognitive, social, or negative valence domains—replicating a pattern seen in our prior group-level work^7^. The stronger alignment for sensorimotor and reward domains may reflect their greater spatial stereotypy, consistent with prior meta-analyses^26,27^, while more distributed and context-sensitive processes (e.g., negative valence, cognitive control) may resist simple spatial encapsulation^5,28^.

Our finding that task-derived community centroids align with the principal gradients of resting-state functional connectivity described by Margulies et al.^12^ provides strong evidence for their biological validity. The positive alignment of fronto-occipital centroids (C1/1a) with the principal gradient indicates that the task-derived factors here recapitulate the unimodal–to–transmodal axis of brain organization observed at rest, whereas the associations of centroids 2–4 with higher-order gradients indicate sensitivity to additional axes of functional specialization beyond the dominant unimodal–transmodal spectrum. The reproducibility of these relationships across independent cohorts further supports that the task-derived centroids here reflect stable, canonical dimensions of brain functional organization. Together, these findings support the view that individual-level task activation patterns are constrained by the same large-scale topographic gradients that govern intrinsic functional connectivity.

### 3.4 Behavioral associations

Behaviorally, we found that individuals whose activation patterns aligned more closely with certain centroids—especially C4 and C3—showed better performance on working memory and relational reasoning tasks. These associations held for both raw accuracy and EFA-derived latent performance scores. This supports the idea that spatial similarity to population-level motifs, rather than overall activation magnitude or connectivity, offers a meaningful index of task-relevant network engagement. These results extend prior findings^29,30^ by demonstrating that centroid similarity predicts trait-like cognitive variation and may offer a scalable approach to individualized phenotyping.

### 3.5 Neural-behavioral convergence and divergence in latent factor structure

At the neural level, the data-driven bifactor model consistently provided the best fit, indicating a robust general component in task-evoked brain activity. At the behavioral level, the best-fitting model was also data-driven in both datasets, but the full ordering differed across samples. In HCP, the data-driven specific factor model provided the best fit, followed by the data-driven bifactor model. In LA5c, by contrast, the data-driven bifactor model provided the best fit, followed by the RDoC bifactor model and then the data-driven specific factor model. Thus, the behavioral results do not support a uniform conclusion that behavior is best captured by specific factors; rather, they indicate that the balance between general and specific variance in behavior is dataset-dependent.

This pattern suggests that support for a general factor in behavioral data depends in part on the breadth and composition of the measurement battery. The HCP behavioral battery spans a broader range of cognitive, affective, social, arousal/regulatory, and sensorimotor constructs, which may favor a more differentiated structure captured by correlated specific factors. By contrast, the LA5c behavioral battery is more cognitively concentrated and lacks several negative valence and arousal/regulatory measures, plausibly increasing covariance across measures and strengthening support for a general factor. Notably, the fact that the RDoC bifactor model outperformed the data-driven specific factor model in LA5c further suggests that, in narrower behavioral batteries, the inclusion of a general factor may matter more than whether the assignments are theory-driven or empirically derived. In contrast, task-evoked neural activity showed more consistent evidence for a robust general factor, potentially reflecting domain-general processes such as attentional engagement, task set maintenance, or global task demands^15^. Simultaneously, we cannot exclude the possibility that differences in neural and behavioral model ordering partly reflect the distinct levels at which the models were estimated, with neural data modeled at the individual level and behavioral data at the group level, in addition to genuine differences in the organization of shared variance across measurement domains.

These findings refine the implications for the RDoC framework. Rather than suggesting a single latent architecture that generalizes uniformly across levels of analysis, our results indicate that the most stable conclusion is that empirically derived models provide an important benchmark for refining predefined ontologies, while the relative contribution of general and specific factors varies across modality, dataset, and measurement battery. Neural task responses appear to contain a consistently strong domain-general component, whereas behavioral structure is more sensitive to which constructs are sampled and how broadly the measurement space is covered. This reinforces the need to validate RDoC constructs across multiple datasets and measurement domains and highlights the value of data-driven approaches in refining cross-level mappings.

### 3.6 Generalization to transdiagnostic clinical populations

A key advance of the present work is the extension of findings to a transdiagnostic clinical dataset. The general factor maps in LA5c were well correlated across healthy and clinical groups, consistent with a transdiagnostic organization of shared task-evoked variance. While primary analyses were conducted in a healthy cohort, replication in LA5c demonstrates that the data-driven structure is not specific to normative populations, but generalizes across multiple psychiatric conditions. The similar community composition across diagnostic groups further supports a transdiagnostic organization of brain activity, consistent with dimensional approaches to psychopathology. Moreover, the association between community-based representations and symptom severity suggests that these data-driven features capture clinically meaningful variation, providing a potential bridge between neural organization and psychiatric symptomatology. In contrast, RDoC-based template alignment did not show comparable associations, highlighting the potential value of empirically derived representations for refining dimensional frameworks.

### 3.7 Mapper and topological validation

Finally, we used TDA to show that data-driven community centroids better capture the functional organization of task-evoked brain states than predefined RDoC domains. Mapper graphs annotated with community centroids exhibited higher modularity, suggesting better alignment with the data’s natural topological structure. In contrast, RDoC annotations—especially those for cognitive and valence domains—showed greater inter-domain overlap and more uneven distribution. These findings echo prior work showing that Mapper reveals modular and behaviorally relevant brain states^13^, and they reinforce the view that empirically derived models offer more topologically coherent functional partitions than expert-defined taxonomies.

### 3.8 Implications for refining RDoC

These findings offer actionable insights for the refinement of cognitive ontologies such as RDoC. First, our results suggest that RDoC constructs could benefit from greater empirical grounding through systematic bottom-up modeling of neuroimaging data at the individual level. Instead of defining domains solely through top-down consensus or behavioral theory, a hybrid approach that integrates expert annotation with data-driven spatial motifs may yield more biologically valid and reproducible constructs. Second, the consistent emergence of a domain-general factor spanning multiple tasks points to the utility of incorporating continuous dimensions—such as macroscale activation gradients—alongside discrete domain categories. Such gradients may serve as organizing axes that unify multiple constructs or identify shared variance across domains. The behavioral and transdiagnostic results further indicate that no single latent architecture should be assumed to generalize uniformly across measurement domains, datasets, and populations; instead, empirically derived models provide a benchmark for refining predefined ontologies under different measurement conditions. Third, adopting a graph-based or topological lens, as shown through Mapper analysis, provides a promising route for assessing whether cognitive taxonomies align with the intrinsic geometry of brain function. We propose that future iterations of RDoC include empirical benchmarking tools such as factor coherence, spatial reproducibility, behavioral relevance, and topological modularity, enabling systematic validation and revision of the proposed domains. Together, these principles can help re-anchor psychiatric constructs in neurobiologically informed, scalable frameworks—advancing RDoC’s original mission of building a precision mental health science.

### 3.9 Limitations & Future directions

Several limitations should be acknowledged. First, although the neural model comparison generalized to LA5c, domain coverage was limited and uneven in both HCP and LA5c. Several domains were represented by only a small number of measures or paradigms, and some were only partially sampled, limiting our ability to capture the full breadth and heterogeneity of constructs within each domain. Our findings should therefore be interpreted as applying to the available operationalizations of RDoC in these datasets rather than as a comprehensive test of the framework. Future work should incorporate broader coverage and multiple paradigms per domain, particularly for underrepresented domains such as arousal/regulatory systems.

Second, although the present study models brain organization at the level of individual participants, it does so within a common parcellated template space and therefore does not capture individualized network boundaries or subject-specific topological variation as in precision functional mapping^19,31,32^. Instead, our focus is on identifying subject-specific latent structure within a standardized representation, enabling direct comparison across individuals and cohorts. Future work incorporating individualized parcellations and subject-specific network topology may further refine these representations and provide a more detailed account of inter-individual variability. In this sense, the present approach is complementary to precision mapping methods, emphasizing generalizable latent structure across individuals rather than individualized boundary estimation.

Third, while Mapper provides rich visualizations of data topology, its parameters are not biologically grounded and may influence results. Future work should therefore evaluate parameter sensitivity and broader cross-dataset replication. Fourth, although our models predict behavior, variability in reaction time may itself modulate BOLD responses^33^, potentially confounding associations between factor scores and latent performance. Finally, integrating multimodal data, including genetics, physiology, and richer symptom phenotyping, and examining variation across sex, age, development, and additional clinical subgroups would further strengthen and contextualize our conclusions.

### 3.10 Conclusions

Despite these limitations, our findings provide strong evidence that empirically derived factor models outperform predefined RDoC models in explaining the organization of task-evoked brain activity and, in complementary analyses, behavioral structure. In neural data, these models reveal two complementary levels of organization: (i) a reproducible general factor that reflects a macroscale axis spanning visual–attentional and default-mode systems, and (ii) topographically distinct, behaviorally and clinically relevant communities that generalize across individuals, datasets, and diagnostic groups. In behavioral data, the optimal empirical model was also data-driven, although the balance between general and specific structure depended on the breadth and composition of the measurement battery. By embedding subject-specific brain activation patterns in scalable, interpretable, and empirically derived frameworks, this work provides a concrete step toward developing neurobiologically grounded cognitive ontologies and supports ongoing efforts to refine and operationalize RDoC in the service of precision psychiatry^34^.

## 4. Methods

### 4.1 Study Cohort

We used task-fMRI data from the publicly available Human Connectome Project (HCP) Young Adult dataset^35^. A total of 962 healthy adults aged 22–35 years with complete task-fMRI data across seven tasks were included; participants with missing data for any task were excluded. Two age-, sex-, race-, and income-matched cohorts were defined for cross-validation: Cohort 1 (N = 412; 52.4% female; mean age = 28.5 years; racial composition: 65.7% White, 14.1% Black, 10.7% Hispanic/Latino, 7.5% Asian, 1.9% Other; income distribution: 46.5% high / 53.5% low) and Cohort 2 (N = 329; 55.0% female; mean age = 28.8 years; racial composition: 69.3% White, 13.5% Black, 8.9% Hispanic/Latino, 7.1% Asian, 1.2% Other; income distribution: 54.3% high / 45.7% low). All participants were unrelated within each cohort.

The HCP task battery included seven paradigms aligned with core RDoC domains: (1) a 2-back task assessing working memory; (2) a motor response task targeting motor function; (3) an auditory story task probing language processing; (4) a stimulus matching task assessing emotional processing; (5) a social cognition task probing theory of mind; (6) a relational matching task assessing higher-order reasoning; and (7) a gambling task assessing reward sensitivity. These tasks are described in detail by Barch et al.^36^.

To assess generalizability beyond the healthy HCP sample, we additionally analyzed an independent transdiagnostic dataset (LA5c^11^), including healthy controls (n = 130) and participants with ADHD (n = 43), bipolar disorder (n = 49), and schizophrenia (n = 50). LA5c was used for external neural replication, behavioral model comparison, and symptom-correlation analyses.

### 4.2 Contrast Map Processing

Individual-level task-fMRI contrast maps were obtained from the Human Connectome Project (HCP) S1200 data release in CIFTI format. All data were processed using the HCP minimal preprocessing pipeline, which includes gradient distortion correction, motion correction, EPI distortion correction, spatial normalization to standard space, and surface-based alignment^37,38^. First-level task analyses were performed within the HCP framework using standardized general linear models (GLMs) to generate contrast maps for each task condition, with extensive quality control incorporated as part of the HCP processing stream.

In the present study, we used these HCP-derived contrast maps and did not reprocess raw fMRI data. For each participant, contrast maps were combined across the left-right (LR) and right-left (RL) phase-encoding runs to generate a single subject-level contrast map, thereby integrating data across independent runs and reducing run-specific dependencies. The resulting maps were then parcellated into 347 regions, comprising 333 cortical parcels from the Gordon atlas^19^ and 14 subcortical regions from the Harvard–Oxford atlas^39^. These atlases were selected for their fine-grained, functionally informed parcellations and their broad validation in both task-based and resting-state fMRI studies.

LA5c contrast maps were generated using the dataset-specific preprocessing and first-level modeling pipeline and then parcellated to the same 347-region atlas used for HCP to enable direct cross-dataset comparison.

### 4.3 Factor Models

We compared four model types: (i) an RDoC-specific factor model (canonical RDoC model), (ii) an RDoC bifactor model, (iii) a data-driven specific factor model, and (iv) a data-driven bifactor model. In the bifactor variants, a general factor (onto which all task contrasts loaded) was included alongside domain- or EFA-derived specific factors. In the specific factor variants, only domain- or data-driven specific factors were modeled, without a general factor. The general factor captures variance shared across all task contrasts, while specific factors reflect variance unique to subsets of contrasts. For both bifactor models, orthogonality constraints were imposed to ensure separation between shared and specific variance components^40^. Complementary group-level behavioral factor analyses are described in Section 4.3.3.

Data-driven models were estimated using a two-step process involving exploratory factor analysis (EFA) followed by confirmatory factor analysis (CFA). In contrast, RDoC-based models were estimated directly via CFA, using predefined domain-to-task assignments. All factor analyses were implemented in R (version 4.3.0) using the psych (version 2.3.3) and lavaan (version 0.6.15) packages.

#### 4.3.1 Data-driven Factor Analysis for each subject

Subject-specific factor models were derived using the following steps: (1) Horn’s parallel analysis to determine the optimal number of specific factors (see below); (2) EFA using principal axis factoring and oblimin rotation; and (3) CFA specifying either a bifactor or specific factor structure.

Each participant’s data consisted of 24 contrast maps (i.e., task conditions), treated as variables, and 347 brain parcels (333 cortical + 14 subcortical) as observations—resulting in a 347 × 24 matrix per subject. EFA was applied to model the covariance structure among the contrast maps based on spatial similarity across parcels. A high factor loading indicated that a task contrast’s spatial activation pattern closely aligned with the topography represented by that factor.

The number of factors to extract was determined using Horn’s parallel analysis^41^, which identifies the point where eigenvalues from the real data intersect those from randomly generated data^42^. Principal axis factoring with oblimin rotation was used to allow for weak correlations among factors. In the subsequent CFA, we used robust maximum likelihood estimation to account for non-normality. Two data-driven models were specified: (i) a bifactor model, where all contrasts loaded on a general factor and one or more EFA-derived specific factors; and (ii) a specific factor model, where only EFA-derived specific factors were included. Consistent with prior literature^43^, specific factors were defined using a threshold of absolute loadings ≥ 0.4.

#### 4.3.2 RDoC Domain Factor Analysis for each subject

RDoC-based models were constructed by grouping whole-brain activation maps into domain-specific factors, based on expert mappings of task descriptions to RDoC domains (Supplementary Table 23). As with data-driven models, we tested both (i) a specific factor model (contrasts loaded only onto their assigned RDoC domains) and (ii) a bifactor model (all contrasts additionally loaded onto a general factor). All RDoC-based models were estimated using CFA with robust maximum likelihood estimation (Fig. 2ai and 2aii).

Task contrasts were assigned to RDoC domains based on the processes targeted by each paradigm, informed by prior literature and established interpretations of the task batteries. HCP contrast-to-domain assignments are summarized in Supplementary Table 23, and the corresponding LA5c task contrast descriptions and RDoC domain assignments are provided in Supplementary Table 24. While some contrasts map naturally onto specific domains, others involve processes that span multiple domains, and therefore the mapping should be interpreted as an operational approximation rather than a definitive classification.

#### 4.3.3 Complementary Group-Level Behavioral Factor Analyses

To complement the individual-level task-fMRI analyses, we analyzed broad sets of behavioral measures in HCP and LA5c at the group level. Behavioral factor models were necessarily estimated at the group level because, unlike the neural analyses where parcels serve as repeated observations across task contrasts, the repeated observations in behavioral data are participants rather than measures; participant-specific behavioral factor models are therefore not identifiable. For the theory-driven (RDoC) models, measures were assigned to domains using pre-assigned RDoC labels. For the data-driven models, measure-to-factor mappings were derived by exploratory factor analysis in an independent sample and then held fixed in the target sample. In HCP, mappings were derived in one cohort and applied to the other cohort, and vice versa. In LA5c, mappings were derived in the healthy control subgroup and applied to the pooled clinical sample. Behavioral measures included in HCP and LA5c are listed in Supplementary Tables 11 and 16, respectively. We evaluated both specific factor and bifactor variants for the theory-driven and data-driven model families using robust maximum likelihood estimation.

Because these behavioral models are defined on group-level covariance matrices, participant-level held-out validation is not directly analogous to the neural analyses. To quantify robustness while minimizing circularity, model fit was evaluated using leave-one-out refitting in the target sample. On each iteration, one participant was omitted, the covariance matrix was recomputed from the remaining participants, the model was refit, and AIC, RMSEA, CFI, and TLI were recomputed. This procedure was repeated once per participant in each sample.

#### 4.3.4 Sensitivity Analysis of Contrast-to-Domain Assignments

To assess the sensitivity of the RDoC comparison to the prespecified HCP contrast-to-domain mapping, we generated null distributions of model fit by randomly reassigning contrast-to-domain labels and refitting the RDoC models. For each permutation, AIC, RMSEA, CFI, and TLI were recomputed, and the percentile rank of the prespecified assignment was calculated relative to the permuted distribution. This analysis tested whether the reported mapping performed better than arbitrary alternatives.

### 4.4 Training and Validation Strategy

To ensure a fair comparison between data-driven and RDoC-based models, we derived the data-driven factor structure from one cohort and tested its generalizability in the other (Supplementary Figure 9). This cross-cohort approach avoids circularity, as data-driven models are inherently optimized to fit the data they are trained on. One cohort was designated as the training set, from which bifactor models were estimated via EFA and CFA. Specific factors were then reduced via principal component analysis (PCA) to extract the top 3–7 components, yielding a low-dimensional representation of the training data.

These PCA components were projected onto the validation cohort by computing a correlation matrix between individual activation maps and the training-derived component scores. A dynamic threshold, set to the p-th quantile (p = 1/k, where k is the number of components), was applied to assign activation maps to components, ensuring balanced factor assignment. This generated individual-level factor scores in the validation set, grounded entirely in the training cohort’s structure.

In contrast, RDoC-based models—being predefined—were directly estimated in each cohort using CFA, with identical factor assignments across training and validation analyses.

To extend the neural model comparison framework beyond HCP, we conducted analogous analyses in LA5c. For projection analyses, data-driven bifactor component solutions were derived in the LA5c healthy control subgroup. Model fit in LA5c healthy controls was evaluated using repeated 80:20 splits within the healthy control subgroup. The resulting healthy control derived component solutions were then projected into the ADHD, bipolar disorder, and schizophrenia groups and compared with RDoC models across 3- to 7-component solutions. As a complementary internal analysis, all four model families (DD, DD-b, RDoC, and RDoC-b) were estimated directly in the pooled LA5c sample.

### 4.5 Model Performance

Model fit was evaluated using robust versions of the Root Mean Square Error of Approximation (RMSEA), Comparative Fit Index (CFI), and Tucker–Lewis Index (TLI), which account for potential non-normality in the data. Additionally, we used the Akaike Information Criterion (AIC), an information-theoretic measure that balances model fit with complexity. Lower AIC values indicate a more favorable trade-off between goodness of fit and parsimony^44^.

### 4.6 Community Detection

To identify shared structure across individuals’ factor representations, we applied Louvain modularity maximization^45^ to the specific factor scores derived from the data-driven bifactor models. Subject-level factor maps were concatenated into a matrix of size (subjects × factors) × regions, and community structure was estimated using an asymmetric treatment of negative edges. This variant preserves meaningful modularity by asymmetrically penalizing anti-correlated nodes. To address the degeneracy of modularity (𝑄) solutions, we ran the algorithm 1,000 times with a fixed resolution parameter (γ = 1). Each run produced a community assignment and associated modularity value. We then computed an agreement matrix based on the frequency of co-assignment across runs, weighted by the modularity of each partition. A consensus clustering procedure was applied to this matrix to derive a stable community solution. Stability was quantified as the adjusted Rand index (ARI) between each run’s partition and the consensus solution, summarized by the mean and standard deviation across all runs.

The resulting communities grouped factor maps with similar spatial profiles. We computed centroid maps—the average activation pattern within each community—for use in downstream analyses, including cross-cohort comparisons and behavioral prediction.

#### 4.6.1 Within-Subject Consistency of Community Assignments

To evaluate the consistency of community assignments across factor maps from the same individual, we computed subject-wise Shannon entropy. For each participant, we tallied how many of their factor maps were assigned to each community and converted these counts into a probability distribution. Entropy was then computed as:

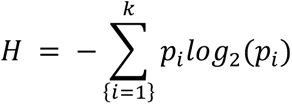

where 𝑝*_i_* is the proportion of maps assigned to the *i-th* community, and 𝑘 is the total number of detected communities. Entropy approaches zero when all maps from an individual fall within a single community and reaches a maximum log_2_(𝑘) when maps are evenly distributed across communities.

### 4.7 Topological Data Analysis

We applied the Mapper algorithm^13,46^, a TDA method, to visualize the intrinsic structure of individual task-evoked brain activation. Mapper constructs a graph-based summary of high-dimensional data, revealing topological features that are often missed by traditional dimensionality reduction techniques.

We began by computing a pairwise Euclidean distance matrix across all contrast maps. A k-nearest neighbor (kNN) graph was constructed to approximate the underlying data manifold, with k set to the square root of the number of data points, following the empirical rule-of-thumb^47^. From the kNN graph, we computed geodesic distances between all data points to better capture the intrinsic structure while preserving local neighborhood relationships. Non-Metric Multidimensional Scaling (NMDS) was then applied using the Sammon stress criterion to obtain a low-dimensional embedding for filtering, emphasizing preservation of local distances.

Mapper was run on this embedded space using overlapping bins and local clustering. The resulting graph consisted of nodes—representing local subgroups of contrast maps—and edges, which indicated shared data points across nodes.

Nodes were annotated using either RDoC domain labels (based on task-condition mappings; Supplementary Table 23) or data-driven community centroid labels (assigned via maximal spatial correlation). To assess the stability of modularity under both annotation schemes, we performed jackknife resampling across subjects. For each iteration, one subject was omitted, and Mapper was re-run on the remaining participants using identical parameters (number of bins, overlap, resolution, gain, k, and embedding dimensionality). Modularity was then computed for each resampled graph after annotating nodes with either RDoC or community labels. This produced empirical distributions of modularity values for both labeling schemes, separately for each cohort.

### 4.8 Statistical Analysis of Model Fit

To compare model performance, paired t-tests were conducted to compare the data-driven bifactor and RDoC models within each cohort for both training and cross-validation analyses. For the training analyses to compare all four model types (data-driven specific, data-driven bifactor, RDoC specific, and RDoC bifactor), we conducted repeated-measures ANOVAs separately for each fit index: robust RMSEA, robust CFI, robust TLI, and AIC. Model type was treated as a four-level within-subject factor, with each participant contributing a set of fit indices for all model types. Mauchly’s test indicated that the assumption of sphericity was violated for all indices; therefore, Greenhouse–Geisser corrections were applied to the degrees of freedom in all tests. ANOVAs were followed by Tukey-adjusted pairwise comparisons to evaluate differences between model families. All analyses were implemented in R (version 4.3.0) using the afex (version 1.5.0) and emmeans (version 1.11.2.8) packages.

For the group-level behavioral analyses in HCP and LA5c, repeated-measures ANOVAs were applied to leave-one-out distributions of AIC, RMSEA, CFI, and TLI across the four model types, followed by Tukey-adjusted pairwise comparisons. For the pooled LA5c neural analyses, the same repeated-measures ANOVA and Tukey framework was used to compare all four model families. For LA5c projection analyses comparing healthy control derived DD-b solutions with RDoC models within each diagnostic group, paired t-tests were computed separately for each fit index and component solution.

### 4.9 Correlations Between Factor Scores

We assessed the topographical stability of community centroids by computing parcel-wise Pearson correlations between centroids derived from Cohort 1 and Cohort 2. To evaluate alignment with existing cognitive ontologies, we also correlated cohort-specific centroid maps with previously published group-level activation maps from an independent dataset^7^, encompassing both data-driven and RDoC-based representations.

To assess cross-dataset consistency, we additionally correlated LA5c community centroids with HCP centroids, LA5c community centroids with LA5c RDoC classical factor maps, and LA5c RDoC classical factor maps with HCP centroids. We also computed pairwise parcel-wise correlations between diagnosis-specific LA5c general factor maps. All correlations were performed across the 347 parcels using unthresholded factor score maps. Statistical significance was evaluated using BrainSMASH^22^, which generates spatially autocorrelated null maps that preserve the intrinsic spatial structure of the data. P values were corrected for spatial autocorrelation and, where appropriate, further corrected for multiple comparisons using the Benjamini–Hochberg false discovery rate procedure. Inter-parcel distances for BrainSMASH were computed using Euclidean distances between the MNI-space centroids of each parcel. All statistical tests were two-tailed.

To examine whether the general factor primarily reflected canonical task-positive organization, we compared the HCP and pooled LA5c general factor maps with independent Neurosynth^20^ association and uniformity maps for the terms “task,” “task positive,” and “default mode.” All maps were parcellated to the same 347-region atlas and z-scored across parcels before analysis. Parcel-wise Pearson correlations were computed between each general factor and each Neurosynth map. We additionally fit linear models using paired task-like and default-mode templates to estimate the proportion of variance in each general factor explained by canonical task-positive and default-mode patterns.

### 4.10 Task Performance Correlates

We tested whether the similarity between an individual’s contrast map and the corresponding community centroid predicted cognitive task performance. Similarity was quantified using both Pearson correlation and continuous (soft) Dice coefficients. While correlation captures linear similarity, soft Dice scores account for spatial overlap weighted by magnitude. For each task contrast map, the continuous Dice similarity^48^ was computed as:

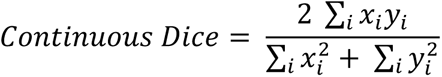

where 𝑥 is the centroid vector and 𝑦 is the individual’s contrast map. This formulation generalizes the traditional Dice coefficient to continuous, real-valued data, allowing sensitivity to both spatial alignment and intensity variation.

Associations between map–centroid similarity and behavior were estimated using linear mixed-effects models. The fixed effect was map–centroid similarity, and a random intercept for Parental ID (defined as the combination of maternal and paternal IDs) accounted for genetic relatedness. Behavioral outcomes included both raw task accuracy and an EFA-derived latent performance score (described below). We focused on the working memory and relational reasoning tasks, which are the only HCP tasks with well-defined individual-level behavioral metrics (see Supplementary Methods 1 for discussion of other tasks).

To derive a general measure of task performance, we conducted EFAs separately for the working memory and relational tasks, analyzing accuracy and median reaction time using principal axis factoring. A single latent factor was extracted, capturing shared variance among the behavioral measures. Each participant’s score on this factor was used as an index of their overall working memory/relational reasoning performance.

### 4.11 Clinical Correlates in LA5c

In LA5c, clinical relevance was evaluated by correlating alignment to data-driven community centroids with symptom measures including Hopkins Global Severity, Young Mania Rating Scale, ASRS v1.1 Screener, and Hamilton Depression Rating Scale. If an individual contributed more than one specific factor to a given community, alignment values were averaged within community before correlation with symptoms. For the RDoC comparison, each participant’s Cognitive Systems or Positive Valence Systems factor was correlated with the corresponding group-average template factor. P values were adjusted using the Benjamini–Hochberg false discovery rate procedure.

### 4.12 Sex Effects Analysis

To assess potential sex differences in task-evoked brain activation, we performed parcel-wise regressions of activation level on sex across all 24 contrasts and 347 parcels in each cohort. After false discovery rate (FDR) correction, 9.1% of tests remained significant in Cohort 1 and 2.4% in Cohort 2. Given the limited reproducibility and small effect sizes, sex was not included as a covariate in the main analyses in order to preserve model parsimony and statistical power. Future work may explore demographic influences such as sex in more depth within this framework.

## Acknowledgements

This work was supported by National Institutes of Mental Health (NIMH) grant R01MH127608 and a Maternal and Child Health Research Institute (MCHRI) Faculty Scholar Award to M.S. L.Q.U. is supported by R21HD111805 from the National Institute of Child Health and Human Development (NICHD) and U01DA050987 from the National Institute on Drug Abuse (NIDA). I.H.G. was supported by NIMH Grant R37MH101495.

## References

1. Cuthbert, B. N. & Insel, T. R. Toward new approaches to psychotic disorders: the NIMH Research Domain Criteria project. Schizophr. Bull. 36, 1061–1062 (2010).

2. Insel, T. et al. Research domain criteria (RDoC): toward a new classification framework for research on mental disorders. Am. J. Psychiatry 167, 748–751 (2010).

3. Cuthbert, B. N. & Insel, T. R. Toward the future of psychiatric diagnosis: the seven pillars of RDoC. BMC Med. 11, 126 (2013).

4. Kozak, M. J. & Cuthbert, B. N. The NIMH research domain criteria initiative: Background, issues, and pragmatics. Psychophysiology 53, 286–297 (2016).

5. Beam, E., Potts, C., Poldrack, R. A. & Etkin, A. A data-driven framework for mapping domains of human neurobiology. Nat. Neurosci. 24, 1733–1744 (2021).

6. Morawetz, C., Hemetsberger, F. J., Laird, A. R. & Kohn, N. Emotion regulation: From neural circuits to a transdiagnostic perspective. Neurosci. Biobehav. Rev. 168, 105960 (2025).

7. Quah, S. K. L. et al. A data-driven latent variable approach to validating the research domain criteria framework. Nat. Commun. 16, 830 (2025).

8. Cuthbert, B. N. Research Domain Criteria (RDoC): Progress and potential. Curr. Dir. Psychol. Sci. 31, 107–114 (2022).

9. Rubin, T. N. et al. Decoding brain activity using a large-scale probabilistic functional-anatomical atlas of human cognition. PLoS Comput. Biol. 13, e1005649 (2017).

10. Kotov, R., Krueger, R. F. & Watson, D. A paradigm shift in psychiatric classification: the Hierarchical Taxonomy Of Psychopathology (HiTOP). World Psychiatry vol. 17 Preprint at 10.1002/wps.20478 (2018).

11. Poldrack, R. A. et al. A phenome-wide examination of neural and cognitive function. Scientific Data 3, (2016).

12. Margulies, D. S. et al. Situating the default-mode network along a principal gradient of macroscale cortical organization. Proc. Natl. Acad. Sci. U. S. A. (2016) doi:10.1073/pnas.1608282113.

13. Saggar, M. et al. Towards a new approach to reveal dynamical organization of the brain using topological data analysis. Nature Communications 2018 9:1 9, 1–14 (2018).

14. Singh, G., Mémoli, F. & Carlsson, G. Topological Methods for the Analysis of High Dimensional Data Sets and 3D Object Recognition. Eurographics Symposium on Point-Based Graphics (2007) doi:10.2312/SPBG/SPBG07/091-100.

15. Bolt, T., Nomi, J. S., Yeo, B. T. T. & Uddin, L. Q. Data-Driven Extraction of a Nested Model of Human Brain Function. J. Neurosci. 37, 7263–7277 (2017).

16. Bolt, T. et al. Ontological dimensions of cognitive-neural mappings. Neuroinformatics 18, 451–463 (2020).

17. Waldman, I. D. et al. Recommendations for adjudicating among alternative structural models of psychopathology. Clin. Psychol. Sci. 11, 616–640 (2023).

18. Rodriguez, A., Reise, S. P. & Haviland, M. G. Evaluating bifactor models: Calculating and interpreting statistical indices. Psychol. Methods 21, 137–150 (2016).

19. Gordon, E. M. et al. Precision Functional Mapping of Individual Human Brains. Neuron 95, 791–807.e7 (2017).

20. Yarkoni, T., Poldrack, R. A., Nichols, T. E., Van Essen, D. C. & Wager, T. D. Large-scale automated synthesis of human functional neuroimaging data. Nat. Methods 8, 665–670 (2011).

21. Vazquez-Trejo, V., Nardos, B., Schlaggar, B. L., Fair, D. A. & Miranda-Dominguez, O. Use of connectotyping on task functional MRI data reveals dynamic network level cross talking during task performance. Front. Neurosci. 16, 951907 (2022).

22. Burt, J. B., Helmer, M., Shinn, M., Anticevic, A. & Murray, J. D. Generative modeling of brain maps with spatial autocorrelation. Neuroimage 220, 117038 (2020).

23. Shine, J. M. et al. Human cognition involves the dynamic integration of neural activity and neuromodulatory systems. Nat. Neurosci. 22, (2019).

24. Huntenburg, J. M., Bazin, P.-L. & Margulies, D. S. Large-scale gradients in human cortical organization. Trends Cogn. Sci. 22, 21–31 (2018).

25. Bolt, T. et al. A parsimonious description of global functional brain organization in three spatiotemporal patterns. Nat. Neurosci. 25, 1093–1103 (2022).

26. Bartra, O., McGuire, J. T. & Kable, J. W. The valuation system: a coordinate-based meta-analysis of BOLD fMRI experiments examining neural correlates of subjective value. Neuroimage 76, 412–427 (2013).

27. Hardwick, R. M., Rottschy, C., Miall, R. C. & Eickhoff, S. B. A quantitative meta-analysis and review of motor learning in the human brain. Neuroimage 67, 283–297 (2013).

28. de la Vega, A., Yarkoni, T., Wager, T. D. & Banich, M. T. Large-scale meta-analysis suggests low regional modularity in lateral frontal cortex. Cereb. Cortex 28, 3414–3428 (2018).

29. Egli, T. et al. Identification of two distinct working memory-related brain networks in healthy young adults. eNeuro 5, (2018).

30. Stern, Y., Gazes, Y., Razlighi, Q., Steffener, J. & Habeck, C. A task-invariant cognitive reserve network. Neuroimage 178, 36–45 (2018).

31. Laumann, T. O. et al. Functional System and Areal Organization of a Highly Sampled Individual Human Brain. Neuron 87, 657–670 (2015).

32. Braga, R. M. & Buckner, R. L. Parallel interdigitated distributed networks within the individual estimated by intrinsic functional connectivity. Neuron 95, 457–471.e5 (2017).

33. Mumford, J. A. et al. The response time paradox in functional magnetic resonance imaging analyses. *Nat*. Hum. Behav. 8, 349–360 (2024).

34. Poldrack, R. A. & Yarkoni, T. From Brain Maps to Cognitive Ontologies: Informatics and the Search for Mental Structure. Annu. Rev. Psychol. 67, 587–612 (2016).

35. Van Essen, D. C. et al. The WU-Minn Human Connectome Project: An overview. Neuroimage 80, 62–79 (2013).

36. Barch, D. M. et al. Function in the human connectome: task-fMRI and individual differences in behavior. Neuroimage 80, 169–189 (2013).

37. Glasser, M. F. et al. The minimal preprocessing pipelines for the Human Connectome Project. Neuroimage 80, 105–124 (2013).

38. Smith, S. M. et al. Resting-state fMRI in the Human Connectome Project. Neuroimage 80, 144–168 (2013).

39. Desikan, R. S. et al. An automated labeling system for subdividing the human cerebral cortex on MRI scans into gyral based regions of interest. Neuroimage 31, 968–980 (2006).

40. Stanton, N. Human Cognitive Abilities: A Survey of Factor-Analytic Studies, by J. B. Carroll, Cambridge University Press, Cambridge (1993), pp. iv + 819, ISBN 0-521-38712-4. Ergonomics 38, 1074–1074 (1995).

41. Horn, J. L. A rationale and test for the number of factors in factor analysis. Psychometrika 30, 179–185 (1965).

42. Hayton, J. C., Allen, D. G. & Scarpello, V. Factor retention decisions in exploratory factor analysis: A tutorial on parallel analysis. Organ. Res. Methods 7, 191–205 (2004).

43. Brooks, S. & Stevens, J. Applied multivariate statistics for the social sciences. Statistician 43, 219 (1994).

44. Burnham, K. P. & Anderson, D. R. Multimodel inference. Sociol. Methods Res. 33, 261–304 (2004).

45. Blondel, V. D., Guillaume, J.-L., Lambiotte, R. & Lefebvre, E. Fast unfolding of communities in large networks. J. Stat. Mech: Theory Exp. 2008, P10008 (2008).

46. Saggar, M., Shine, J. M., Liégeois, R., Dosenbach, N. U. F. & Fair, D. Precision dynamical mapping using topological data analysis reveals a hub-like transition state at rest. Nature Communications 2022 13:1 13, 1–19 (2022).

47. Duda, R. O. Pattern Classification. (John Wiley & Sons, Nashville, TN, 2022).

48. Deep Learning and Data Labeling for Medical Applications. (Springer International Publishing, Cham, Switzerland, 2016).

