## Supplementary Information for "Refining RDoC Using Individual-Level Task fMRI Factor Models Reveals Reproducible and Clinically Relevant Brain-Wide Motifs"

* These authors contributed equally

**Supplementary Figure 1: Data-driven bifactor models show superior model fit compared to RDoC-based models across training and validation cohorts, as measured by CFI and TLI.** Violin plots of comparative fit index (CFI) and Tucker–Lewis index (TLI) for confirmatory factor analysis models. (a) Model fit for data-driven bifactor (DD) and RDoC-based models trained within Cohort 1 (top) and Cohort 2 (bottom). (b) Validation of data-driven models projected onto held-out cohorts using principal components (3–7 components, 3C–7C) compared with RDoC models. Blue indicates data-driven models; red indicates RDoC-based models. Higher values indicate better fit. Data-driven models consistently outperformed RDoC-based models across all comparisons. ****p* < .001.

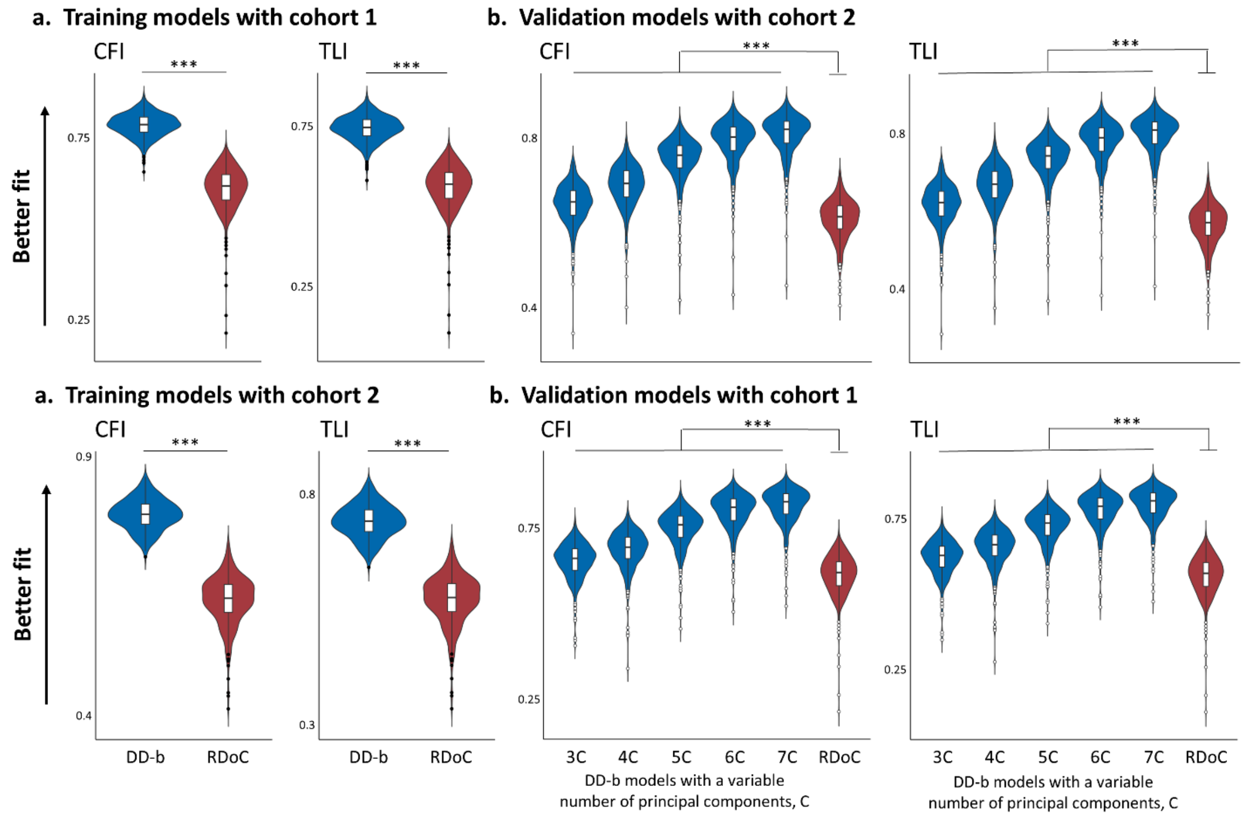

**Supplementary Figure 2: Comparison of model fit across four model types.** Training model fit comparison using (a) robust RMSEA, (b) AIC, (c) robust CFI, and (d) robust TLI. Cohorts 1 and 2 shown separately. Violin and box plots compare four models: DD (data-driven specific), DD-b (data-driven bifactor), RDoC (RDoC specific), RDoC-b (RDoC bifactor). Y-axis for (a) and (b) inverted so lower RMSEA and AIC appears higher ("better fit"). Horizontal brackets indicate Tukey-adjusted pairwise differences from a repeated-measures ANOVA within a cohort. Across all comparisons, data-driven models outperformed RDoC-based models, and bifactor model variants outperformed their corresponding specific model variants. ***$p < .001$.

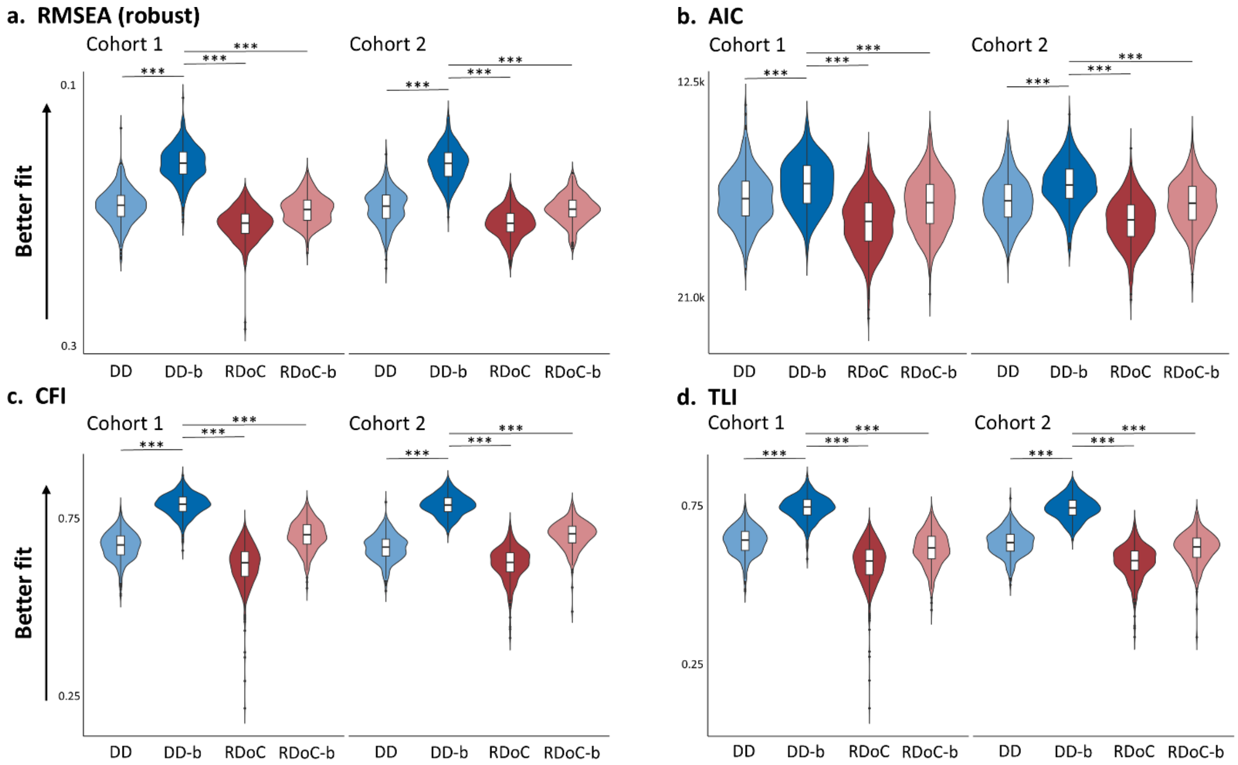


The ANOVAs indicated significant differences in fit across all model types for robust RMSEA (Cohort 1: $F(2.05, 701.85) = 2188.5$, $p < .001$; Cohort 2: $F(2.17, 606.22) = 2025.5$, $p < .001$), AIC (Cohort 1: $F(1.46, 592.74) = 794.6$, $p < .001$; Cohort 2: $F(1.39, 453.37) = 833.4$, $p < .001$), robust CFI (Cohort 1: $F(1.89, 646.92) = 2143.7$, $p < .001$; Cohort 2: $F(1.95, 544.83) = 2023.8$, $p < .001$), and robust TLI (Cohort 1: $F(1.86, 635.63) = 1724.7$, $p < .001$; Cohort 2: $F(1.91, 533.75) = 1593.5$, $p < .001$). Post hoc Tukey tests confirmed that the data-driven bifactor model consistently outperformed all other models across model fit indices in both cohorts (all ***$p < .001$). Both data-driven models outperformed both RDoC models in all instances except when measured by RMSEA in Cohort 2 only ($p = .067$). Full Tukey pairwise comparisons are provided in Supplementary Table 3.

**Supplementary Table 1: Statistics for t-test comparing data-driven validation models with 3–7 factors using principal component projection from Cohort 1 to Cohort 2 and RDoC validation models.** Metrics include AIC, RMSEA, CFI, and TLI with t-statistics, mean differences, 95% confidence intervals, and p-values.

| Components | Metric | *t* | Mean | 95% CI Lower | 95% CI Upper | *p* |
|---|---|---|---|---|---|---|
| 3 | RMSEA | -24.7 | -0.013 | -0.014 | -0.012 | <.001 |
| 4 | | -42.6 | -0.026 | -0.027 | -0.025 | <.001 |
| 5 | | -63.2 | -0.046 | -0.048 | -0.045 | <.001 |
| 6 | | -69.9 | -0.061 | -0.063 | -0.059 | <.001 |
| 7 | | -77.1 | -0.067 | -0.069 | -0.066 | <.001 |
| 3 | AIC | -18.7 | -346.7 | -383.3 | -310.2 | <.001 |
| 4 | | -37.3 | -767.7 | -808.1 | -727.2 | <.001 |
| 5 | | -58.6 | -1368.3 | -1414.2 | -1322.4 | <.001 |
| 6 | | -67.2 | -1762.8 | -1814.4 | -1711.2 | <.001 |
| 7 | | -74.3 | -1915.0 | -1965.7 | -1864.3 | <.001 |
| 3 | CFI | 18.4 | 0.035 | 0.032 | 0.039 | <.001 |
| 4 | | 38.3 | 0.079 | 0.075 | 0.084 | <.001 |
| 5 | | 62.8 | 0.142 | 0.138 | 0.147 | <.001 |
| 6 | | 74.5 | 0.184 | 0.179 | 0.188 | <.001 |
| 7 | | 84.3 | 0.200 | 0.195 | 0.204 | <.001 |
| 3 | TLI | 24.6 | 0.0526 | 0.048 | 0.057 | <.001 |
| 4 | | 43.8 | 0.1008 | 0.096 | 0.105 | <.001 |
| 5 | | 67.5 | 0.1696 | 0.165 | 0.175 | <.001 |
| 6 | | 78.7 | 0.2148 | 0.209 | 0.220 | <.001 |
| 7 | | 88.4 | 0.2325 | 0.227 | 0.238 | <.001 |

**Supplementary Table 2: Statistics for t-test comparing data-driven validation models with 3–7 factors using principal component projection from Cohort 2 to Cohort 1 and RDoC validation models.** Metrics include AIC, RMSEA, CFI, and TLI with t-statistics, mean differences, 95% confidence intervals, and p-values.

| Components | Metric | *t* | Mean | 95% CI Lower | 95% CI Upper | *p* |
|---|---|---|---|---|---|---|
| 3 | RMSEA | -27.0 | -0.014 | -0.015 | -0.013 | <.001 |
| 4 | | -39.2 | -0.024 | -0.025 | -0.023 | <.001 |
| 5 | | -68.7 | -0.044 | -0.045 | -0.042 | <.001 |
| 6 | | -78.7 | -0.061 | -0.063 | -0.060 | <.001 |
| 7 | | -85.6 | -0.067 | -0.069 | -0.066 | <.001 |
| 3 | AIC | -20.2 | -387.0 | -424.7 | -349.3 | <.001 |
| 4 | | -32.5 | -701.3 | -743.7 | -658.9 | <.001 |
| 5 | | -61.1 | -1301.4 | -1343.3 | -1259.6 | <.001 |
| 6 | | -72.2 | -1773.6 | -1821.9 | -1725.3 | <.001 |
| 7 | | -78.7 | -1928.2 | -1976.3 | -1880.0 | <.001 |
| 3 | CFI | 19.4 | 0.040 | 0.036 | 0.045 | <.001 |
| 4 | | 31.5 | 0.073 | 0.069 | 0.078 | <.001 |
| 5 | | 62.2 | 0.136 | 0.132 | 0.140 | <.001 |
| 6 | | 76.4 | 0.186 | 0.181 | 0.191 | <.001 |
| 7 | | 83.7 | 0.202 | 0.197 | 0.207 | <.001 |
| 3 | TLI | 25.0 | 0.058 | 0.054 | 0.063 | <.001 |
| 4 | | 36.2 | 0.094 | 0.089 | 0.099 | <.001 |
| 5 | | 66.5 | 0.163 | 0.158 | 0.168 | <.001 |
| 6 | | 79.9 | 0.217 | 0.212 | 0.223 | <.001 |
| 7 | | 86.9 | 0.235 | 0.230 | 0.241 | <.001 |

**Supplementary Table 3: Statistics for Tukey post hoc pairwise comparisons of model fit indices for training models.** Comparisons were conducted across the four model types: data-driven specific (DD), data-driven bifactor (DD-b), RDoC-specific (RDoC), and RDoC bifactor (RDoC-b) models in both Cohort 1 and Cohort 2. Reported values include estimated mean differences (Estimate), standard errors (SE), degrees of freedom (df), t-ratios, and adjusted p-values. All p-values were adjusted using Tukey's method for multiple comparisons within a group of four estimates. Greenhouse–Geisser corrections were applied to account for violations of sphericity.

| Contrast | Estimate | SE | df | t-ratio | *p* |
|---|---|---|---|---|---|
| DD – DD.b | 0.03265 | 0.000503 | 342 | 64.899 | <.0001 |
| DD – RDoC | -0.01412 | 0.000691 | 342 | -20.435 | <.0001 |
| DD – RDoC.b | -0.00279 | 0.000655 | 342 | -4.259 | 0.0002 |
| DD.b – RDoC | -0.04676 | 0.000693 | 342 | -67.442 | <.0001 |
| DD.b – RDoC.b | -0.03543 | 0.000659 | 342 | -53.752 | <.0001 |
| RDoC – RDoC.b | 0.01133 | 0.000372 | 342 | 30.435 | <.0001 |

| Contrast | Estimate | SE | df | t-ratio | *p* |
|---|---|---|---|---|---|
| DD – DD.b | 0.0335 | 0.000599 | 280 | 55.886 | <.0001 |
| DD – RDoC | -0.0126 | 0.00071 | 280 | -17.728 | <.0001 |
| DD – RDoC.b | -0.0017 | 0.000689 | 280 | -2.468 | 0.0673 |
| DD.b – RDoC | -0.0461 | 0.00067 | 280 | -68.715 | <.0001 |
| DD.b – RDoC.b | -0.0352 | 0.00067 | 280 | -52.498 | <.0001 |
| RDoC – RDoC.b | 0.0109 | 0.000337 | 280 | 32.311 | <.0001 |

| Contrast | Estimate | SE | df | t-ratio | *p* |
|---|---|---|---|---|---|
| DD – DD.b | -0.1155 | 0.00162 | 342 | -71.323 | <.0001 |
| DD – RDoC | 0.0549 | 0.00269 | 342 | 20.448 | <.0001 |
| DD – RDoC.b | -0.0295 | 0.00211 | 342 | -13.954 | <.0001 |
| DD.b – RDoC | 0.1704 | 0.00272 | 342 | 62.634 | <.0001 |
| DD.b – RDoC.b | 0.086 | 0.00208 | 342 | 41.261 | <.0001 |
| RDoC – RDoC.b | -0.0844 | 0.00152 | 342 | -55.651 | <.0001 |

| Contrast | Estimate | SE | df | t-ratio | p-value |
|---|---|---|---|---|---|
| DD – DD.b | -0.1205 | 0.00187 | 280 | -64.452 | <.0001 |
| DD – RDoC | 0.0465 | 0.00276 | 280 | 16.837 | <.0001 |
| DD – RDoC.b | -0.0358 | 0.00233 | 280 | -15.339 | <.0001 |
| DD.b – RDoC | 0.167 | 0.00258 | 280 | 64.607 | <.0001 |
| DD.b – RDoC.b | 0.0847 | 0.00214 | 280 | 39.595 | <.0001 |
| RDoC – RDoC.b | -0.0823 | 0.00131 | 280 | -62.936 | <.0001 |

| Contrast | Estimate | SE | df | t-ratio | *p* |
|---|---|---|---|---|---|
| DD – DD.b | -0.1044 | 0.00179 | 342 | -58.328 | <.0001 |
| DD – RDoC | 0.0707 | 0.00299 | 342 | 23.598 | <.0001 |
| DD – RDoC.b | 0.0242 | 0.00256 | 342 | 9.444 | <.0001 |
| DD.b – RDoC | 0.1751 | 0.00309 | 342 | 56.598 | <.0001 |
| DD.b – RDoC.b | 0.1286 | 0.00266 | 342 | 48.332 | <.0001 |
| RDoC – RDoC.b | -0.0464 | 0.00167 | 342 | -27.877 | <.0001 |

| Contrast | Estimate | SE | df | t-ratio | *p* |
|---|---|---|---|---|---|
| DD – DD.b | -0.1096 | 0.00209 | 280 | -52.412 | <.0001 |
| DD – RDoC | 0.0617 | 0.00308 | 280 | 20.037 | <.0001 |
| DD – RDoC.b | 0.0174 | 0.00281 | 280 | 6.202 | <.0001 |
| DD.b – RDoC | 0.1713 | 0.00295 | 280 | 58.132 | <.0001 |
| DD.b – RDoC.b | 0.127 | 0.00273 | 280 | 46.595 | <.0001 |
| RDoC – RDoC.b | -0.0443 | 0.00143 | 280 | -30.951 | <.0001 |

| Contrast | Estimate | SE | df | t-ratio | *p* |
|---|---|---|---|---|---|
| DD – DD.b | 568 | 42.2 | 405 | 13.447 | <.0001 |
| DD – RDoC | -1008 | 44 | 405 | -22.929 | <.0001 |
| DD – RDoC.b | -247 | 42.7 | 405 | -5.767 | <.0001 |
| DD.b – RDoC | -1576 | 21.1 | 405 | -74.556 | <.0001 |
| DD.b – RDoC.b | -814 | 15.7 | 405 | -51.849 | <.0001 |
| RDoC – RDoC.b | 762 | 14.7 | 405 | 51.864 | <.0001 |

| Contrast | Estimate | SE | df | t-ratio | *p* |
|---|---|---|---|---|---|
| DD – DD.b | 694 | 39.5 | 326 | 17.588 | <.0001 |
| DD – RDoC | -847 | 41.8 | 326 | -20.251 | <.0001 |
| DD – RDoC.b | -111 | 41.3 | 326 | -2.696 | 0.0369 |
| DD.b – RDoC | -1541 | 18.2 | 326 | -84.772 | <.0001 |
| DD.b – RDoC.b | -806 | 16.3 | 326 | -49.388 | <.0001 |
| RDoC – RDoC.b | 735 | 10.9 | 326 | 67.539 | <.0001 |

**Supplementary Table 4, Summary statistics for factor quality indices in the HCP data-driven task-fMRI bifactor models.** Mean, standard deviation, median, interquartile range, minimum, and maximum values are shown for omega total, omega hierarchical (Omega H), omega hierarchical subscale (Omega HS), explained common variance (ECV), item-level ECV (I-ECV), percentage of uncontaminated correlations (PUC), factor determinacy, and construct replicability (H). Results are shown separately for Cohort 1 and Cohort 2.

**Cohort 1**

| Metric | Mean | SD | Median | IQR | Min | Max |
|---|---|---|---|---|---|---|
| Omega total | 0.9708 | 0.01083 | 0.9719 | 0.01073 | 0.9072 | 0.9967 |
| Omega H (general) | 0.7396 | 0.1909 | 0.8082 | 0.1134 | 0.09335 | 0.9122 |
| Omega HS mean (specifics) | 0.03985 | 0.03422 | 0.02667 | 0.02004 | 0.009915 | 0.1847 |
| Omega HS max (specifics) | 0.1489 | 0.1939 | 0.07463 | 0.06542 | 0.01567 | 0.8897 |
| ECV (general) | 0.4676 | 0.1761 | 0.5252 | 0.116 | 0.005489 | 0.6762 |
| I-ECV (general) | 0.5141 | 0.06931 | 0.523 | 0.08566 | 0.2161 | 0.6709 |
| ECV mean (specifics) | 0.09113 | 0.03164 | 0.08032 | 0.02231 | 0.05376 | 0.244 |
| ECV max (specifics) | 0.2426 | 0.2535 | 0.151 | 0.04693 | 0.07988 | 0.989 |
| PUC | 0.7714 | 0.07288 | 0.779 | 0.1132 | 0.5616 | 0.9167 |
| Determinacy (general) | 0.9832 | 0.01214 | 0.9861 | 0.01521 | 0.9345 | 1 |
| Determinacy mean (specifics) | 0.9351 | 0.02576 | 0.9383 | 0.0314 | 0.7986 | 0.9906 |
| Determinacy min (specifics) | 0.8617 | 0.09199 | 0.8917 | 0.08507 | 0.1345 | 0.9683 |
| H (general) | 0.9955 | 0.006666 | 1 | 0.009021 | 0.9613 | 1 |
| H mean (specifics) | 0.9027 | 0.03953 | 0.9036 | 0.04887 | 0.7572 | 1 |
| H min (specifics) | 0.7862 | 0.1259 | 0.8254 | 0.1054 | 0.0183 | 1 |

**Cohort 2**

| Metric | Mean | SD | Median | IQR | Min | Max |
|---|---|---|---|---|---|---|
| Omega total | 0.9703 | 0.009223 | 0.9708 | 0.009393 | 0.9289 | 0.9943 |
| Omega H (general) | 0.7667 | 0.1561 | 0.8155 | 0.08619 | 0.1301 | 0.9124 |
| Omega HS mean (specifics) | 0.03507 | 0.02961 | 0.02496 | 0.01515 | 0.01072 | 0.2132 |
| Omega HS max (specifics) | 0.1216 | 0.157 | 0.06895 | 0.03986 | 0.02004 | 0.826 |
| ECV (general) | 0.488 | 0.15 | 0.5281 | 0.09881 | 0.008175 | 0.7019 |
| I-ECV (general) | 0.514 | 0.06481 | 0.5215 | 0.07842 | 0.2203 | 0.6593 |
| ECV mean (specifics) | 0.08676 | 0.02905 | 0.07866 | 0.02047 | 0.05353 | 0.2474 |
| ECV max (specifics) | 0.2125 | 0.2158 | 0.1463 | 0.03905 | 0.08329 | 0.9809 |
| PUC | 0.7722 | 0.06618 | 0.779 | 0.09058 | 0.5761 | 0.913 |
| Determinacy (general) | 0.9841 | 0.01187 | 0.9866 | 0.01379 | 0.9312 | 1 |
| Determinacy mean (specifics) | 0.9305 | 0.0263 | 0.9341 | 0.03174 | 0.8276 | 0.9862 |
| Determinacy min (specifics) | 0.842 | 0.113 | 0.881 | 0.1297 | 0.05204 | 0.9593 |
| H (general) | 0.9953 | 0.006985 | 1 | 0.008947 | 0.9607 | 1 |
| H mean (specifics) | 0.8936 | 0.03965 | 0.8969 | 0.05239 | 0.7772 | 1 |
| H min (specifics) | 0.7515 | 0.1511 | 0.809 | 0.195 | 0.002765 | 1 |

**Supplementary Figure 3: Permutation analysis of contrast to domain assignment in HCP RDoC factor models.** Observed RDoC model fit was compared with null distributions obtained by permuting contrast-to-domain assignments. Percentile ranks are shown for RMSEA, CFI, TLI, and AIC in Cohort 1 and Cohort 2. Across all fit indices, the prespecified assignment fell near the extreme upper tail of the permutation distribution, indicating that the reported mapping was not interchangeable with arbitrary assignments.

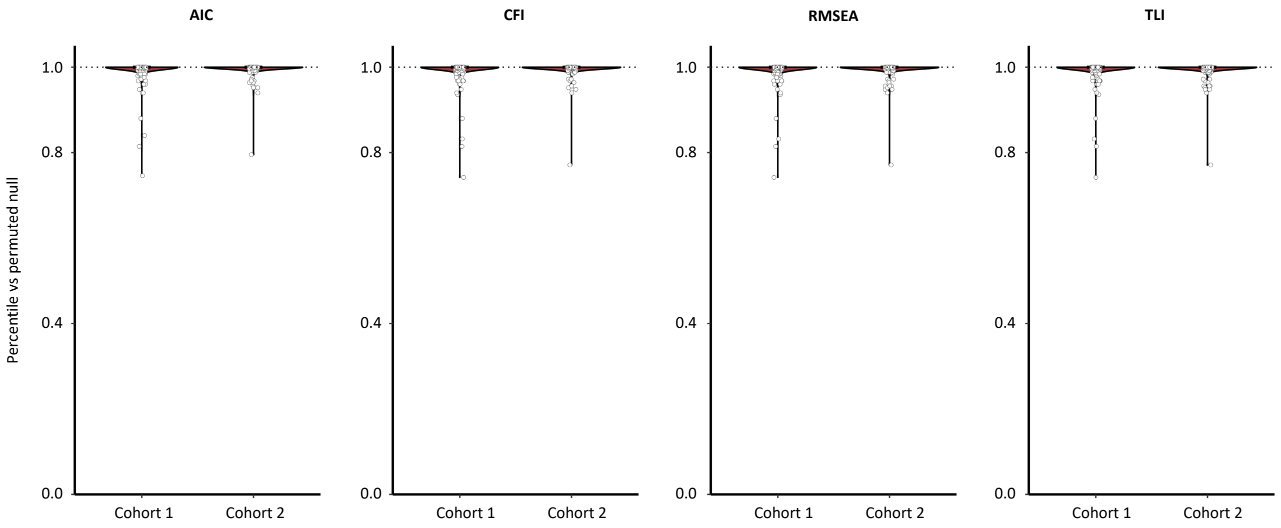

**Supplementary Table 5: Percentile ranks of observed model fit relative to permuted contrast-to-domain assignments.** Percentile rank values are presented as mean (SD) and 95% confidence intervals for Cohort 1 and Cohort 2. Values range from 0 to 1, with higher values indicating that the prespecified assignment outperformed a larger proportion of permuted assignments.

| Measure | Cohort 1 (n=411) Mean (SD) | 95% CI | Cohort 2 (n=329) Mean (SD) | 95% CI |
|---|---|---|---|---|
| RMSEA percentile | 0.996 (0.020) | 0.994–0.998 | 0.997 (0.015) | 0.995–0.999 |
| CFI percentile | 0.996 (0.020) | 0.994–0.998 | 0.997 (0.015) | 0.995–0.999 |
| TLI percentile | 0.996 (0.020) | 0.994–0.998 | 0.997 (0.015) | 0.995–0.999 |
| AIC percentile | 0.996 (0.020) | 0.994–0.998 | 0.997 (0.013) | 0.996–0.999 |

**Supplementary Table 6: Table of p-values of task-derived general factors with resting-state gradients.** Table shows parcel-wise p-values between the general factor recovered from data-driven bifactor models and the first five principal gradients of resting-state functional connectivity reported by Margulies et al. (2016). Results are presented separately for Cohort 1 and Cohort 2. "G" indicates the data-driven general factor. G1–G5 indicate resting-state gradients 1–5;

**Cohort 1**

| | **G1** | **G2** | **G3** | **G4** | **G5** |
|---|---|---|---|---|---|
| **G** | 0.46 | 0 | 0 | 0.032 | 0.46 |

**Cohort 2**

| | **G1** | **G2** | **G3** | **G4** | **G5** |
|---|---|---|---|---|---|
| **G** | 0.441 | 0 | 0 | 0.043 | 0.441 |

**Supplementary Figure 4. External meta-analytic benchmark of the general factor against canonical task positive and default mode templates** Shown are brain maps for the external Neurosynth task positive association map, the external Neurosynth default mode association map, the HCP Cohort 1 general factor map, and the pooled LA5c general factor map. All maps were parcellated to the same 347 region atlas and z-scored across parcels before comparison. Warm colors indicate relatively more positive parcel values and cool colors indicate relatively more negative parcel values. The Neurosynth association maps were selected for the main visual benchmark because they more specifically index preferential term activation relationships than uniformity maps.

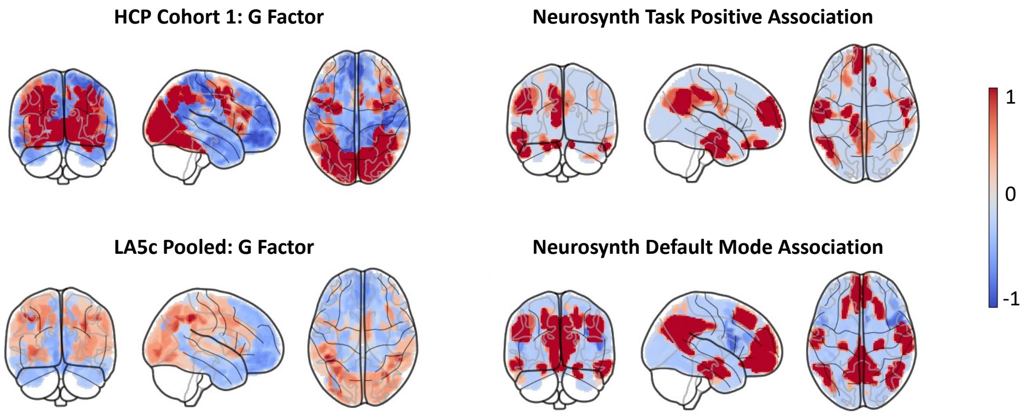

**Supplementary Table 7. External Neurosynth benchmark of the general factor.** For each dataset, parcelwise spatial correlations and regression-based variance explained are shown for four external Neurosynth template pairs: Task Association / Default Mode Association, Task Positive Association / Default Mode Association, Task Positive Uniformity / Default Mode Uniformity, and Task Uniformity / Default Mode Uniformity. r (Task) and r (DMN) are parcelwise correlations between the general-factor map and the task-like and default-mode templates, respectively. $R^2$ (Task Only) and $R^2$ (DMN Only) are the proportions of parcelwise variance explained by each template individually. $R^2$ (Full Model) is the variance explained by the two-template model G ~ "task-like template" + "default-mode template". Variance Remaining equals 1- $R^2$ . All maps were parcellated to the 347-region atlas and z-scored across parcels before analysis.

| G factor | Neurosynth comparison | r (Task) | r (DMN) | $R^2$ (Task Only) | $R^2$ (DMN Only) | $R^2$ (Full Model) | Variance Remaining |
|---|---|---|---|---|---|---|---|
| HCP Cohort 1 | Task Association / Default Mode Association | 0.506 | -0.251 | 0.256 | 0.063 | 0.289 | 0.711 |
| HCP Cohort 1 | Task Positive Association / Default Mode Association | -0.127 | -0.251 | 0.016 | 0.063 | 0.069 | 0.931 |
| HCP Cohort 1 | Task Positive Uniformity / Default Mode Uniformity | 0.283 | 0.026 | 0.08 | 0.001 | 0.108 | 0.892 |
| HCP Cohort 1 | Task Uniformity / Default Mode Uniformity | 0.509 | 0.026 | 0.259 | 0.001 | 0.323 | 0.677 |
| HCP Cohort 2 | Task Association / Default Mode Association | 0.505 | -0.257 | 0.255 | 0.066 | 0.29 | 0.71 |
| HCP Cohort 2 | Task Positive Association / Default Mode Association | -0.129 | -0.257 | 0.017 | 0.066 | 0.072 | 0.928 |
| HCP Cohort 2 | Task Positive Uniformity / Default Mode Uniformity | 0.278 | 0.021 | 0.077 | 0 | 0.106 | 0.894 |
| HCP Cohort 2 | Task Uniformity / Default Mode Uniformity | 0.505 | 0.021 | 0.255 | 0 | 0.32 | 0.68 |
| LA5c pooled | Task Association / Default Mode Association | 0.413 | -0.277 | 0.171 | 0.077 | 0.22 | 0.78 |
| LA5c pooled | Task Positive Association / Default Mode Association | -0.206 | -0.277 | 0.042 | 0.077 | 0.099 | 0.901 |
| LA5c pooled | Task Positive Uniformity / Default Mode Uniformity | 0.266 | 0.029 | 0.071 | 0.001 | 0.093 | 0.907 |
| LA5c pooled | Task Uniformity / Default Mode Uniformity | 0.493 | 0.029 | 0.243 | 0.001 | 0.301 | 0.699 |

**Supplementary Table 8. Direct cross-dataset correspondence of general factor maps.** Parcel wise correlations between the HCP Cohort 1, HCP Cohort 2, and pooled LA5c general factor maps are shown in the same 347 brain region atlas. These values provide a direct cross-dataset benchmark for correspondence of the general factor topography across the healthy HCP sample and the pooled LA5c sample. The pooled LA5c map includes healthy controls, ADHD, bipolar disorder, and schizophrenia participants.

| **G Factor Comparison** | **r** | **$R^2$** |
|---|---|---|
| HCP Cohort 1 vs HCP Cohort 2 | 0.999 | 0.998 |
| LA5c pooled vs HCP Cohort 1 | 0.583 | 0.34 |
| LA5c pooled vs HCP Cohort 2 | 0.583 | 0.34 |

**Supplementary Table 9. Spatial-autocorrelation-corrected p-values for cross-cohort correspondence of HCP community centroids.** Adjusted p-values are shown for parcel-wise correlations between community centroid maps derived from HCP Cohort 1 and Cohort 2. P-values were corrected for spatial autocorrelation using BrainSMASH.

**Adjusted P-values for correlation of community centroids from Cohort 1 and 2**

| | **C1a** | **C1b** | **C2** | **C3** | **C4** |
|---|---|---|---|---|---|
| **C1** | <.001 | <.001 | <.001 | <.001 | 0.158 |
| **C2** | <.001 | <.001 | <.001 | 0.067 | <.001 |
| **C3** | <.001 | 0.052 | 0.258 | <.001 | <.001 |
| **C4** | 0.046 | 0.713 | <.001 | <.001 | <.001 |

**Supplementary Table 10. Spatial-autocorrelation- and FDR-corrected p-values for HCP community centroid correspondence with group-level factor maps and resting-state gradients.**

**a. Correlation of Cohort 1 Community Centroids and Specific factors from Group-level t-fMRI Bifactor Models**

i. Group-level Data-Driven Specific Factors

| | Motor | Language | Attention | Reward Valuation | Theory-of-Mind | Working Memory | Social Processes | Negative valence |
|---|---|---|---|---|---|---|---|---|
| **C1** | 0 | 0.184381 | 0 | 0.190545 | 0 | 0.002667 | 0 | 0.807 |
| **C2** | 0 | 0 | 0.376615 | 0.004923 | 0.1136 | 0.679226 | 0.044 | 0.386286 |
| **C3** | 0.418133 | 0.224 | 0 | 0.1136 | 0 | 0.270667 | 0.023467 | 0.409379 |
| **C4** | 0 | 0 | 0.36992 | 0 | 0.067765 | 0.009143 | 0.386286 | 0.076444 |

ii. Group-level RDoC Specific Factors

| | Sensorimotor Systems | Positive Valence | Social Processes | Cognitive Systems | Negative Valence |
|---|---|---|---|---|---|
| **C1** | 0 | 0 | 0 | 0.076 | 0.207692 |
| **C2** | 0 | 0.114545 | 0.028889 | 0.644444 | 0.23 |
| **C3** | 0.54625 | 0 | 0 | 0.42 | 0.997 |
| **C4** | 0 | 0.196667 | 0.744211 | 0 | 0.636471 |

**b. Correlation of Cohort 2 Community Centroids and Specific factors from Group-level t-fMRI Bifactor Models**

i. Group-level Data-Driven Specific Factors

| | Motor | Language | Attention | Reward Valuation | Theory-of-Mind | Working Memory | Social Processes | Negative valence |
|---|---|---|---|---|---|---|---|---|
| **C1a** | 0 | 0.923158 | 0 | 0.696667 | 0.509565 | 0.933 | 0.457143 | 0.696667 |
| **C1b** | 0.033333 | 0.144615 | 0.53 | 0.457143 | 0.018182 | 0.008889 | 0.5312 | 0.696667 |
| **C2** | 0.005 | 0.005 | 0.692121 | 0.282667 | 0.569231 | 0.933 | 0.457143 | 0.692121 |
| **C3** | 0.718919 | 0.666667 | 0 | 0.509091 | 0 | 0.666667 | 0.448889 | 0.692121 |
| **C4** | 0.012 | 0 | 0.666667 | 0 | 0.448889 | 0.448889 | 0.64 | 0.24 |

ii. Group-level RDoC Specific Factors

| | Sensorimotor Systems | Positive Valence | Cognitive Systems | Social Processes | Negative Valence |
|---|---|---|---|---|---|
| **C1a** | 0 | 0.360417 | 0.3325 | 0.813158 | 0.645588 |
| **C1b** | 0.039286 | 0.016667 | 0.347727 | 0 | 0.813158 |
| **C2** | 0.00625 | 0.5875 | 0.419231 | 0.875 | 0.5875 |
| **C3** | 0.831818 | 0 | 0.3 | 0.845652 | 0.919 |
| **C4** | 0.015 | 0.553571 | 0.831818 | 0.3 | 0.831818 |

**ci. Correlation of Cohort 1 Community Centroids C1-4 and Principal Gradients G1-5 from Resting-state fMRI**

| | G1 | G2 | G3 | G4 | G5 |
|---|---|---|---|---|---|
| **C1** | 0.013333 | 0.07 | 0.412727 | 0.331111 | 0.44 |
| **C2** | 0.03 | 0.082857 | 0 | 0.78 | 0.925 |
| **C3** | 0.78 | 0.924211 | 0.78 | 0 | 0.78 |
| **C4** | 0.883333 | 0.675385 | 0.404 | 0.07 | 0.331111 |

**cii. Correlation of Cohort 2 Community Centroids C1a-4 and Principal Gradients G1-5 from Resting-state fMRI**

| | G1 | G2 | G3 | G4 | G5 |
|---|---|---|---|---|---|
| **C1a** | 0 | 0.551923 | 0.583929 | 0.923529 | 0.497917 |
| **C1b** | 0.9375 | 0.0375 | 0 | 0 | 0.94 |
| **C2** | 0.008333 | 0.136111 | 0 | 0.931579 | 0.94 |
| **C3** | 0.94 | 0.94 | 0.931579 | 0 | 0.923529 |
| **C4** | 0.94 | 0.705 | 0.345455 | 0.032143 | 0.345455 |

**Supplementary Methods 1: Justification for behavioral measures used in correlation analyses.** We examined all available behavioral measures in the HCP task battery to determine suitability for linking with brain–centroid similarity. Only the working memory and relational processing tasks had metrics that met two criteria: (1) sufficient inter-individual variability to detect associations and (2) a well-defined measure of task performance (as opposed to behavioral metrics). General behavior measures were not expected to correlate strongly with activation similarity to centroids, because the centroids reflect task-specific, stereotyped activation patterns.

Gambling: Outcomes tied to pre-defined reward/loss blocks rather than trial-by-trial decision-making, limiting meaningful performance quantification.

Motor: No performance metrics were available in the dataset.

Social cognition: Nearly all participants correctly identified theory-of-mind (ToM) trials, producing ceiling effects.

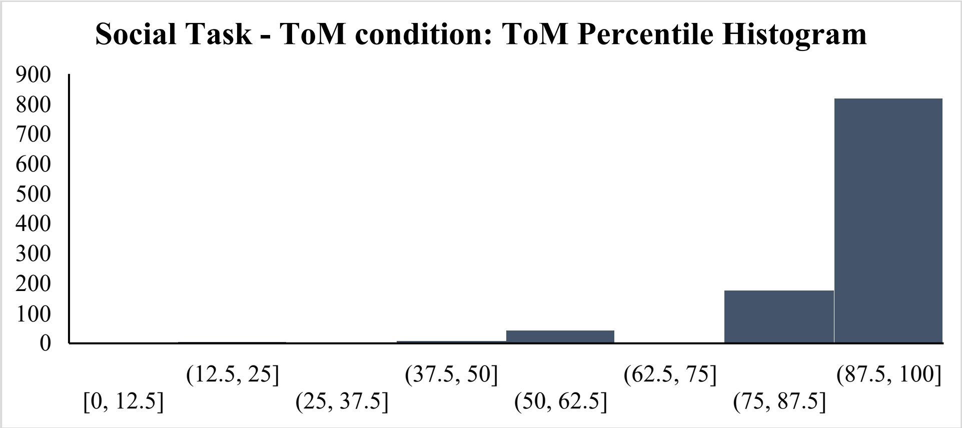


Emotion processing: Accuracy in matching emotional faces showed little variation; reaction time largely reflected general motor behavior rather than task-specific processing.

Language: Participant's accuracies were stratified and the distribution was highly skewed.

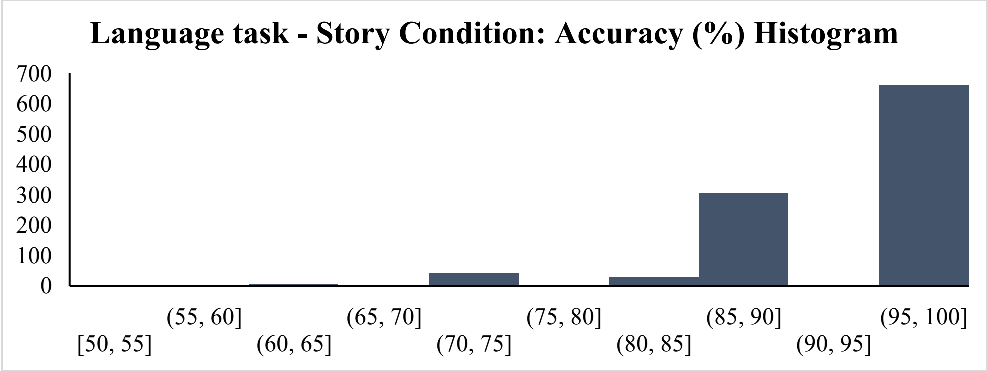


For these reasons, analyses focused on working memory and relational processing, which provided both accuracy and reaction time measures with adequate distributional properties.

**Supplementary Figure 5: Dice similarity between individual task maps and community centroids predicts behavioral performance.** Scatterplots showing associations between continuous Dice similarity of individual task maps with selected community centroids (C4 for working memory and C3 for relational reasoning) and task performance (raw accuracy and latent factor scores).

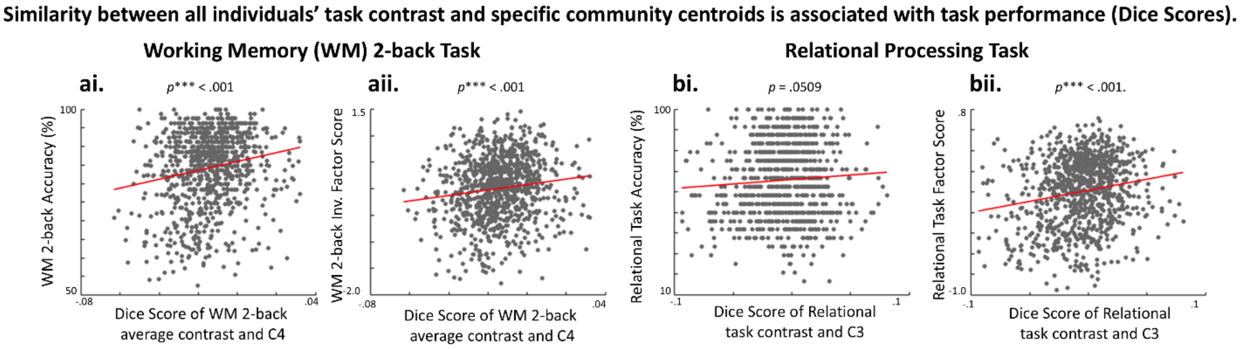

**Supplementary Table 11, Behavioral task/measure descriptions and RDoC domain assignments in HCP.** Measures drawn from the HCP behavioral battery were grouped into Cognitive Systems, Negative Valence Systems, Positive Valence Systems, Social Processes, Arousal and Regulatory Systems, and Sensorimotor Systems.

| **Task/Measure Description** | **Assigned Domain** |
|---|---|
| Short Penn CPT: sustained attention sensitivity | Cognitive Systems |
| Short Penn CPT: sustained attention specificity | Cognitive Systems |
| Short Penn CPT: true-positive median RT | Cognitive Systems |
| Flanker task: inhibitory control and attention (age-adjusted) | Cognitive Systems |
| Dimensional Change Card Sort: cognitive flexibility (age-adjusted) | Cognitive Systems |
| Relational task: relational block accuracy | Cognitive Systems |
| Relational task: relational block median RT | Cognitive Systems |
| List Sorting: working memory sequencing (age-adjusted) | Cognitive Systems |
| N-back task: 2-back accuracy | Cognitive Systems |
| N-back task: 2-back median RT | Cognitive Systems |
| Picture Sequence Memory: episodic sequence memory (age-adjusted) | Cognitive Systems |
| Penn Word Memory: total correct | Cognitive Systems |
| Penn Word Memory: correct-trial median RT | Cognitive Systems |
| Oral Reading Recognition: reading decoding (age-adjusted) | Cognitive Systems |
| Picture Vocabulary: receptive vocabulary (age-adjusted) | Cognitive Systems |
| Language task: story comprehension accuracy | Cognitive Systems |
| Penn Progressive Matrices: total correct | Cognitive Systems |
| Pattern Comparison: processing speed (age-adjusted) | Cognitive Systems |
| Penn Line Orientation: total correct | Cognitive Systems |
| Words-in-Noise: speech perception in noise | Cognitive Systems |
| Mars contrast sensitivity: final score | Cognitive Systems |

| | |
|---|---|
| Electronic visual acuity: Snellen denominator | Cognitive Systems |
| Odor Identification: olfactory identification (age-adjusted) | Cognitive Systems |
| Taste Intensity: regional taste intensity (age-adjusted) | Cognitive Systems |
| NIH Toolbox Fear-Affect: fearful affect / panic | Negative Valence Systems |
| NIH Toolbox Fear-Somatic Arousal: anxiety somatic symptoms | Negative Valence Systems |
| NIH Toolbox Sadness: sadness symptoms | Negative Valence Systems |
| NIH Toolbox Anger-Hostility: hostility / cynicism | Negative Valence Systems |
| NIH Toolbox Anger-Physical Aggression: aggression tendency | Negative Valence Systems |
| NIH Toolbox Perceived Stress: perceived uncontrollability / overload | Negative Valence Systems |
| Delay discounting AUC ($200) | Positive Valence Systems |
| Delay discounting AUC ($40,000) | Positive Valence Systems |
| NIH Toolbox Friendship: perceived companionship / affiliation | Social Processes |
| NIH Toolbox Loneliness: perceived social isolation | Social Processes |
| NIH Toolbox Emotional Support: perceived empathic support | Social Processes |
| NIH Toolbox Instrumental Support: perceived practical support | Social Processes |
| Penn Emotion Recognition (ER40): correct-trial median RT | Social Processes |
| Penn Emotion Recognition (ER40): anger identifications correct | Social Processes |
| Penn Emotion Recognition (ER40): fear identifications correct | Social Processes |
| Penn Emotion Recognition (ER40): happy identifications correct | Social Processes |
| Penn Emotion Recognition (ER40): neutral identifications correct | Social Processes |
| Penn Emotion Recognition (ER40): sad identifications correct | Social Processes |

| | |
|---|---|
| Emotion task: face-block accuracy | Social Processes |
| Emotion task: face-block median RT | Social Processes |
| Social task: Theory-of-Mind judgments (% social responses) | Social Processes |
| Social task: Theory-of-Mind judgments median RT | Social Processes |
| PSQI component 2: sleep latency | Arousal and Regulatory Systems |
| PSQI component 3: sleep duration | Arousal and Regulatory Systems |
| PSQI component 4: habitual sleep efficiency | Arousal and Regulatory Systems |
| PSQI component 7: daytime dysfunction | Arousal and Regulatory Systems |
| 2-minute walk: endurance (age-adjusted) | Sensorimotor Systems |
| 4-meter walk: gait speed | Sensorimotor Systems |
| 9-hole pegboard: manual dexterity (age-adjusted) | Sensorimotor Systems |
| Grip strength dynamometry: strength (age-adjusted) | Sensorimotor Systems |

**Supplementary Figure 6. Leave-one-out fit distributions for group-level behavioral factor models in HCP.** Violin and box plots show leave-one-out fit estimates for group-level behavioral factor models in Cohort 1 and Cohort 2. Four models are compared: data-driven specific (DD), data-driven bifactor (DD-b), RDoC specific (RDoC), and RDoC bifactor (RDoC-b). For the RDoC models, behavioral measures were assigned using fixed RDoC labels. For the data-driven models, measure-to-factor mappings were derived by EFA in the opposite cohort and applied cross-cohort. Across both cohorts, data-driven models outperformed RDoC-based models. In contrast to the task-fMRI analyses, the data-driven specific-factor model showed better fit than the data-driven bifactor model.

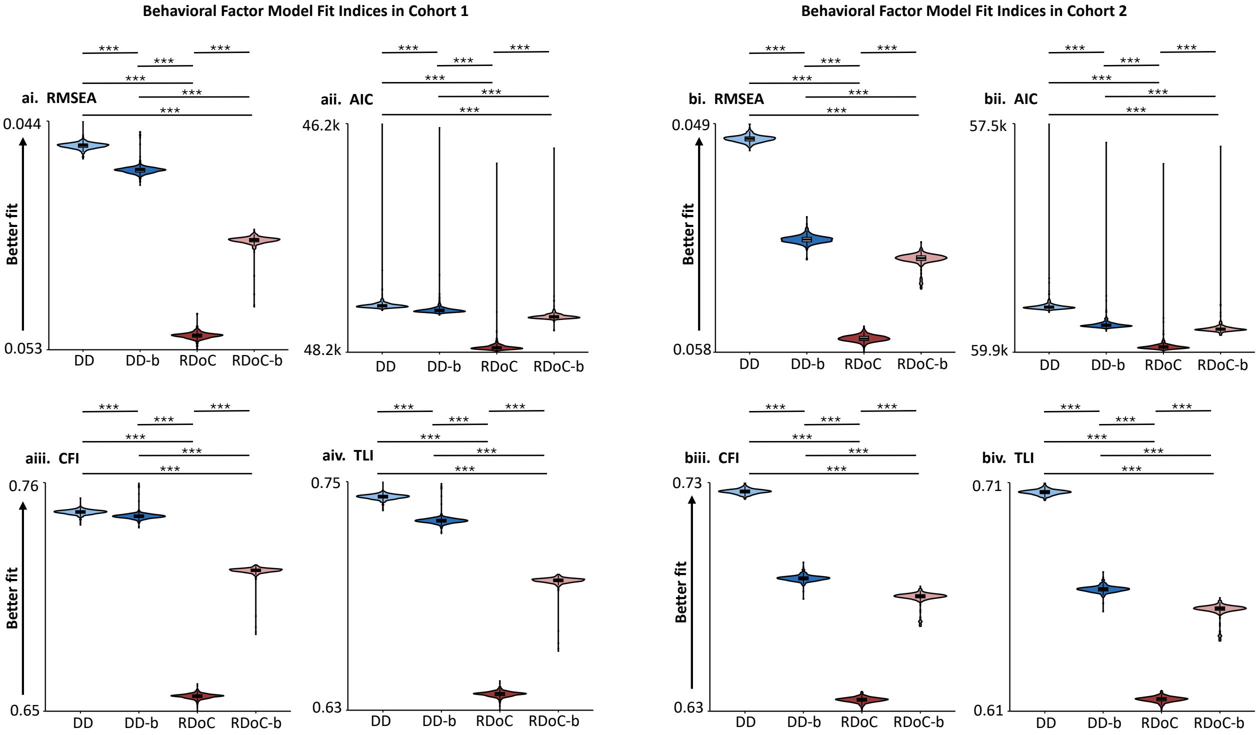

**Supplementary Table 12. Tukey-adjusted pairwise comparisons for leave-one-out behavioral model fit indices in HCP.** Comparisons were conducted across the four group-level model types: data-driven specific (DD), data-driven bifactor (DD-b), RDoC specific (RDoC), and RDoC bifactor (RDoC-b), separately within Cohort 1 and Cohort 2. Each cohort contributed one fit estimate per omitted participant iteration, obtained by refitting the model after leaving out one participant on each iteration. Reported values include estimated mean differences (Estimate), standard errors (SE), degrees of freedom (df), t-ratios, and adjusted p-values.

**Cohort 1:**

| Metric | Contrast | Estimate | SE | df | t-ratio | p |
|---|---|---|---|---|---|---|
| AIC | DD - (DD-b) | -45.970146 | 0.64637713 | 1239.0073 | -71.119698 | <.0001 |
| AIC | DD - RDoC | -423.08471 | 0.64637713 | 1239.0073 | -654.54777 | <.0001 |
| AIC | DD - (RDoC-b) | -112.2033 | 0.64637713 | 1239.0073 | -173.58797 | <.0001 |
| AIC | (DD-b) - RDoC | -377.11456 | 0.64637713 | 1239.0073 | -583.42807 | <.0001 |
| AIC | (DD-b) - (RDoC-b) | -66.233149 | 0.64637713 | 1239.0073 | -102.46827 | <.0001 |
| AIC | RDoC - (RDoC-b) | 310.881415 | 0.64637713 | 1239.0073 | 480.959797 | <.0001 |
| CFI | DD - (DD-b) | 0.00171798 | 1.18E-04 | 1239.0073 | 14.5437423 | <.0001 |
| CFI | DD - RDoC | 0.08396778 | 1.18E-04 | 1239.0073 | 710.838274 | <.0001 |
| CFI | DD - (RDoC-b) | 0.02684401 | 1.18E-04 | 1239.0073 | 227.250828 | <.0001 |
| CFI | (DD-b) - RDoC | 0.0822498 | 1.18E-04 | 1239.0073 | 696.294532 | <.0001 |
| CFI | (DD-b) - (RDoC-b) | 0.02512603 | 1.18E-04 | 1239.0073 | 212.707086 | <.0001 |
| CFI | RDoC - (RDoC-b) | -0.0571238 | 1.18E-04 | 1239.0073 | -483.58745 | <.0001 |
| RMSEA | DD - (DD-b) | -8.94E-04 | 1.02E-05 | 1239.0074 | -87.816998 | <.0001 |
| RMSEA | DD - RDoC | -0.0070653 | 1.02E-05 | 1239.0074 | -694.0188 | <.0001 |
| RMSEA | DD - (RDoC-b) | -0.0035345 | 1.02E-05 | 1239.0074 | -347.19095 | <.0001 |
| RMSEA | (DD-b) - RDoC | -0.0061713 | 1.02E-05 | 1239.0074 | -606.20181 | <.0001 |
| RMSEA | (DD-b) - (RDoC-b) | -0.0026405 | 1.02E-05 | 1239.0074 | -259.37395 | <.0001 |
| RMSEA | RDoC - (RDoC-b) | 0.00353079 | 1.02E-05 | 1239.0074 | 346.827855 | <.0001 |

| | | | | | | |
|---|---|---|---|---|---|---|
| TLI | DD - (DD-b) | 0.01052635 | 1.27E-04 | 1239.0073 | 82.6250439 | <.0001 |
| TLI | DD - RDoC | 0.08822165 | 1.27E-04 | 1239.0073 | 692.483135 | <.0001 |
| TLI | DD - (RDoC-b) | 0.03770348 | 1.27E-04 | 1239.0073 | 295.948012 | <.0001 |
| TLI | (DD-b) - RDoC | 0.0776953 | 1.27E-04 | 1239.0073 | 609.858091 | <.0001 |
| TLI | (DD-b) - (RDoC-b) | 0.02717713 | 1.27E-04 | 1239.0073 | 213.322969 | <.0001 |
| TLI | RDoC - (RDoC-b) | -0.0505182 | 1.27E-04 | 1239.0073 | -396.53512 | <.0001 |

**Cohort 2:**

| Metric | Contrast | Estimate | SE | df | t-ratio | *p* |
|---|---|---|---|---|---|---|
| AIC | DD - (DD-b) | -157.76269 | 0.48097208 | 990.009146 | -328.008 | <.0001 |
| AIC | DD - RDoC | -346.68417 | 0.48097208 | 990.009146 | -720.79894 | <.0001 |
| AIC | DD - (RDoC-b) | -192.46393 | 0.48097208 | 990.009146 | -400.15613 | <.0001 |
| AIC | (DD-b) - RDoC | -188.92147 | 0.48097208 | 990.009146 | -392.79094 | <.0001 |
| AIC | (DD-b) - (RDoC-b) | -34.701237 | 0.48097208 | 990.009146 | -72.148132 | <.0001 |
| AIC | RDoC - (RDoC-b) | 154.220238 | 0.48097208 | 990.009146 | 320.642808 | <.0001 |
| CFI | DD - (DD-b) | 0.04423966 | 1.36E-04 | 990.009147 | 325.832981 | <.0001 |
| CFI | DD - RDoC | 0.10616011 | 1.36E-04 | 990.009147 | 781.88805 | <.0001 |
| CFI | DD - (RDoC-b) | 0.05401239 | 1.36E-04 | 990.009147 | 397.810815 | <.0001 |
| CFI | (DD-b) - RDoC | 0.06192044 | 1.36E-04 | 990.009147 | 456.055069 | <.0001 |
| CFI | (DD-b) - (RDoC-b) | 0.00977272 | 1.36E-04 | 990.009147 | 71.9778343 | <.0001 |
| CFI | RDoC - (RDoC-b) | -0.0521477 | 1.36E-04 | 990.009147 | -384.07723 | <.0001 |
| RMSEA | DD - (DD-b) | -0.0041619 | 1.12E-05 | 990.009147 | -370.80331 | <.0001 |
| RMSEA | DD - RDoC | -0.0082571 | 1.12E-05 | 990.009147 | -735.65976 | <.0001 |
| RMSEA | DD - (RDoC-b) | -0.0049823 | 1.12E-05 | 990.009147 | -443.89603 | <.0001 |

| | | | | | | |
|---|---|---|---|---|---|---|
| RMSEA | (DD-b) - RDoC | -0.0040952 | 1.12E-05 | 990.009147 | -364.85645 | <.0001 |
| RMSEA | (DD-b) - (RDoC-b) | -8.20E-04 | 1.12E-05 | 990.009147 | -73.092728 | <.0001 |
| RMSEA | RDoC - (RDoC-b) | 0.00327479 | 1.12E-05 | 990.009147 | 291.76372 | <.0001 |
| TLI | DD - (DD-b) | 0.04961077 | 1.47E-04 | 990.009146 | 338.42446 | <.0001 |
| TLI | DD - RDoC | 0.10584055 | 1.47E-04 | 990.009146 | 722.001161 | <.0001 |
| TLI | DD - (RDoC-b) | 0.06018126 | 1.47E-04 | 990.009146 | 410.532078 | <.0001 |
| TLI | (DD-b) - RDoC | 0.05622978 | 1.47E-04 | 990.009146 | 383.576701 | <.0001 |
| TLI | (DD-b) - (RDoC-b) | 0.0105705 | 1.47E-04 | 990.009146 | 72.1076177 | <.0001 |
| TLI | RDoC - (RDoC-b) | -0.0456593 | 1.47E-04 | 990.009146 | -311.46908 | <.0001 |

**Supplementary Table 13. Summary statistics for factor quality indices in the HCP data-driven behavioral bifactor models.** Mean, standard deviation, median, interquartile range, minimum, and maximum values are shown for omega total, omega hierarchical (Omega H), omega hierarchical subscale (Omega HS), explained common variance (ECV), item-level ECV (I-ECV), percentage of uncontaminated correlations (PUC), factor determinacy, and construct replicability (H). Statistics were computed across leave-one-out refits and are shown separately for Cohort 1 and Cohort 2.

Cohort 1:

| Metric | Mean | SD | Median | IQR | Min | Max |
|---|---|---|---|---|---|---|
| Omega total | 0.5908 | 0.006727 | 0.5902 | 0.001005 | 0.5741 | 0.6497 |
| Omega H (general) | 0.06281 | 0.045 | 0.05724 | 0.001659 | 0.05128 | 0.4488 |
| Omega HS mean (specifics) | 0.08801 | 0.006401 | 0.0888 | 0.000308 | 0.03349 | 0.08965 |
| Omega HS max (specifics) | 0.3835 | 0.03269 | 0.3876 | 0.001208 | 0.1094 | 0.3924 |
| ECV (general) | 0.2824 | 0.006309 | 0.2817 | 0.001266 | 0.273 | 0.335 |
| I-ECV (general) | 0.2402 | 0.03295 | 0.2363 | 0.000962 | 0.2187 | 0.511 |
| ECV mean (specifics) | 0.1196 | 0.001051 | 0.1197 | 0.000211 | 0.1108 | 0.1212 |
| ECV max (specifics) | 0.2564 | 0.009464 | 0.2577 | 0.001188 | 0.1772 | 0.2613 |
| PUC | 0.7924 | 0 | 0.7924 | 0 | 0.7924 | 0.7924 |
| Determinacy (general) | 0.9182 | 0.000902 | 0.9182 | 0.000667 | 0.9128 | 0.9223 |
| Determinacy mean (specifics) | 0.891 | 0.005247 | 0.8916 | 0.001529 | 0.8502 | 0.9 |
| Determinacy min (specifics) | 0.7522 | 0.004929 | 0.7526 | 0.003214 | 0.7209 | 0.7669 |
| H (general) | 0.9927 | 0.02137 | 0.9999 | 5.48E-06 | 0.8896 | 1 |
| H mean (specifics) | 0.8341 | 0.01065 | 0.8355 | 0.002911 | 0.7511 | 0.8513 |
| H min (specifics) | 0.6677 | 0.01738 | 0.6698 | 0.005669 | 0.5272 | 0.6909 |

Cohort 2:

| Metric | Mean | SD | Median | IQR | Min | Max |
|---|---|---|---|---|---|---|
| Omega total | 0.978 | 0.00664 | 0.98 | 0.002536 | 0.9358 | 0.9835 |
| Omega H (general) | 0.005167 | 0.00748 | 0.003608 | 0.000557 | 0.002692 | 0.05971 |
| Omega HS mean (specifics) | 0.09729 | 0.001331 | 0.09763 | 0.000297 | 0.08761 | 0.09807 |
| Omega HS max (specifics) | 0.9468 | 0.01831 | 0.9518 | 0.005935 | 0.8235 | 0.9611 |
| ECV (general) | 0.002197 | 0.000772 | 0.001983 | 0.000246 | 0.001625 | 0.007593 |
| I-ECV (general) | 0.3212 | 0.0297 | 0.3168 | 0.01203 | 0.2929 | 0.4862 |

| | | | | | | |
|---|---|---|---|---|---|---|
| ECV mean (specifics) | 0.09978 | 7.72E-05 | 0.0998 | 2.46E-05 | 0.09924 | 0.09984 |
| ECV max (specifics) | 0.9893 | 0.00354 | 0.9902 | 0.002038 | 0.9587 | 0.9927 |
| PUC | 0.877 | 0 | 0.877 | 0 | 0.877 | 0.877 |
| Determinacy (general) | 0.9022 | 0.004598 | 0.903 | 0.000725 | 0.8748 | 0.9081 |
| Determinacy mean (specifics) | 0.8813 | 0.03951 | 0.893 | 0.06804 | 0.7627 | 0.9228 |
| Determinacy min (specifics) | 0.5921 | 0.1973 | 0.7367 | 0.3543 | 0.1612 | 0.7704 |
| H (general) | 0.9999 | 4.83E-05 | 0.9999 | 1.75E-05 | 0.9997 | 1 |
| H mean (specifics) | 0.814 | 0.05472 | 0.8236 | 0.09127 | 0.6698 | 0.8761 |
| H min (specifics) | 0.3911 | 0.2084 | 0.5427 | 0.4106 | 0.02612 | 0.596 |

**Supplementary Figure 7. Comparison of model fit across four model types in the pooled LA5c sample.** Internal LA5c model fit comparison using (a) RMSEA, (b) AIC, (c) CFI, and (d) TLI. Violin and box plots compare four models: DD, DD-b (data-driven bifactor), RDoC, and RDoC-b (RDoC bifactor). Y-axis for RMSEA and AIC inverted so lower values appear higher ("better fit"). Horizontal brackets indicate Tukey-adjusted pairwise differences from GG-corrected repeated-measures ANOVAs. Across all comparisons, the DD-b model showed the best fit, followed by DD, RDoC-b, and RDoC. ***p < .001.

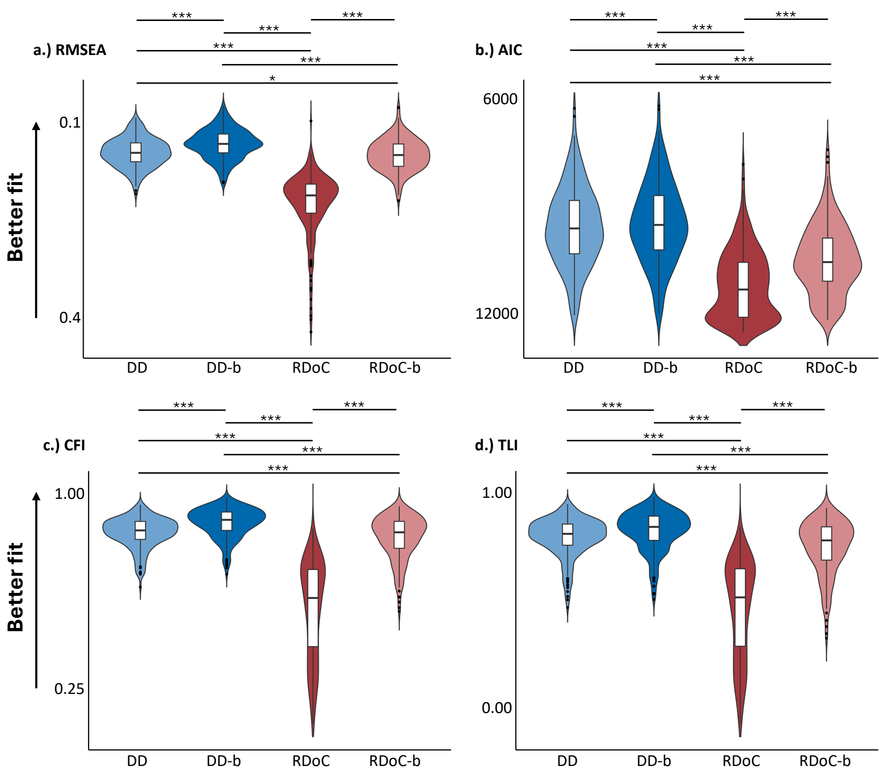


The GG-corrected repeated-measures ANOVAs indicated significant differences in fit across the four model types for robust RMSEA, $F(1.42, 297.03) = 541.6$, $p < .001$, AIC, $F(1.41, 293.80) = 499.6$, $p < .001$, robust CFI, $F(1.18, 246.76) = 646.6$, $p < .001$, and robust TLI, $F(1.23, 257.56) = 521.3$, $p < .001$. Post hoc Tukey tests showed a consistent ordering across all fit indices, with the data-driven bifactor model showing the best fit, followed by the data-driven specific model, the RDoC bifactor model, and the RDoC specific model. The data-driven bifactor model significantly outperformed all other models across RMSEA, AIC, CFI, and TLI (all $p < .001$). Both data-driven models outperformed both RDoC-based models across all fit indices, although the DD versus RDoC-b comparison for RMSEA was comparatively modest ($p = .037$). Within both the data-driven and RDoC model families, bifactor variants outperformed their corresponding specific models across all fit indices (all $p < .001$).

**Supplementary Table 14: Statistics for t-tests comparing data-driven bifactor (DD-b) and RDoC models in the LA5c sample.** Metrics include RMSEA, AIC, CFI, and TLI with t-statistics, mean differences, 95% confidence intervals, and p-values. Results are shown separately for healthy controls, ADHD, bipolar disorder, and schizophrenia.

**HC:**

| Components | Metric | *t* | Mean | 95% CI Lower | 95% CI Upper | *p* |
|---|---|---|---|---|---|---|
| 3 | RMSEA | -18.1 | -0.081 | -0.09 | -0.072 | <.001 |
| 4 | | -21.2 | -0.098 | -0.107 | -0.089 | <.001 |
| 5 | | -22.6 | -0.106 | -0.115 | -0.097 | <.001 |
| 6 | | -24.2 | -0.112 | -0.121 | -0.103 | <.001 |
| 7 | | -24.6 | -0.117 | -0.127 | -0.108 | <.001 |
| 3 | AIC | -14.1 | -708.8 | -808.6 | -609.1 | <.001 |
| 4 | | -15.9 | -812.8 | -914.1 | -711.5 | <.001 |
| 5 | | -16.7 | -861.7 | -964 | -759.4 | <.001 |
| 6 | | -17.5 | -896.7 | -998.4 | -795 | <.001 |
| 7 | | -17.7 | -921.7 | -1024.6 | -818.8 | <.001 |
| 3 | CFI | 15.1 | 0.248 | 0.216 | 0.281 | <.001 |
| 4 | | 16.9 | 0.286 | 0.252 | 0.319 | <.001 |
| 5 | | 17.5 | 0.303 | 0.269 | 0.337 | <.001 |
| 6 | | 18.2 | 0.316 | 0.282 | 0.351 | <.001 |
| 7 | | 18.6 | 0.325 | 0.29 | 0.36 | <.001 |
| 3 | TLI | 15.2 | 0.306 | 0.266 | 0.346 | <.001 |
| 4 | | 17 | 0.351 | 0.31 | 0.392 | <.001 |
| 5 | | 17.6 | 0.372 | 0.33 | 0.414 | <.001 |
| 6 | | 18.3 | 0.387 | 0.346 | 0.429 | <.001 |
| 7 | | 18.6 | 0.398 | 0.356 | 0.44 | <.001 |

**ADHD:**

| Components | Metric | *t* | Mean | 95% CI Lower | 95% CI Upper | *p* |
|---|---|---|---|---|---|---|

| | | | | | | |
|---|---|---|---|---|---|---|
| 3 | RMSEA | -9.9 | -0.073 | -0.088 | -0.058 | <.001 |
| 4 | | -11.2 | -0.086 | -0.102 | -0.071 | <.001 |
| 5 | | -11.9 | -0.095 | -0.111 | -0.079 | <.001 |
| 6 | | -12.9 | -0.101 | -0.117 | -0.085 | <.001 |
| 7 | | -13.4 | -0.105 | -0.121 | -0.089 | <.001 |
| 3 | AIC | -8 | -609.4 | -763.7 | -455 | <.001 |
| 4 | | -9 | -690.8 | -846.7 | -534.8 | <.001 |
| 5 | | -9.5 | -741.9 | -900.2 | -583.6 | <.001 |
| 6 | | -10.1 | -773.4 | -929.3 | -617.5 | <.001 |
| 7 | | -10.3 | -797.3 | -953.6 | -641.1 | <.001 |
| 3 | CFI | 9.3 | 0.225 | 0.176 | 0.274 | <.001 |
| 4 | | 10.6 | 0.257 | 0.208 | 0.306 | <.001 |
| 5 | | 11.2 | 0.278 | 0.228 | 0.329 | <.001 |
| 6 | | 11.9 | 0.291 | 0.241 | 0.34 | <.001 |
| 7 | | 12.3 | 0.3 | 0.25 | 0.349 | <.001 |
| 3 | TLI | 9.5 | 0.278 | 0.219 | 0.338 | <.001 |
| 4 | | 10.8 | 0.316 | 0.257 | 0.376 | <.001 |
| 5 | | 11.3 | 0.342 | 0.28 | 0.403 | <.001 |
| 6 | | 12 | 0.357 | 0.296 | 0.417 | <.001 |
| 7 | | 12.4 | 0.368 | 0.307 | 0.428 | <.001 |

**Bipolar:**

| **Components** | **Metric** | ***t*** | **Mean** | **95% CI Lower** | **95% CI Upper** | ***p*** |
|---|---|---|---|---|---|---|
| 3 | RMSEA | -10 | -0.074 | -0.089 | -0.059 | <.001 |
| 4 | | -11.7 | -0.087 | -0.102 | -0.072 | <.001 |
| 5 | | -13.2 | -0.096 | -0.111 | -0.081 | <.001 |
| 6 | | -13.8 | -0.101 | -0.115 | -0.086 | <.001 |
| 7 | | -13.9 | -0.105 | -0.12 | -0.09 | <.001 |
| 3 | | -7.6 | -625.1 | -790.4 | -459.8 | <.001 |
| 4 | | -8.7 | -708.4 | -873.1 | -543.7 | <.001 |

| | | | | | | |
|---|---|---|---|---|---|---|
| 5 | AIC | -9.4 | -761 | -924.1 | -598 | <.001 |
| 6 | | -9.7 | -787.3 | -950.3 | -624.3 | <.001 |
| 7 | | -9.9 | -811.4 | -976.9 | -645.8 | <.001 |
| 3 | | 10.3 | 0.233 | 0.187 | 0.278 | <.001 |
| 4 | | 12.2 | 0.267 | 0.223 | 0.312 | <.001 |
| 5 | CFI | 13.5 | 0.291 | 0.247 | 0.334 | <.001 |
| 6 | | 13.8 | 0.302 | 0.258 | 0.346 | <.001 |
| 7 | | 13.9 | 0.311 | 0.266 | 0.356 | <.001 |
| 3 | | 10.4 | 0.287 | 0.232 | 0.343 | <.001 |
| 4 | | 12.4 | 0.329 | 0.276 | 0.383 | <.001 |
| 5 | TLI | 13.6 | 0.357 | 0.304 | 0.41 | <.001 |
| 6 | | 13.9 | 0.37 | 0.317 | 0.424 | <.001 |
| 7 | | 14 | 0.382 | 0.327 | 0.437 | <.001 |

**Schizophrenia:**

| **Components** | **Metric** | ***t*** | **Mean** | **95% CI Lower** | **95% CI Upper** | ***p*** |
|---|---|---|---|---|---|---|
| 3 | | -9.9 | -0.07 | -0.084 | -0.056 | <.001 |
| 4 | | -11.1 | -0.081 | -0.095 | -0.066 | <.001 |
| 5 | RMSEA | -13 | -0.092 | -0.106 | -0.078 | <.001 |
| 6 | | -13.7 | -0.097 | -0.111 | -0.083 | <.001 |
| 7 | | -14.1 | -0.101 | -0.115 | -0.086 | <.001 |
| 3 | | -7.5 | -595.4 | -755.9 | -434.9 | <.001 |
| 4 | | -8.2 | -665.8 | -828.9 | -502.8 | <.001 |
| 5 | AIC | -9.2 | -736.8 | -898.3 | -575.3 | <.001 |
| 6 | | -9.6 | -765.6 | -926.7 | -604.6 | <.001 |
| 7 | | -9.8 | -787.2 | -948.9 | -625.4 | <.001 |
| 3 | | 11 | 0.271 | 0.222 | 0.321 | <.001 |
| 4 | | 12.2 | 0.303 | 0.253 | 0.353 | <.001 |
| 5 | CFI | 14.3 | 0.341 | 0.293 | 0.389 | <.001 |
| 6 | | 15.1 | 0.357 | 0.309 | 0.404 | <.001 |
| 7 | | 15.5 | 0.367 | 0.319 | 0.415 | <.001 |

| | | | | | | |
|---|---|---|---|---|---|---|
| 3 | | 11.2 | 0.336 | 0.276 | 0.396 | <.001 |
| 4 | | 12.4 | 0.374 | 0.313 | 0.434 | <.001 |
| 5 | TLI | 14.4 | 0.419 | 0.361 | 0.478 | <.001 |
| 6 | | 15.3 | 0.438 | 0.38 | 0.496 | <.001 |
| 7 | | 15.6 | 0.45 | 0.392 | 0.508 | <.001 |

**Supplementary Table 15. Summary statistics for factor quality indices in the LA5c data-driven task-fMRI bifactor models.** Mean, standard deviation, median, interquartile range, minimum, and maximum values are shown for omega total, omega hierarchical (Omega H), omega hierarchical subscale (Omega HS), explained common variance (ECV), item-level ECV (I-ECV), percentage of uncontaminated correlations (PUC), factor determinacy, and construct replicability (H). Statistics were computed across individual-level LA5c task-fMRI data-driven bifactor models.

| Metric | Mean | SD | Median | IQR | Min | Max |
|---|---|---|---|---|---|---|
| Omega total | 0.9191 | 0.05914 | 0.9316 | 0.06567 | 0.6704 | 0.9978 |
| Omega H (general) | 0.3302 | 0.2586 | 0.3103 | 0.4817 | 9.40E-07 | 0.9635 |
| Omega HS mean (specifics) | 0.1752 | 0.1089 | 0.1621 | 0.1487 | 0.008566 | 0.4796 |
| Omega HS max (specifics) | 0.4525 | 0.2911 | 0.4194 | 0.552 | 0.01878 | 0.9837 |
| ECV (general) | 0.2618 | 0.1774 | 0.258 | 0.253 | 0.0009781 | 0.9835 |
| I-ECV (general) | 0.3219 | 0.1443 | 0.311 | 0.2182 | 0.03728 | 0.7164 |
| ECV mean (specifics) | 0.2131 | 0.08726 | 0.1973 | 0.1021 | 0.004125 | 0.4887 |
| ECV max (specifics) | 0.4551 | 0.2644 | 0.3571 | 0.3417 | 0.01252 | 0.9938 |
| PUC | 0.6585 | 0.1317 | 0.6667 | 0.21 | 0.4091 | 0.8889 |
| Determinacy (general) | 0.9099 | 0.06031 | 0.9143 | 0.1021 | 0.7456 | 1 |
| Determinacy mean (specifics) | 0.9052 | 0.05436 | 0.9123 | 0.07495 | 0.7234 | 0.999 |
| Determinacy min (specifics) | 0.8099 | 0.1353 | 0.8279 | 0.136 | 0.008632 | 0.9968 |
| H (general) | 0.9688 | 0.0593 | 1 | 0.04144 | 0.7023 | 1 |
| H mean (specifics) | 0.8825 | 0.07477 | 0.8935 | 0.1085 | 0.6462 | 1 |
| H min (specifics) | 0.7497 | 0.1535 | 0.7687 | 0.2087 | 0.08294 | 1 |

**Supplementary Table 16, Behavioral task/measure descriptions and RDoC domain assignments in LA5c.** Measures drawn from the LA5c behavioral battery were grouped into Positive Valence Systems, Cognitive Systems, Social Processes, and Sensorimotor Systems.

| Task/Measure Description | Assigned Domain |
|---|---|
| BART: total adjusted pumps | Positive Valence Systems |
| BART: mean blue pumps after explosion | Positive Valence Systems |
| Delay discounting task: log total k | Positive Valence Systems |
| Chapman Physical Anhedonia: total score | Positive Valence Systems |
| TCI Novelty Seeking | Positive Valence Systems |
| TCI Persistence | Positive Valence Systems |
| ANT: conflict RT effect | Cognitive Systems |
| ANT: mean neutral RT | Cognitive Systems |
| Color Trails Test: time 2 | Cognitive Systems |
| Short Penn CPT: hits | Cognitive Systems |
| Short Penn CPT: false alarms | Cognitive Systems |
| CVLT: total correct | Cognitive Systems |
| D-KEFS: total errors | Cognitive Systems |
| SCAP: maximum capacity | Cognitive Systems |
| Task switching: short cost | Cognitive Systems |
| WAIS Letter-Number Sequencing: total raw | Cognitive Systems |
| WAIS Matrix Reasoning: total raw | Cognitive Systems |
| WMS Visual Reproduction II Delayed Recall: total raw | Cognitive Systems |
| Chapman Social Anhedonia: total score | Social Processes |
| Eysenck Extraversion score | Social Processes |
| TCI Reward Dependence | Social Processes |
| Stop-signal task: SSRT (quantile) | Sensorimotor Systems |
| Stop-signal task: percent inhibition | Sensorimotor Systems |
| Stop-signal task: mean RT | Sensorimotor Systems |
| Stop-signal task: direction errors | Sensorimotor Systems |

**Supplementary Figure 8. Leave-one-out fit distributions for group-level behavioral factor models in LA5c.** Violin and box plots show leave-one-out fit estimates for group-level behavioral factor models in the pooled LA5c sample. Four models are compared: data-driven specific (DD), data-driven bifactor (DD-b), RDoC specific (RDoC), and RDoC bifactor (RDoC-b). For the RDoC models, behavioral measures were assigned using fixed RDoC labels. For the data-driven models, measure-to-factor mappings were derived by EFA in the healthy-control subgroup and then applied to the pooled clinical sample. The data-driven bifactor model showed the best fit, followed by RDoC-b, DD, and RDoC.

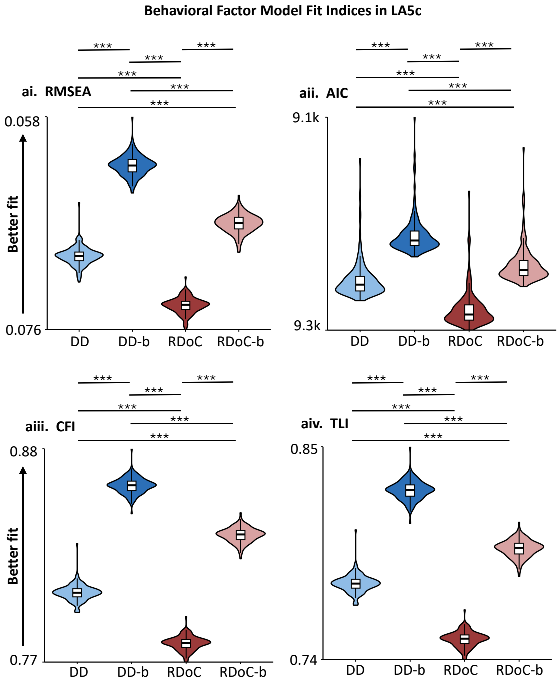

**Supplementary Table 17. Tukey-adjusted pairwise comparisons for leave-one-out behavioral model fit indices in pooled clinical sample of LA5c.** Comparisons were conducted across the four group-level model types: data-driven specific (DD), data-driven bifactor (DD-b), RDoC specific (RDoC), and RDoC bifactor (RDoC-b). There was one fit estimate per omitted participant iteration, obtained by refitting the model after leaving out one participant on each iteration. Reported values include estimated mean differences (Estimate), standard errors (SE), degrees of freedom (df), t-ratios, and adjusted p-values.

| Metric | Contrast | Estimate | SE | df | t-ratio | p |
|---|---|---|---|---|---|---|
| AIC | DD - (DD-b) | 36.1958927 | 0.22374058 | 429.021272 | 161.776164 | <.0001 |
| AIC | DD - RDoC | -23.888721 | 0.22374058 | 429.021272 | -106.76973 | <.0001 |
| AIC | DD - (RDoC-b) | 12.3708885 | 0.22374058 | 429.021272 | 55.2912151 | <.0001 |
| AIC | (DD-b) - RDoC | -60.084614 | 0.22374058 | 429.021272 | -268.54589 | <.0001 |
| AIC | (DD-b) - (RDoC-b) | -23.825004 | 0.22374058 | 429.021272 | -106.48495 | <.0001 |
| AIC | RDoC - (RDoC-b) | 36.2596098 | 0.22374058 | 429.021272 | 162.060946 | <.0001 |
| CFI | DD - (DD-b) | -0.0527445 | 1.92E-04 | 429.021269 | -274.33712 | <.0001 |
| CFI | DD - RDoC | 0.02493666 | 1.92E-04 | 429.021269 | 129.70181 | <.0001 |
| CFI | DD - (RDoC-b) | -0.0284339 | 1.92E-04 | 429.021269 | -147.89191 | <.0001 |
| CFI | (DD-b) - RDoC | 0.07768112 | 1.92E-04 | 429.021269 | 404.038928 | <.0001 |
| CFI | (DD-b) - (RDoC-b) | 0.02431054 | 1.92E-04 | 429.021269 | 126.445207 | <.0001 |
| CFI | RDoC - (RDoC-b) | -0.0533706 | 1.92E-04 | 429.021269 | -277.59372 | <.0001 |
| RMSEA | DD - (DD-b) | 0.00797703 | 3.92E-05 | 429.021277 | 203.563815 | <.0001 |
| RMSEA | DD - RDoC | -0.0043336 | 3.92E-05 | 429.021277 | -110.58872 | <.0001 |
| RMSEA | DD - (RDoC-b) | 0.0028879 | 3.92E-05 | 429.021277 | 73.6955264 | <.0001 |
| RMSEA | (DD-b) - RDoC | -0.0123107 | 3.92E-05 | 429.021277 | -314.15254 | <.0001 |
| RMSEA | (DD-b) - (RDoC-b) | -0.0050891 | 3.92E-05 | 429.021277 | -129.86829 | <.0001 |
| RMSEA | RDoC - (RDoC-b) | 0.00722152 | 3.92E-05 | 429.021277 | 184.284249 | <.0001 |

| | | | | | | |
|---|---|---|---|---|---|---|
| TLI | DD - (DD-b) | -0.0467803 | 2.25E-04 | 429.021276 | -208.06041 | <.0001 |
| TLI | DD - RDoC | 0.0278104 | 2.25E-04 | 429.021276 | 123.689789 | <.0001 |
| TLI | DD - (RDoC-b) | -0.0176076 | 2.25E-04 | 429.021276 | -78.311874 | <.0001 |
| TLI | (DD-b) - RDoC | 0.07459069 | 2.25E-04 | 429.021276 | 331.750196 | <.0001 |
| TLI | (DD-b) - (RDoC-b) | 0.02917265 | 2.25E-04 | 429.021276 | 129.748533 | <.0001 |
| TLI | RDoC - (RDoC-b) | -0.045418 | 2.25E-04 | 429.021276 | -202.00166 | <.0001 |

**Supplementary Table 18. Summary statistics for factor quality indices in the LA5c data-driven behavioral bifactor model.** Mean, standard deviation, median, interquartile range, minimum, and maximum values are shown for omega total, omega hierarchical (Omega H), omega hierarchical subscale (Omega HS), explained common variance (ECV), item-level ECV (I-ECV), percentage of uncontaminated correlations (PUC), factor determinacy, and construct replicability (H). Statistics were computed across leave-one-out refits in the pooled LA5c clinical sample.

| Metric | Mean | SD | Median | IQR | Min | Max |
|---|---|---|---|---|---|---|
| Omega total | 0.4354 | 0.08272 | 0.4207 | 0.004842 | 0.4067 | 0.9902 |
| Omega H (general) | 0.09048 | 0.04879 | 0.07952 | 0.004224 | 0.003555 | 0.2749 |
| Omega HS mean (specifics) | 0.08624 | 0.02511 | 0.08531 | 0.001015 | 0.04995 | 0.2467 |
| Omega HS max (specifics) | 0.1608 | 0.122 | 0.1478 | 0.00514 | 0.08538 | 0.9838 |
| ECV (general) | 0.2012 | 0.066 | 0.1887 | 0.00655 | 0.002392 | 0.4294 |
| I-ECV (general) | 0.2789 | 0.07993 | 0.2534 | 0.01193 | 0.2322 | 0.5966 |
| ECV mean (specifics) | 0.1997 | 0.0165 | 0.2028 | 0.001638 | 0.1427 | 0.2494 |
| ECV max (specifics) | 0.3064 | 0.1072 | 0.3019 | 0.006263 | 0.1638 | 0.9947 |
| PUC | 0.6433 | 0 | 0.6433 | 0 | 0.6433 | 0.6433 |
| Determinacy (general) | 0.8437 | 0.02887 | 0.8346 | 0.002265 | 0.83 | 0.9406 |
| Determinacy mean (specifics) | 0.9407 | 0.00438 | 0.9401 | 0.004993 | 0.9291 | 0.9545 |
| Determinacy min (specifics) | 0.8933 | 0.004739 | 0.8938 | 0.003141 | 0.8553 | 0.9024 |
| H (general) | 0.8328 | 0.07192 | 0.7992 | 0.02804 | 0.7785 | 1 |
| H mean (specifics) | 0.8999 | 0.007959 | 0.8987 | 0.0101 | 0.8758 | 0.9256 |
| H min (specifics) | 0.8363 | 0.01275 | 0.8401 | 0.003582 | 0.7533 | 0.8458 |

**Supplementary Table 19. Pairwise correlations among group-level general factor maps in LA5c.** Pearson correlations were computed across the 347 parcels between average general factor maps estimated separately for participants with ADHD, bipolar disorder, schizophrenia, and healthy controls. Higher positive values indicate greater spatial similarity between group-level general factor maps. Diagonal values indicate self-correlations.

| | **ADHD** | **BIPOLAR** | **HC** | **SCHZ** |
|---|---|---|---|---|
| **ADHD** | 1 | 0.888 | 0.891 | 0.827 |
| **BIPOLAR** | 0.888 | 1 | 0.838 | 0.796 |
| **HC** | 0.891 | 0.838 | 1 | 0.881 |
| **SCHZ** | 0.827 | 0.796 | 0.881 | 1 |

**Supplementary Table 20: Diagnostic composition of participants within communities in the LA5c sample.** Percentages and counts of ADHD, bipolar disorder, healthy controls, and schizophrenia are shown for each community, together with total sample size.

| Community | ADHD | BIPOLAR | HC | SCHZ | Total n |
|---|---|---|---|---|---|
| 1 | 13.30% (27) | 17.73% (36) | 50.25% (102) | 18.72% (38) | 203 |
| 2 | 15.35% (33) | 17.21% (37) | 48.37% (104) | 19.07% (41) | 215 |
| 3 | 16.56% (26) | 19.11% (30) | 45.22% (71) | 19.11% (30) | 157 |

**Supplementary Table 21. BrainSMASH and FDR-corrected p-values for parcel-wise correlations between LA5c data-driven centroids, LA5c RDoC classical factor maps, and HCP centroids.** Panel a shows correlations between LA5c DD-specific motifs and LA5c RDoC classical factors (CS, PVS). Panels b and c show correlations between LA5c community centroids and HCP Cohort 1 and Cohort 2 centroids. Panels d and e show correlations between HCP centroids and LA5c RDoC classical factors. P-values were corrected for spatial autocorrelation using BrainSMASH and then adjusted for multiple comparisons using the Benjamini-Hochberg false discovery rate procedure. Corresponding Pearson correlation coefficients are shown in Figure 8g, 8i, and 8j.

**a.) Adjusted FDR-corrected p-values for correlation of DD-specific motifs and RDoC classical factors**

| | CS | PVS |
|---|---|---|
| **C1** | 0.497 | 0.297 |
| **C2** | <.001 | <.001 |
| **C3** | 0.671 | 0.002 |

**b.) Adjusted FDR-corrected p-values for correlation of community centroids from HCP Cohort 1 and DD-specific motifs**

| | C1 | C2 | C3 |
|---|---|---|---|
| **C1** | 0.817 | 0.012 | 0.052 |
| **C2** | 0.817 | 0.024 | 0.391 |
| **C3** | 0.795 | 0.917 | 0.426 |
| **C4** | 0.795 | 0.138 | 0.917 |

**c.) Adjusted FDR-corrected p-values for correlation of community centroids from HCP Cohort 2 and DD-specific motifs**

| | C1 | C2 | C3 |
|---|---|---|---|
| **C1a** | 0.813 | 0.045 | 0.05 |
| **C1b** | 0.787 | 0.482 | 0.787 |
| **C2** | 0.787 | 0.05 | 0.369 |
| **C3** | 0.787 | 0.787 | 0.707 |
| **C4** | 0.787 | 0.24 | 0.841 |

**d.) Adjusted FDR-corrected p-values for correlation of community centroids from HCP Cohort 1 and RDoC classical factors**

| | CS | PVS |
|---|---|---|
| **C1** | 0.008 | 0.993 |
| **C2** | 0.012 | 0.472 |
| **C3** | 0.993 | 0.334 |
| **C4** | 0.323 | 0.993 |

**e.) Adjusted FDR-corrected p-values for correlation of community centroids from HCP Cohort 2 and RDoC classical factors**

| | CS | PVS |
|---|---|---|
| **C1a** | 0.01 | 0.487 |
| **C1b** | 0.433 | 0.413 |
| **C2** | 0.01 | 0.494 |
| **C3** | 0.751 | 0.433 |
| **C4** | 0.433 | 0.751 |

**Supplementary Table 22: Correlations between factor alignment and clinical measures in the LA5c sample.** The first panel shows correlation between alignment of individual specific factor to data-driven community centroids and clinical measures. The second panel shows the corresponding analyses using RDoC classical factor maps for Cognitive Systems (CS) and Positive Valence Systems (PVS). Community based alignment showed significant associations with symptom severity, whereas classical RDoC factor alignment did not. For data-driven community analyses, if an individual contributed more than one specific factor to a given community, alignments were averaged within that community before correlation with clinical measures. For the RDoC comparison, each participant's CS or PVS factor was correlated with the corresponding group average template factor.

**Data Driven:**

| Measure | Community | n | Pearson r | Pearson p | Pearson p BH-FDR |
|---|---|---|---|---|---|
| Hopkins Global Severity | 1 | 203 | -0.1484 | 0.03462 | 0.05193 |
| Hopkins Global Severity | 2 | 215 | -0.2034 | 0.002729 | 0.008187 |
| Hopkins Global Severity | 3 | 157 | -0.06708 | 0.4039 | 0.4039 |
| YMRS score | 1 | 101 | -0.1576 | 0.1155 | 0.1733 |
| YMRS score | 2 | 111 | -0.2262 | 0.01697 | 0.05091 |
| YMRS score | 3 | 86 | -0.02531 | 0.817 | 0.817 |
| ASRS v1.1 Screener | 1 | 203 | -0.1283 | 0.06808 | 0.2043 |
| ASRS v1.1 Screener | 2 | 215 | -0.06008 | 0.3807 | 0.5394 |
| ASRS v1.1 Screener | 3 | 157 | -0.04934 | 0.5394 | 0.5394 |
| Hamilton Depression Rating Scale | 1 | 101 | 0.001259 | 0.99 | 0.99 |
| Hamilton Depression Rating Scale | 2 | 111 | -0.2762 | 0.003349 | 0.01005 |
| Hamilton Depression Rating Scale | 3 | 86 | 0.05208 | 0.6339 | 0.9508 |

**RDoC:**

| Measure | Domain | n | Pearson r | Pearson p | Pearson p BH-FDR |
|---|---|---|---|---|---|
| Hopkins Global Severity | CS | 245 | -0.013 | 0.8395 | 0.8395 |
| YMRS score | CS | 131 | -0.03407 | 0.6992 | 0.8235 |
| Hamilton Depression Rating Scale | CS | 131 | -0.009053 | 0.9183 | 0.9183 |
| ASRS v1.1 Screener | CS | 245 | -0.02668 | 0.6777 | 0.8769 |
| Hopkins Global Severity | PVS | 236 | -0.01698 | 0.7953 | 0.8395 |
| YMRS score | PVS | 125 | -0.02015 | 0.8235 | 0.8235 |
| Hamilton Depression Rating Scale | PVS | 125 | 0.08392 | 0.3521 | 0.7042 |
| ASRS v1.1 Screener | PVS | 236 | -0.01014 | 0.8769 | 0.8769 |

**Supplementary Figure 9: Validation procedure for cross-cohort factor models.** Schematic illustrating the validation approach, where principal components derived from training cohort data-driven factor models were projected onto validation cohorts. Model fit scores were then compared between data-driven factor models and predefined RDoC domain-based models.

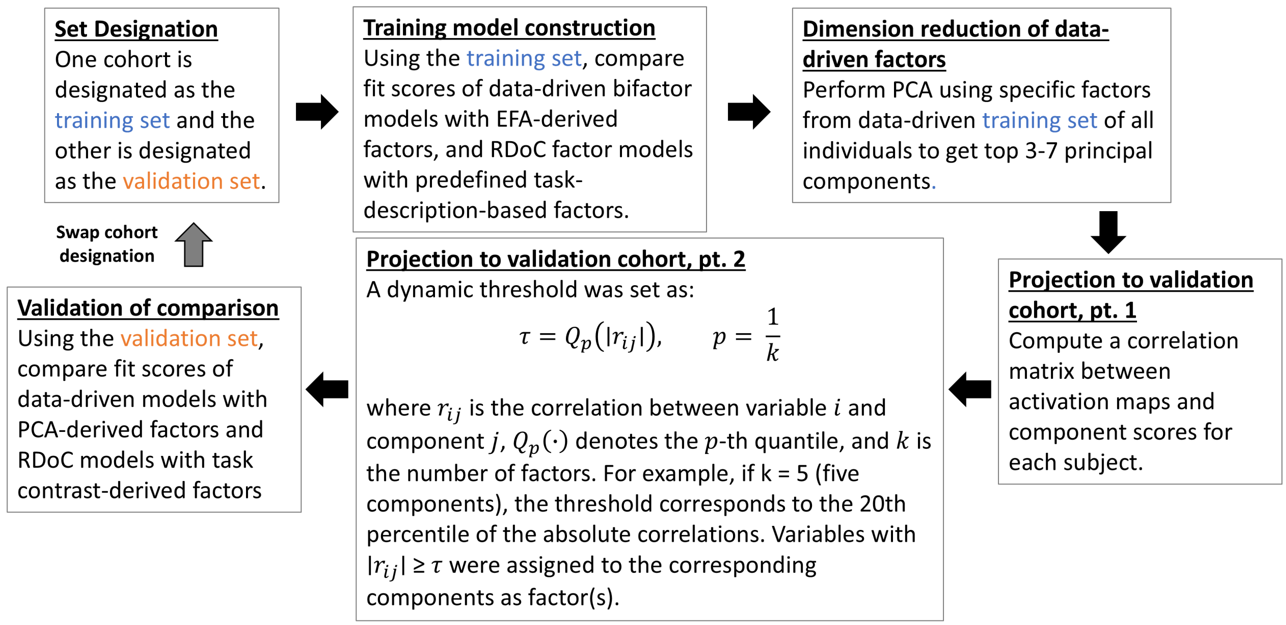

**Supplementary Table 23: Task contrast descriptions and RDoC domain assignments in HCP.** Task contrast codes, brief task descriptions, and their assigned RDoC domains with corresponding domain codes used in RDoC factor model construction.

| Contrast Code | Task Description | Assigned Domain | Domain code |
|---|---|---|---|
| L1 | LANGUAGE: Math problems | Cognitive Systems | CS |
| L2 | LANGUAGE: Story comprehension | Cognitive Systems | CS |
| R1 | RELATIONAL: Shape matching | Cognitive Systems | CS |
| R2 | RELATIONAL: Relational matching | Cognitive Systems | CS |
| S1 | SOCIAL: Random motion perception | Cognitive Systems | CS |
| S2 | SOCIAL: Theory-of-Mind (ToM) | Social Processes | SP |
| E1 | EMOTION: Faces | Negative Valence Systems | NVS |
| E2 | EMOTION: Shapes | Cognitive Systems | CS |
| W1 | WORKING MEMORY: 2-back Body | Cognitive Systems | CS |
| W2 | WORKING MEMORY: 2-back Face | Cognitive Systems | CS |
| W3 | WORKING MEMORY: 2-back Place | Cognitive Systems | CS |
| W4 | WORKING MEMORY: 2-back Tool | Cognitive Systems | CS |
| W5 | WORKING MEMORY: 0-back Body | Cognitive Systems | CS |
| W6 | WORKING MEMORY: 0-back Face | Cognitive Systems | CS |
| W7 | WORKING MEMORY: 0-back Place | Cognitive Systems | CS |
| W8 | WORKING MEMORY: 0-back Tool | Cognitive Systems | CS |
| M1 | MOTOR: Cue | Cognitive Systems | CS |
| M2 | MOTOR: Left Foot | Sensorimotor Systems | SS |
| M3 | MOTOR: Left Hand | Sensorimotor Systems | SS |
| M4 | MOTOR: Right Foot | Sensorimotor Systems | SS |
| M5 | MOTOR: Right Hand | Sensorimotor Systems | SS |
| M6 | MOTOR: Tongue | Sensorimotor Systems | SS |
| G1 | GAMBLING: Punishment | Negative Valence Systems | NVS |
| G2 | GAMBLING: Reward | Positive Valence Systems | PVS |

**Supplementary Table 24: Task contrast descriptions and RDoC domain assignments in LA5c.** Task contrast codes, brief task descriptions, and their assigned RDoC domains with corresponding domain codes used in RDoC factor model construction.

| Contrast Code | Task Description | Assigned Domain | Domain code |
|---|---|---|---|
| bart_1 | BART: Accept | Positive Valence Systems | PVS |
| bart_13 | BART: Reject | Positive Valence Systems | PVS |
| stopsignal_1 | Stop Signal: Go | Cognitive Systems | CS |
| stopsignal_5 | Stop Signal: Successful stop | Cognitive Systems | CS |
| stopsignal_7 | Stop Signal: Unsuccessful stop | Cognitive Systems | CS |
| taskswitch_5 | Task Switching: Congruent switch, short | Cognitive Systems | CS |
| taskswitch_9 | Task Switching: Congruent switch, long | Cognitive Systems | CS |
| taskswitch_13 | Task Switching: Congruent no-switch, short | Cognitive Systems | CS |
| taskswitch_17 | Task Switching: Congruent no-switch, long | Cognitive Systems | CS |
| taskswitch_21 | Task Switching: Incongruent switch, short | Cognitive Systems | CS |
| taskswitch_25 | Task Switching: Incongruent switch, long | Cognitive Systems | CS |
| taskswitch_29 | Task Switching: Incongruent no-switch, short | Cognitive Systems | CS |
| taskswitch_33 | Task Switching: Incongruent no-switch, long | Cognitive Systems | CS |
